# Deep Phenotyping of the Lipidomic Response in COVID and non-COVID Sepsis

**DOI:** 10.1101/2023.06.02.543298

**Authors:** Hu Meng, Arjun Sengupta, Emanuela Ricciotti, Antonijo Mrčela, Divij Mathew, Liudmila L. Mazaleuskaya, Soumita Ghosh, Thomas G. Brooks, Alexandra P. Turner, Alessa Soares Schanoski, Nicholas F. Lahens, Ai Wen Tan, Ashley Woolfork, Greg Grant, Katalin Susztak, Andrew G. Letizia, Stuart C. Sealfon, E. John Wherry, Krzysztof Laudanski, Aalim M. Weljie, Nuala B. Meyer, Garret A. FitzGerald

## Abstract

Lipids may influence cellular penetrance by pathogens and the immune response that they evoke. Here we find a broad based lipidomic storm driven predominantly by secretory (s) phospholipase A_2_ (sPLA_2_) dependent eicosanoid production occurs in patients with sepsis of viral and bacterial origin and relates to disease severity in COVID-19. Elevations in the cyclooxygenase (COX) products of arachidonic acid (AA), PGD_2_ and PGI_2_, and the AA lipoxygenase (LOX) product, 12-HETE, and a reduction in the high abundance lipids, ChoE 18:3, LPC-O-16:0 and PC-O-30:0 exhibit relative specificity for COVID-19 amongst such patients, correlate with the inflammatory response and link to disease severity. Linoleic acid (LA) binds directly to SARS-CoV-2 and both LA and its di-HOME products reflect disease severity in COVID-19. AA and LA metabolites and LPC-O-16:0 linked variably to the immune response. These studies yield prognostic biomarkers and therapeutic targets for patients with sepsis, including COVID-19. An interactive purpose built interactive network analysis tool was developed, allowing the community to interrogate connections across these multiomic data and generate novel hypotheses.

## Introduction

The biological importance of membrane lipids has long been recognized, both as barriers to infection and, once cleaved, as circulating or locally acting modulators of the immune response to pathogens (1–3). While perturbation of the serum lipidome by infections is well-documented (4–8), attention has focused on high abundance lipids such as phospholipids, ceramides and sphingomyelins. This has been true also in COVID-19 where the high abundance lipidome has been implicated in prognosis (9), pathogenesis (10) and disease severity (11–20).

Relatively few reports have focused on eicosanoids, less abundant oxylipins that act locally, but are nonetheless powerful modulators of the immune response (4). Elevated levels of eicosanoids have been reported in plasma (21), tracheal aspirate (22), urine (23) and in stimulated peripheral blood monocytes (24) of patients with severe COVID-19. Snider et al. (25) proposed the eicosanoid generating enzyme, group IIA secretory (s) phospholipase (PL)A_2_, as a marker of disease severity in COVID-19. However, many published studies, including this last, compare the lipid response in patients with that in healthy volunteers, so it is difficult to know the specificity of the response for infection with SARS-CoV-2 compared to other infections.

Both statins (26, 27) and drugs modulating eicosanoid formation or action may be of benefit in preventing or treating COVID-19 (28–30). However, none, to date, including aspirin, has been validated by randomized control trials (31). Despite early concerns, treatment with other non-steroidal anti-inflammatory drugs does not appear to affect adversely the clinical course of COVID-19 (32).

Here, we compare the impact of severe COVID-19 on lipids in plasma, urine and PBMCs with that in both healthy volunteers and patients with sepsis caused by other viruses or bacteria. We integrate this response with clinical parameters, the plasma proteome and peripheral immune cells to reveal immunoregulatory hubs driven by elements of the lipidome that predict disease severity and patient prognosis. This approach defines the relative specificity of these responses. We also addressed the hypothesis that a milder lipidomic signal of inflammation might accompany symptomatic seroconversion in healthy individuals infected with SARS-CoV-2. Finally, we provide a network analysis tool to generate novel mechanistic and therapeutic hypotheses.

## METHODS

### Human Populations and Clinical Data Preparation

Patients were recruited into two research programs at the University of Pennsylvania. All protocols were approved by the Institutional Review Board and were in accordance with the Declaration of Helsinki. Clinical data on the enrolled participants were collected from the electronic medical records (EMRs). Sample sizes were based on availability of biological samples rather than a prespecified effect size.

The first, the COVID-19 Immune Prediction for Lung Recovery study (PICOBS; #IRB843311), was part of the COVID-19 BioBank Collection (IRB#813913) and recruited prospectively from patients admitted to the Penn Presbyterian Medical Center between 04.2020 through 09.2020. All patients were hospitalized and had a positive SARS-CoV-2 test by reverse transcription polymerase chain reaction (RT-PCR). Vulnerable populations, such as children or pregnant women were excluded. After securing consent the patient’s blood and urine was drawn at enrollment and then 2, 7 and 90 days thereafter, according to the patient’s availability in either in- or outpatients’ settings.

The larger research program involved patients recruited to one of 3 prospectively enrolling cohort studies. Hospitalized inpatients admitted either to the Hospital of the University of Pennsylvania or the Penn Presbyterian Medical Center with acute illness due to SARS-CoV-2 were eligible for the Molecular Epidemiology of SepsiS in the ICU (MESSI-COVID, IRB# 808542) study if they were confirmed to have a positive SARS-CoV-2 by RT-PCR and the reason for hospitalization was adjudicated by physician investigators as related to the infection (33–36). Prior to March 2020, subjects were eligible for the same cohort (MESSI) if they were admitted to the medical intensive care unit with sepsis and acute organ failure (35, 37, 38). Most MESSI subjects had confirmed bacterial infection (38). Subjects were excluded if they were admitted to the hospital from a long-term acute care hospital signifying chronic critical illness; if they desired exclusively palliative measures on admission; or if the subject or their proxies were unwilling or unable to provide informed consent prior to discharge or within 3 days of admission. Study personnel screened all hospitalized patients with positive SARS-CoV-2 tests daily, or prior to 2020, screened all patients admitted to the ICU for sepsis and performed informed consent discussions by phone or in person with the subject if he or she were capable of informed consent discussions or with the patient’s proxy by phone if the subject was incapacitated due to illness.

Subjects who provided informed consent had blood drawn at enrollment (time 0) which was within 72 hours of admission and repeated on days 2 and 7. Urine was also collected at these timepoints. Clinical data were abstracted from the EMR into standardized case report forms which included demographic detail, comorbidities, and medications. Clinical laboratory data were obtained from the EMR for the date closest to the time of blood draw. COVID-19 severity was categorized using a scale adapted for clinical trials from the World Health Organization (WHO) ordinal scale as described (37). Severity of illness was assessed using the Acute Physiology and Chronic Health Evaluation III (39) score based on physiologic data at the time of enrollment. Thus, the distinction between moderate and severe cases was based on an ordinal scale, (1, not hospitalized, capable of resuming normal activities; 2, not hospitalized but unable to resume normal activities; 3, hospitalized, not requiring O2 supplementation; 4, hospitalized and requiring O2 therapy; 5, hospitalized and requiring high flow nasal O2 therapy, non-invasive MV, or both; 6, ICU hospitalization, requiring invasive MV or ECMO, or both; 7, death), with scores 1-4 designating mild or moderate cases, and scores 5-7 severe cases. Acute kidney injury was defined according to the Kidney Disease Improving Global Outcomes (KDIGO) creatinine criteria (40) and defining baseline creatinine as the average of outpatient or hospital discharge creatinine values from days 365 days before to 7 days before hospital admission (41,42). If prior creatinine values were missing, baseline creatinine was defined as the lowest value prior to enrollment (42). Survival status was assessed at 30 and 90 days.

Plasma and serum were collected after spinning the tube at 1000×g, 10 min, at 4°C. Aliquot plasma and serum samples were stored at -80 °C. Urine was collected in the morning and aliquots were stored at -80 °C.

Plasma, serum, PBMC and urine samples were also collected from the healthy volunteers (IRB#826459)

Serum samples were also obtained from healthy U.S. Marines who participated in a study conducted by the Icahn School of Medicine at Mount Sinai and the Naval Medical Research Center under a protocol approved by the institutional IRBs (43). The study protocol was approved by the Naval Medical Research Center Institutional Review board (protocol number NMRC.2020.0006) in compliance with all applicable Federal regulations governing the protection of human subjects. The volunteers underwent a 2-week quarantine at home followed by a second supervised 2-week quarantine at a closed college campus that involved mask wearing, social distancing, and daily temperature and symptom monitoring. Study volunteers were tested for SARS-CoV-2 by RT-PCR assay of nares swab specimens obtained between the time of arrival and the second day of supervised quarantine and on days 7 and 14. Serum specimens were obtained at enrollment, the day when participants converted positive by RT-PCR during infection and post infections. Aliquot serum samples were stored at -80 °C. A subset of these serum samples was transferred to the University of Pennsylvania for quantitative measurement of eicosanoids.

## Eicosanoids

### Chemicals and reagents

The following standard compounds and their deuterated analogs were purchased from Cayman Chemicals: Creatinine, d3-Creatinine, PGEM (11α-hydroxy-9,15-dioxo-2,3,4,5-tetranor-prostane-1,20-dioic acid), PGDM (11,15-dioxo-9-hydroxy-2,3,4,5-tetranorprostane-1,20-dioic acid), PGIM (2,3-dinor-6-keto-PGF_1α_), TxM (11deHydro-TxB_2_), 8,12-*iso*-iPF_2α_-VI, iPF_2α_-III, d6-PGEM, d6-PGDM, d3-PGIM, d4-TxM, d11-8,12-*iso*-iPF_2α_-VI, d4-iPF_2α_-III, 14(15)-dihydroxy epoxyeicosatrienoic acid (DHET), 11(12)-DHET, 8(9)-DHET, 5(6)-DHET, 14(15)-EET, 11(12)-EET, 8(9)-EET, 15-hydroxy eicosatetraenoic acid (HETE), 12-HETE, 5-HETE, LTB_4_, LTE_4_, 20-OH-LTB_4_, 13-hydroxy-octadecenoate (HODE), 9-HODE, 12(13)-dihydroxyoctadecenoic acid (DiHOME), 9(10)-DiHOME, d11-14(15)-DHET, d11-11(12)-DHET, d11-8(9)-DHET, d11-14(15)-EET, d11-11(12)-EET, d11-8(9)-EET, d4-LTB_4_, d5-LTE_4_, d8-5-HETE, d8-12-HETE, d8-15-HETE, d4-12(13)-DiHOME, d4-13-HODE, d4-12(13)-EpOME, PGE_2_, PGD_2_, PGF_2α_, 6-keto-PGF_1α_, TxB_2_, linoleic acid (LA), arachidonic acid (AA), eicosapentaenoic acid (EPA), docosahexanoic acid (DHA), lipoxin (Lx)A4, LxB4, resolvin (Rv)D1, RvD2, RvD3, RvE1, RvD5, 18-hydroxyeicosapentaenoic acid (HEPE), d4-PGE_2_, d4-PGD_2_, d4-PGF_2α_, d4-6-keto-PGF_1α_, TxB_2_.

### Sample Collection

Plasma for eicosanoid analysis was separated from whole blood samples by centrifugation at 4°C, snap-frozen, and stored at -80°C. Spot urine samples were also collected within 48 hours of admission and stored at -80°C.

The COVID-19 group included 153 plasma and 176 urine samples drawn from the programs above. The healthy volunteer group included 16 plasma and 26 urine samples, and the non-COVID-19 ICU patient group included 249 plasma and 11 urine samples.

Aliquots of plasma and urine samples were thawed at room temperature just before sample preparation for LC-MS analysis of eicosanoids.

### LC-MS Analysis of Eicosanoid Metabolites in Urine

Two aliquots of 500 μL urine were processed separately. The first aliquot was for the analysis of PGEM, PGDM, PGIM, TxM, 8,12-*iso*-iPF_2α_-VI, and iPF_2α_-III. Stable isotope-labeled internal standards were added to 0.5 ml of urine. The internal standards used were d6-PGEM (25 ng), d6-PGDM (25 ng), d3-PGIM (5 ng), d4-TxM (5 ng), d11-8,12-*iso*-iPF_2α_-VI (5 ng), and d4-iPF_2α_-III (5 ng) in 50 μL of acetonitrile. Methoxyamine (MO) HCl solution (250 μL of 100 g MO HCl solid in 100 mL water) was added, and the sample was allowed to equilibrate for 30 min. The sample solution was brought to a total volume of 1 mL by adding 200 μL of water. The second aliquot was for the measurement of 14(15)-DHET, 11(12)-DHET, 8(9)-DHET, 5(6)-DHET, 14(15)-EET, 11(12)-EET, 8(9)-EET, 15-HETE, 12-HETE, 5-HETE, LTB4, LTE4, 20-OH-LTB4, 13-HODE, 9-HODE, 12(13)-DiHOME, and 9(10)-DiHOME, an internal standard solution containing 5ng of deuterated standards of d11-14(15)-DHET, d11-11(12)-DHET, d11-8(9)-DHET, d11-14(15)-EET, d11-11(12)-EET, d11-8(9)-EET, d4-LTB4, d5-LTE4, d8-5-HETE, d8-12-HETE, d8-15-HETE, d4-12(13)-DiHOME, d4-13-HODE, and d4-12(13)-EpOME in 50 μL acetonitrile was added to 0.5 mL of each urine sample. Two aliquots of urine were purified by solid phase extraction (SPE) using Strata-X 33μm polymeric reversed phase cartridges (Phenomenex, 8B-S100-TAK). The eluate was collected and dried using an Eppendorf vacufuge, and the residue was reconstituted in 100 μL of 50% methanol in water before injection into the HPLC-MS/MS system. Separation was carried out using a Waters ACQUITY UPLC system with a UPLC column, 2.1 x 150 mm with 1.7 μm particles (Waters ACQUITY UPLC BEH C18) and a mobile phase consisting of water (mobile phase A) and acetonitrile:methanol (mobile phase B) with 0.1% formic acid. The flow rate was 350 μL/min, and various linear solvent gradients were used for separation. Quantification was performed using a single point calibration with a standard mix.

To normalize the urinary metabolite concentrations, urinary creatinine was quantified by the same LC-MS system. A stable isotope-labeled internal standard (10 μg/mL d3-creatinine in 3% H₂O/acetonitrile) was added to 10 μL of each urine sample, and the mixture was diluted with 200 μL acetonitrile. Separation was carried out with a UPLC column, 2.1 x 50 mm with 2.5μm particles (Waters XBridge BEH HILIC), and a mobile phase consisting of 100% acetonitrile (mobile phase A) and 5mM ammonium formate water solution (pH = 3.98) (mobile phase B) with a flow rate of 350 μL/min.

### LC-MS Analysis of Eicosanoid Metabolites in Plasma and Serum

For the analysis of eicosanoid metabolites in plasma (including PGE_2_, PGD_2_, PGF_2α_, 6-keto-PGF_1α_, TxB_2_, 8,12-*iso*-iPF_2α_-VI, iPF_2α_-III, 14(15)-DHET, 11(12)-DHET, 8(9)-DHET, 5(6)-DHET, 14(15)-EET, 11(12)-EET, 8(9)-EET, 15-HETE, 12-HETE, 5-HETE, LTB_4_, LTE_4_, 20-OH-LTB_4_, 13-HODE, 9-HODE, 12(13)-DiHOME, 9(10)-DiHOME, LA, AA), a 50 μL sample was mixed with 300 μL of acetonitrile internal standard solution containing 5 ng of deuterated standards for each metabolite (except 1000 ng for d8-AA). Proteins in the plasma or serum sample were then precipitated out with the organic solvent and the sample was centrifuged at 21,000 x g for 2 minutes. The supernatant was transferred to a Phree Phospholipid Removal cartridge (Phenomenex, 8B-S133-TAK) and the eluate was collected and dried in an Eppendorf vacufuge. The resulting residue was reconstituted in 100 μL of 50% methanol in water, transferred to an autosampler vial, and 30 μL of the sample was injected into the HPLC-MS/MS system. The chromatography was carried out using a Waters ACQUITY UPLC system with a UPLC column, 2.1 x 150 mm with 1.7 μm particles (Waters ACQUITY UPLC BEH C18) with a mobile phase A prepared from water and mobile phase B prepared from 95:5 (v/v) acetonitrile:methanol, both containing 0.1% formic acid. The flow rate was 350 μL/min, and separations were carried out with various linear solvent gradients. Quantitation was done by a single point calibration with a standard mix and processed with each batch of samples.

### LC-MS Analysis of Specialized Pro-Resolving Mediators in plasma

We adopted LC-MS methods (44) using multiple reaction monitoring (MRM) to analyze 8 specialized pro-resolving mediators (SPMs) and 2 related fatty acids in plasma samples. To create a standard solution for single point calibration, we prepared a solution of 100 ng/mL of EPA, DHA, LxA4, LxB4, RvD1, RvD2, RvD3, RvE1, RvD5, and 18-HEPE. We followed the same procedures used for eicosanoid measurement in urine, plasma, serum and endobronchial washings to process the standard solution. We used the same internal standards as in the corresponding eicosanoid measurements, with the deuterated lipid compound closest to each SPM peak serving as the internal standard for that compound.

### Endocannabinoid Analysis: Anandamide (AEA) and 2 Arachidonoyl Glycerol (2-AG) in plasma

Endocannabinoids (ECs) were extracted from 50 µL plasma, based on a previously described protocol (45). Briefly, plasma was added to 500µL of chilled methanol/Tris buffer [50mmol/l, pH8] containing internal standards. To this mixture was added 500µL of Tris buffer and 1.5 ml of (2:1) methanol–chloroform. This mixture was centrifuged 500g for 2 minutes, and the chloroform phase was removed to a glass tube. This extraction was repeated twice, and the combined mixture was dried. The dried residue was reconstituted in chloroform. Further, 2 ml of acetone was added to it and the solution was centrifuged at 16,000 x g for 5 minutes. The clear supernatant was collected, dried, and reconstituted in 100 µl methanol. ECs were analyzed by LCMS/MS. MS/MS analysis was performed using Waters Xevo TQ-S instrument equipped with electrospray ionization. Mobile phase A consisted of water/B (95/5) with 0.1% formic acid and mobile phase B consisted of (95/5) acetonitrile and methanol with 0.1% formic acid. A gradient used in the run is as follows: 0 min 30% B; 5 min 50% B; 15 min 100% B; 17 min 100% B; 17.5 min 30% B; 20.5 min 30% B at a flow rate of 0.25 ml/min with a total run time of 20.5 min. Positive electrospray ionization data were acquired using MRM. The mass transitions, collision energy, cone used were: AEA (348.2/287.3, 14, 30), AEA d4 (352.1/287.3, 14, 30), 2-AG:(379.4/287.2, 14, 30); 2-AG d5: (384.4/287.4, 14, 30).

### DESI-MS analysis of high abundance lipids in blood plasma

Metabolites were extracted from blood plasma samples (50 µl) using a modified Bligh-dyer biphasic extraction protocol described elsewhere (46). The nonpolar fraction of the extract was used for DESI-MS based lipidomic analysis. 1 µL samples were spotted onto a PTFE-coated slide and analyzed using high-throughput screening DESI-MS. Slides that were not immediately analyzed were stored at -80°C and warmed up to room temperature for analysis.

DESI-MS experiments were completed using a XevoG2-XS QToF mass spectrometer (Waters, Milford, MA, United States) with a two-dimensional DESI stage source (Prosolia, Zionsville, IN, United States) with optimized parameters. A red sharpie and black Staedtler marker were used to assess calibration. The solvent system consisted of 98% UPLC-MS grade methanol (Fisher Scientific, Watham, MA) and 2% MilliQ water (Milipore Sigma, Burlington, MA) with 50 pg/µL leucine-enkephalin (used as a lock spray reference, Waters, Milford, MA) for negative mode. The same composition was used for positive mode with the addition of 0.1% formic acid (Fisher Scientific) to help with ionization efficiency. Experiments were completed using a flow rate of 2 µL /min with a Harvard syringe pump.

Raw data were exported and analyzed using RStudio (version 4.12 Posit, PBC, Boston, MA) and the MSI.EAGLE package developed in house. This package provides a graphical user interface via R Shiny and uses the Cardinal package (version 2.10) for mass spectral analysis (47). Data were processed using 10% of randomly sampled pixels for peak picking using median absolute deviation (MAD) method. Minimum peak frequency was 0.01. For peak binning, 15 ppm tolerance was allowed. Segmentation UMAP analyses (48) were used to isolate each sample spot. Targeted data analysis was performed by mapping the picked peaks on a recently identified set of serum lipids (49).

### Analysis of Peripheral Blood Mononuclear cells

Peripheral blood mononuclear cells (PBMCs) were collected from 11 PICOBS subjects (6 females and 5 males), aged from 46 to 73 (Supplementary Table 1). Among them, 6 patients were admitted to the ICU. Serum samples were collected from 17 subjects (6 females and 11 males), aged from 19 to 44 (Supplementary Table 2. Among them, 6 patients were admitted to the ICU. The healthy volunteer cohort included 16 participants (8 females and 8 males), aged from 29 to 75 (Supplementary Table 2).

Frozen PBMCs were thawed and resuspended in RPMI 1640 media supplemented with 10% FBS. After centrifugation, the cell pellet was resuspended in RPMI 1640 media supplemented with 2% FBS and split in two aliquots. One aliquot of PBMC was used for flow-cytometry analysis and another aliquot was added in one well of a U-bottom 12-well non-treated plate (Thermo-Fisher) and incubated at 37 °C and 5% CO_2_ for 18 hr. After incubation, the 12-well plate was centrifuged. The cell culture medium was collected for the lipidomic analysis.

#### Flow-cytometry analysis of PBMCs

High-dimensional flow cytometry analysis was performed after thawing frozen PBMCs from COVID-19 patients and healthy controls. The flow-cytometry panel allowed for simultaneous detection of the T-lymphocytes (naïve, cytotoxic, central memory, effector memory (EM), and EMRA phenotypes of CD8+ T-cells and CD4+ T cells, exhausted like CD8+ T-cells; CD4+ follicular helper T cells); B-lymphocytes; monocytes (classical and non-classical); natural killer cells, and dendritic cells populations as previously described (35). Cells were stained for viability exclusion using Live/Dead Ghost Dye in PBS for 10 minutes with Fc Block at room temperature (RT). Cells were then washed and stained for 30 min at RT with the surface antibody master mix containing antibodies (Supplementary Table 3) in fluorescence-activated cell sorting (FACS) buffer (PBS containing 2% fetal bovine serum) and Brilliant stain buffer (BD Biosciences, San Jose, CA). After RT incubation, cells were washed and fixed with 1% PFA prior to data acquisition on a BD LSR cytometer II (BD Biosciences). Data were analyzed using FlowJo software (version 10.6.2, Tree Star, Ashland, OR).

Data and materials availability: All compensated flow cytometry files are publicly available at https://hpap.pmacs.upenn.edu/explore/download?otherdonor

#### Lipidomic analysis of PBMC cell culture samples

Oxylipins were measured by liquid chromatography-mass spectrometry as described above. Briefly, 800ul of the PBMCs medium was used for solid phase extraction (SPE). A cocktail of stable isotope-labeled internal standards was added to each sample in 50 μl of acetonitrile prior to SPE. SPE was conducted using Strata X, 30mg, 1 mL tubes (Phenomenex). The cartridge was conditioned with 1ml acetonitrile followed by 0.25 ml water. Samples were loaded to the cartridge followed by washing with 1ml solvent constituting 5% (methanol/water). The cartridges were dried by vacuum for 15 mins and samples were eluted with 0.1% formic acid in 1 ml methanol. The extracts were dried under a stream of nitrogen and reconstituted in 120µL solvent constituting 50:50 (v/v) methanol and water. The samples were transferred to the autosampler tube, 30µL of samples were injected to ultra-performance liquid chromatography tandem mass spectrometry (UPLC-MS/MS). A Waters Acquity UPLC BEH column (2.1 x 150mm) was used for the separation of the compounds. Mobile phase A consisted of H2O/B (95/5) + 0.1% Formic acid. Mobile phase B consisted of Acetonitrile/Methanol (95/5) + 0.1% Formic acid. Following gradient was run: 0 min 20% B; 5 min 2% B; 6 min 10% B; 12 min 60% B; 15 min 2% B; at a flow rate of 0.35 mL min−1. A Waters Xevo TQ-S instrument was run in negative ionization mode for detection of the compounds.

### Measurement of Phospholipases in plasma

Plasma Group 11A secretory phospholipaseA_2_ (sPLA_2_-IIA) concentrations were determined by ELISA (Cayman Chemical). Plasma samples were diluted (1:40–1:2560) and were subjected to the instructions provided by the manufacturer. Concentrations of sPLA2 -IIA in plasma were calculated using standard curves.

Plasma Group 11D secretory phospholipase A_2_ (PLA_2_G2D) was also measured in plasma by ELISA (My Biosource). Plasma samples were diluted fourfold using PBS (pH=7.0) and assayed as per manufacturer’s instruction. Concentrations were calculated using standard curves.

Plasma cytosolic (c)PLA_2_ was measured as part of the O-link proteomic analysis, a relative quantification from Ct values.

### Proteomic Analysis in plasma

Proteomic analysis was performed using the O-link platform, as previously described (35,50). Briefly, whole blood was spun within 2 hours of blood collection (3,000 rpm, 15 min), and plasma was collected, aliquoted, and frozen at –80°C until assay. Plasma was not immunodepleted. We used the O-link Proximity Extension Assay to measure 713 unique proteins. In this assay, oligonucleotide-labeled monoclonal or polyclonal antibodies are used to bind each protein target in a pairwise manner upon which the paired oligonucleotides hybridize. The unique hybridization product is amplified by PCR, and multiplex detection occurs in a high throughput fluidic chip system. On each plate, a common interplate control of pooled “healthy” plasma acquired and processed at O-link facilities was used for normalization resulting in a semiquantitative measurement for each protein on log2-transformed scale referred to as the normalized protein expression (NPX).

### Statistical analysis

High abundance lipid data were analyzed in Rstudio running on R 4.1.2. Non-parametric statistical analyses (Kruskal Wallis and Mann Whitney tests) was performed using base R stats package and in house scripts.

Statistical significance was reported after multiple testing correction using Benjamini Hochberg method. All plots were generated using ggplot2 (v3.4.1), pheatmap (v1.0.12) packages and in house scripts. High abundance lipid data was analyzed in Rstudio running on R 4.1.2. Non-parametric statistical analysis (Kruskal Wallis and Mann Whitney tests) was performed using base R stats package and in house scripts. All plots were generated using ggplot2, pheatmap packages and in house scripts. Multivariate analysis was performed using Simca-P 17 (Startorius Stedim, Bohemia, NY).

To evaluate the relationship between eicosanoid concentrations and both the disease severity score and the frequency of immune cell subtypes, we computed the Spearman correlation coefficient based on the rank of these variations. We used a t-distribution with n-2 degrees of freedom, where n is the sample size, to calculate the associated p-value for the Spearman correlation coefficient. To assess differences in eicosanoid concentrations between different patient groups and the healthy control group, we used the Kruskal-Wallis test, calculated using Prism 9 software.

To assess differences in serum LA metabolite concentrations between at different time points (before infection, during infection, and after infection) in the US Marines, we used a one-way analysis of variance (ANOVA) with subsequent pairwise comparisons as appropriate, all calculated using Prism 9 software.

### Integrative correlation analysis

The data used in the integration analysis were obtained from non-COVID ICU patients with sepsis and COVID-19 patients as described above. Non-COVID patients were more critically ill than COVID-19 patients, as reflected by their APACHE scores.

The data comprise measurements of clinical features, proteins, metabolites, and cell populations as described above. Table 1 summarizes the numbers of various features used in the integrative analysis. Not all measurements were available across all samples. The sample sizes for several cohorts of interest are shown in Supplementary Table 4.

**Table 1:** Sizes of feature sets used in integrative analysis. Plasma protein features include sPLA_2_ and PLA_2_G2D. Some proteins were measured in multiple Olink panels, and such proteins are represented by two or more, usually highly correlated, features.

|  | Number of features used in the integrative analysis |
| --- | --- |
| Plasma proteins | 730 |
| Plasma eicosanoids | 18 |
| Urine eicosanoids | 18 |
| Positive mode plasma lipids | 393 |
| Negative mode plasma lipids | 205 |
| Flow cytometry data | 216 |
| Mass cytometry data | 71 |
| Endocannabinoids | 2 |
| Clinical and demographic data | 100 |

Four correlation matrices were obtained by estimating Spearman’s correlation coefficients using available measurements from all four cohorts of interest: COVID-19, non-COVID ICU, moderate COVID-19, and severe COVID-19.

The correlation matrix for the COVID-19 cohort was transformed into a distance matrix by substituting each correlation coefficient estimate ρ with 1−ρ^2^. This distance matrix was provided as an input to UMAP to visualize the correlation network of measured features. Certain pairs of features had insufficient sample size to calculate an estimate of their correlation. The distance between such feature pairs was set to 1 (maximum), and consequently, they were not included as edges of the target simplicial set, making UMAP rely on other feature pairs, for which correlation estimates were feasible, when positioning features in the plane.

To visualize and explore the differences between the correlation network of features in COVID-19 and non-COVID ICU cohorts, we calculated the differential correlation matrix using the formula ½ (ρ_COVID-19_ − ρ_non-COVID_) and provided the corresponding distance matrix to UMAP. We also constructed an undirected and unweighted differential correlation graph in which nodes correspond to measured features and the existence of an edge between a pair of features indicates significant (p < 0.05) change between correlation coefficients ρ_non-COVID_ and ρ_COVID-19_ for the pair of features. The p-values for differential correlation were based on statistics (Z_non-COVID_ – Z_COVID-19_)/((N_non-COVID_ – 3)_–1_ – (N_COVID-19_ – 3)_–1_)_1/2_, where Z_non-COVID_ and Z_COVID-19_ denote the Fisher’s Z-transformation of coefficients ρ_non-COVID_ and ρ_COVID-19_ respectively. Features with highest degree centrality and betweenness centrality exhibit the largest differences in how they correlate with other features in COVID-19 vs. non-COVID-19 cohorts and these were singled out. A similar analysis was performed to compare moderate COVID-19 and severe COVID-19 correlation structures, but it did not produce as many significant results, most likely due to reduced sample sizes (Supplementary Table 4). Differences were apparent in individual data sets (e.g., urinary eicosanoids) where sample sizes were greater.

The data are presented as an interactive feature browser for further hypothesis generation at covid.itmat.org.

The analysis was performed using Python 3.10.9 with additional packages SciPy (version 1.10.0), UMAP-learn (version 0.5.3) and NetworkX (version 2.8.8). STRING (58) version 11.5 was used for enrichment analyses.

## RESULTS

### 1. Phospholipases

sPLA2-IIA concentrations were measured in plasma from 181 subjects. Supplementary Fig 1A shows the distribution of sPLA_2_-IIA in the healthy volunteer, ICU non-COVID-19 and ICU COVID-19 groups. The median (interquartile range) concentration of sPLA_2_-IIA is elevated significantly in COVID-19 patients with median values of 53.9 ng/ml (23.7 ng/ml, 173.5ng/ml; p < 0.0001) and also in non-COVID patients with sepsis with median values of 123.3 ng/ml (36.06 ng/ml, 356.47 ng/ml; p < 0.0001) with respect to control subjects 9.8 ng/ml (7.4ng/ml, 22.2ng/ml).

sPLA_2_-IIA levels reflected disease severity in COVID-19 patients and was positively correlated with the associated ordinal score 7 days after admission (Spearman’s ρ = 0.31, p = 0.01) while also correlating with other indices of disease severity such as platelet to lymphocyte ratio (PLR), neutrophil to lymphocytet ratio (NLR), and C reactive protein (CRP), all of which differed significantly between patients with severe and moderate COVID-19 (Spearman’s ρ = 0.38, 0.38, 0.67, p<0.01). These results are consistent with the suggestion that sPLA_2_ reflects severity in COVID-19 (25).

sPLA_2_ was positively correlated with diverse biochemical measures which reflected and/or predicted disease severity (Supplementary Table 5). These included urinary TxM, LPC-O-16:0 (see below) and the proteins such as the interleukin 13 receptor subunit alpha 1, disulfide isomerase (P4HB) and E3 Ubiquitin protein ligase, indices of lymphocyte and monocyte chemotaxis, such as MCP-1, MCP-3, CXCL10, IL-6, and S100A12 and immune markers of CD4+ and CD8+ T cell activation like HLADR+ and CD38+ in the central memory CD4+ T cells. These observations implicate sPLA_2_ as the likely source of the eicosanoid storm. Positioning of sPLA_2_ within the inflammatory connectome is discussed further below.

In our cohort, plasma concentrations of cPLA_2_ did not discriminate between the COVID-19 vs non COVID-19 sepsis patients, while PLA_2_G2D was lower in the COVID-19 patients than in both the non-COVID sepsis group and the healthy controls (Supplementary Fig 1b, c). Plasma concentrations of PGE_2_ and 5(6)-DHET were correlated with cPLA_2_, while plasma 12-HETE was correlated with PLA_2_G2D.

Interestingly, PLA_2_G2D (negatively) and sPLA_2_ (positively) correlated with blood glucose – the latter relationship as previously described (52). PLA_2_G2D was also positively correlated with expression of proteins relevant to lipid metabolism (Perilipin)-, cellular adhesion (myocillin) and inflammation (C-type lectin domain family 7 member A: CLEC7A) and previously characterized using the O-link platform in a subset of the MESSI cohort (35).

In summary, sPLA_2_ relates to disease severity in COVID – 19 but is also elevated in patients with other forms of sepsis in the ICU. It serves as a source of eicosanoid generation in patients with sepsis and these downstream products themselves reflect and/or predict the course of the disease and correlate with distinct immune features and proteins as described below. This is consistent with their importance in shaping the immune response and clinical outcome in COVID-19.

### 2. Eicosanoids

#### (i)#Urine and Plasma eicosanoids in ICU patients with sepsis compared to healthy controls

The normalized differences of average concentrations of measured metabolites between patient groups and healthy controls are shown in Figure 1 A,B. Differences in concentrations between patient groups and the heathy volunteer group were normalized by dividing by the standard deviation of healthy controls (σHC). Urinary PGEM, PGDM, PGIM and TxM were all significantly elevated in both severe COVID-19 patients and in non-COVID ICU patients when compared to controls (Fig 1A). This was also true of urinary 8, (9)-DHET and the LA metabolites, 9,10- and 12,13-Di-HOMEs. Consistent with these observations, the concentrations of the substrates, AA and LA in plasma, were also significantly higher in the patients in ICU than in healthy controls, irrespective of their COVID status (Fig 1B). While the relative elevations of these compounds usually reflected the more severe disease evident in non-COVID-19 vs COVID-19 patients (e.g., PGEM, 5- and 15-HETE, DHETs and DiHOMEs), the elevations in urinary PGDM, PGIM and 12-HETE were more pronounced in COVID patients, despite having less severe disease.

**Figure 1:**
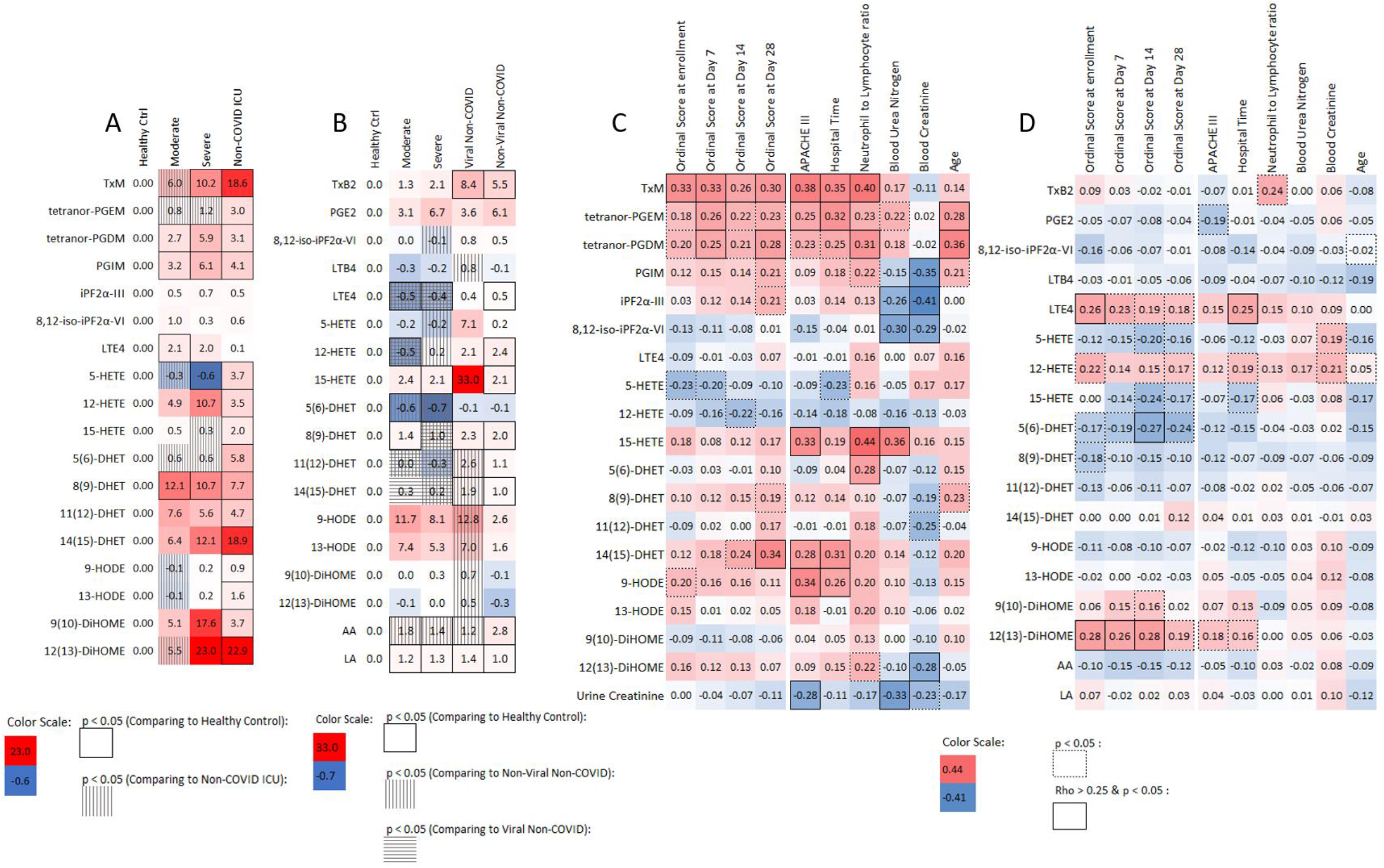
Eicosanoids and Disease Severity. Comparison of eicosanoid profiles and Spearman correlation with severity scores and organ damage indicators between COVID-19 patients, non-COVID patients, and healthy controls. Heat maps show the normalized average levels of eicosanoids and linoleic acid metabolites in A) urine and B) plasma, with values in each box representing the difference in concentration between the patient group and healthy controls, divided by the standard deviation of the healthy controls. P values were obtained by performing a two-tailed, equal variance Student’s t-test. The heat maps highlight significant differences in eicosanoid profiles between patient groups and healthy controls. Additionally, the Spearman correlation coefficient (ρ) between eicosanoids and different severity scores and organ damage indicators in C) urine and D) plasma are presented. The color scale represents the strength and direction of the correlation. Warmer colors indicate positive correlations and cooler colors indicate negative correlations. The p-value was calculated and marked in the heat maps.

In plasma, a similar pattern of eicosanoids reflecting disease severity, rather than origin in virally infected patients was apparent (Tx, PGE_2_) while in some instances (5- and 15-HETE and 9- and 13-HODEs), the elevations in non-COVID patients whose disease was viral were more pronounced than in those with bacterial disease despite similar clinical severity.

We tested a subset of urine, plasma, serum samples and endobronchial washings but did not observe peaks for LxA4, LxB4, RvD1, RvD2, RvD3, RvE1, or RvD5.

In conclusion, there is an eicosanoid storm evident in patients admitted to the ICU with sepsis caused by COVID-19 or other pathogens. While there is no signature uniquely distinct for COVID, some products (urinary PGDM, PGIM, 12-HETE) appear to be disproportionally elevated in COVID compared to other patients with sepsis despite their greater disease severity, while others (5- and 15-HETE and 9- and 13-HODEs) appear to be more elevated in patients with severe viral rather than bacterial origins to their disease.

#### (ii) Eicosanoids as predictors of disease severity

Next, we calculated the Spearman correlation between the eicosanoids and different clinical severity scores and indices of organ damage. As expected, metabolites of major pro-inflammatory eicosanoids, TxM, PGEM and PGDM in urine reflected disease severity correlating with the COVID-19 ordinal scale described above (Fig. 1C). Less pronounced relationships were evident for 14(15)-DHET, 9- and 13-HODEs, the isoprostanes and 12(13)-DiHOME. Similarly, in plasma LTE_4_, 12-HETE, and DiHOMEs were significantly correlated with NIH ordinal scale (Fig. 1D). Further analysis (Fig. 2) showed that levels of urinary TxM, PGEM, and PGDM and plasma LTE_4_, 12-HETE and DiHOMEs increased with disease severity. Thus, more severely ill patients had a more intense inflammatory response and higher levels of metabolites from the COX pathway in urine. Although the levels of LTE_4_, 12-HETE, and DiHOMEs in plasma were not significantly higher than those in healthy controls, these metabolites also increased with disease severity. These lipids were also correlated with other indicators of disease severity, such as the APACHE III scores, and time in hospital (Fig. 1C). A well-accepted index of COVID-19 severity, the NLR correlated positively with urinary TxM, PGEM and PGDM and with LTE_4_ in plasma. Urinary PGEM and PGDM were also positively correlated with age, another marker of disease severity in COVID-19.

**Figure 2.**
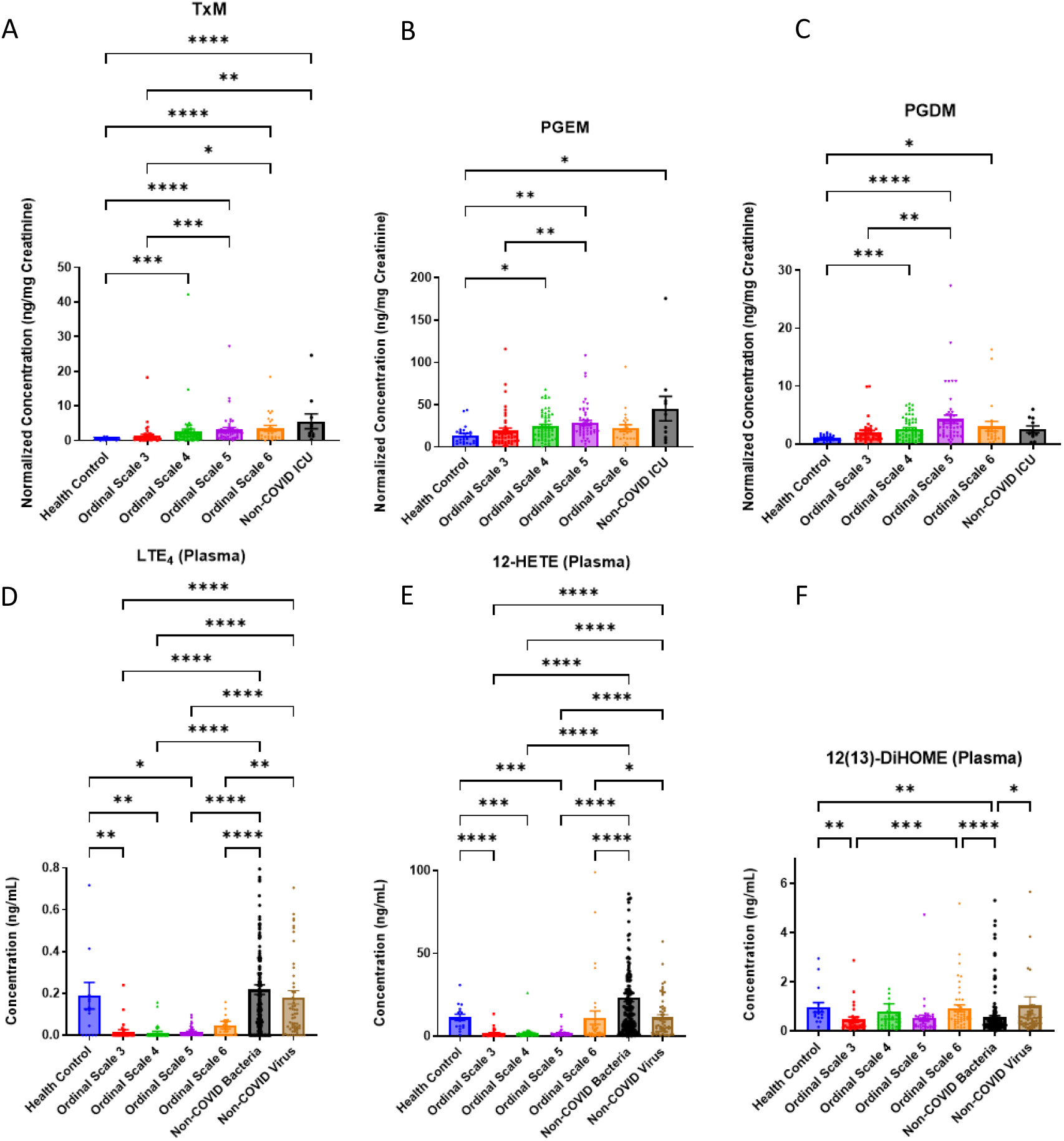
Selected eicosanoids in each severity group. The figure shows the normalized concentrations of urinary TxM(A), PGEM(B), PGDM(C) and concentrations of plasma LTE_4_(D), 12-HETE(E), 12(13)-DiHOME(F) in each severity group, with error bars indicating standard error of the mean. The statistical significance was determined using Kruskal-Wallis test and marked as follows: * P<0.05, ** P<0.01, *** P<0.001, **** P<0.0001.

Pro-inflammatory eicosanoids – reflected by TxM, PGEM, PGDM, PGIM, the isoprostane, iPF_2α_-III, and 14(15)-DHET in urine and LTE_4_, 12-HETE, 12(13)-DiHOME in plasma - remained correlated with the ordinal scale of disease severity 28 days after admission (Fig. 1). Thus, elevation of these pro-inflammatory lipids at admission associated with persistent pneumonia severity at 28d.

The endocannabinoids, AEA and 2-AG were elevated in COVID-19 (Supplementary Figure 1D,E). 2AG was significantly correlated with a history of pulmonary disease and disease severity in sepsis patients. It was also correlated with other plasma lipids linked to disease severity (TxM, PGE_2_, LTB_4_ and LPC-O-16:0). Similarly AEA correlated with hospital mortality and 12(13)-DiHOME.

Bacterial infection triggered a broader eicosanoid response than observed in viral sepsis (Suppl Fig. 2). In both viral sepsis cohorts, lipid species belonging to the same pathway tended to correlate positively amongst themselves and showed no significant associations with those from different pathways (Suppl Fig. 2A). In contrast, bacterial sepsis resulted in widespread positive cross correlations amongst products of different eicosanoid pathways (Suppl Fig. 2B). Here also, eicosanoids strongly correlated with disease severity. Thus, 15-HETE, 11(12)-DHET and 14(15)-DHET positively correlated with the APACHE III disease severity score, while PGE_2_, Tx and 12-HETE correlated with the NLR and monocyte-to-lymphocyte ratio (MLR) (Suppl Fig. 2C).

In some cases, the deterioration of renal function - BUN and plasma creatinine correlated with COVID-19 severity – may have limited clearance of proinflammatory markers into urine and undermined their relationship to disease severity. For example, 8,12-*iso*-iPF_2α_-VI, 5-HETE, and 12-HETE in urine were negatively correlated with BUN, creatinine and disease severity.

#### (iii) Eicosanoid response to COVID – 19 seroconversion in healthy Marines

Here we addressed the hypothesis that healthy individuals undergoing seroconversion after infection with COVID-19 might express a milder lipidomic signature of inflammation compared to that observed in patients with sepsis and that this might be reflected by mild inflammatory symptomatology. Having segregated patients based on symptomatology we were then surprised to find that LA increased significantly after infection only in those who seroconverted asymptomatically but not in the symptomatic group (Supplementary Figure 3A). Consistently, its 9(10)-DiHOME and 11(12)-DiHOME metabolites also increased only in the asymptomatic group and continued to increase even after patients tested positive. (Supplementary Figure 3B, C) Another two LA metabolites, 9-HODE and 13-HODE followed the same trend as the DiHOMEs, but the changes were not statistically significant (Supplementary Figure 3D, E). Additionally, LA and 5(6)-DHET were positively correlated with cycle threshold (CT) values of the qPCR test, indicating that concentrations of these two compounds were negatively correlated with viral load.

#### (iv) Eicosanoids and immune cells

Although lymphocytes have a low capacity to generate them, eicosanoids are potent modulators of the immune response to viral infection (4). The Spearman correlations (ρ) between immune cell types were significantly correlated with COVID-19 severity (q-Value<0.05) and eicosanoids (Fig. 3). Urinary TxM, PGEM, PGDM, and 15-HETE correlated with CD4 T cells expressing CD38 and HLA-DR markers of CD4+ T cells activation. Similar to urinary PGEM and TxM, urinary PGDM was correlated with CD4+ T cell activation. Its relationship with the frequency of non-T non-B cells was more pronounced than that of either PGEM or TxM. Urinary 15-HETE was correlated with CD8 T cells expressing CD38, HLA-DR and KI67. It was also positively correlated with plasmablasts and negatively correlated with B cells non-plasmablast.

**Figure 3.**
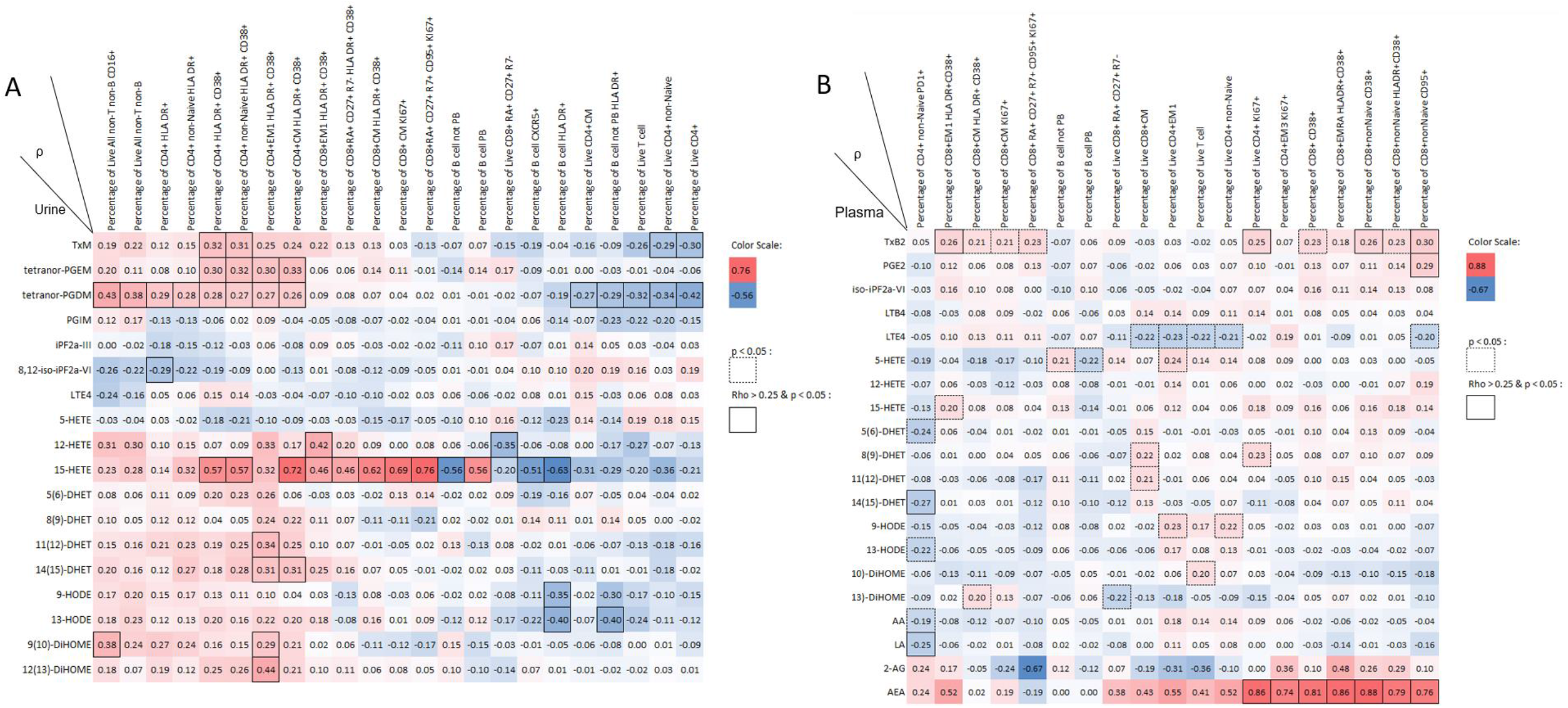
Correlation between eicosanoid concentration and the frequency of immune cell subtypes. Heat maps show the Spearman correlation coefficient (ρ) between eicosanoid concentrations and the frequency of immune cell subtypes in A) urine and B) plasma. The color scale represents the strength and direction of the correlation, with warmer colors indicating positive correlations and cooler colors indicating negative correlations. The p-value was calculated and marked in the heat maps.

The endocannabinoid AEA was correlated with activation and proliferation of T cells. For example, it correlated with activation of CD8 T cells, with the activated CD8 EMRA subset, with proliferating CD4 T cells and with expression of apoptotic markers such as Fas on non-naïve CD8 T cells (Fig.3B).

In summary, these results are consistent with eicosanoids – specifically, PGE_2_ TxA_2_ and LTE_4_ and to a lesser extent, PGD_2_ promoting the activation of CD4+ T cells in patients with severe COVID-19. PGD_2_, an eicosanoid that exhibits some selectivity for sepsis due to COVID-19, appears to exhibit a particular influence on non-B non-T cells and CD4+ T cells while the endocannabinoid, AEA is linked to CD8 T cell activation.

### PBMCs

The relative differences of average concentrations, corrected by cell number, of measured metabolites between COVID-19 patients and healthy controls are shown in Fig. 4A. PBMC PGE_2_ was significantly elevated in COVID-19 patients, consistent with our results above in urine and plasma and with other studies (21,47). In addition, PBMC release of PGF_2α_ and AA were also modestly but significantly increased in the COVID-19 group compared to the healthy controls.

**Figure 4.**
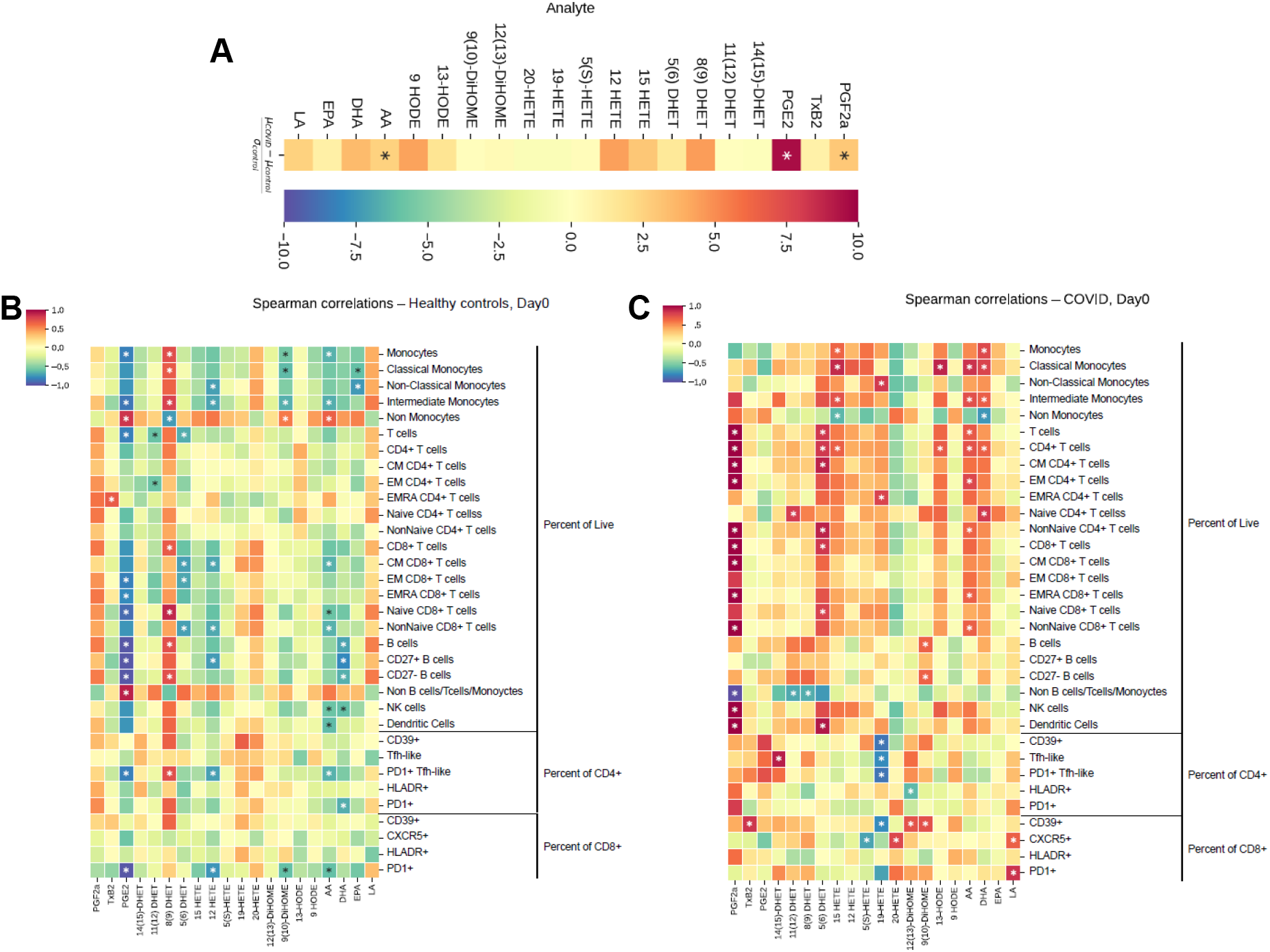
(A) Changes in the lipidomic profile of PBMC supernatant from COVID-19 patients compared to healthy volunteers. Heat map shows differences of normalized average levels of arachidonic acid and linoleic acid metabolites, corrected by cell number. P values were obtained by performing Mann-Whitney U-test. (B-C) **Correlation between eicosanoid concentrations and frequency of immune cell subtypes in PBMCs.** Heat map shows Spearman’s ρ of correlation between lipid metabolites and frequency of immune cell subtypes in PBMCs from healthy controls (B) and COVID-19 patients (C). p value was calculated and marked in the heat maps. * indicates P<0.05. The color scale reports the Spearman’s correlation coefficient ρ. Warmer colors indicate positive correlations, while cooler colors indicate negative correlations.

### Correlation between lipid profile and immune features

The Spearman ρ of the correlations between cell types and lipid metabolites are depicted in Fig. 4 B-C. As expected, there was a striking difference in the relationships observed between the two groups. In PBMCs from healthy controls, PGE_2_ was negatively correlated with monocytes, B cells and some subpopulations of memory T cells; 8,(9)-DHET was positively correlated with classical monocytes, CD8+ T-cells, follicular T helper cells and B-cells, while it was negatively correlated with non-monocytes (Fig. 4B). In contrast, in PBMCs from COVID-19 patients, PGF_2α_ was positively correlated with different sub-populations of memory CD4+ and CD8+ T-cells, dendritic and NK cells. AA and 5(6)-DHET were positively correlated with subpopulation of CD4+ T cells. TxB_2_ and, as previously described, 9(10)-DiHOME and 12(13)-DiHOME (52) were positively correlated with CD39+ CD8+T-cells (53). LA was positively correlated with PD1 and CXCR5 expression in CD4+ and CD8+ T cells, suggesting a shared module in T cells associated with LA. 15-HETE, AA, DHA and 13-HODE were positively correlated with classical monocytes. 19-HETE was positively correlated with non-classical monocytes, while it was negatively correlated with Tfh cells and with CD39+ expression on CD4+ and CD8+ T-cells (Fig. 4C).

In summary, PGE_2_ and PGF_2α_ production are elevated in PBMCs from patients with COVID-19. PGF_2α_ release correlates with broad populations of immune cells in PBMCs from patients with COVID-19.

### 3. High abundance lipids

#### Ether phospholipids and ChoE-18:3 are related to severity of COVID-19

High abundance lipidomics data were obtained from the plasma of 67 COVID-19 patients, 66 non-COVID patients and 16 age-matched healthy controls at the day of enrolment (day 0) and after a week (day 7). Positive (1139 features) and negative (985 features) ionization mode data were analyzed to obtain untargeted lipid data. Supervised orthogonal partial least square discriminant analysis (OPLS-DA) model from both datasets yielded distinct clustering of the subject groups (Supplementary Figure 4). The subject groups showed similar clustering at both time-points. We sought to determine if the lipidomic response differed between the two time-points using a shared and unique structure (SUS) analysis of the OPLS-DA models of respective days (Supplementary Figure 5). This analysis demonstrates that the differential lipid responses in positive mode lipids across the three classes are similar between the two timepoints, suggesting a week of treatment/ICU intervention was not enough to induce discernible changes in this subset of the lipidome.

Interestingly, several negative mode lipid features showed opposing trends between day 0 and day 7. We analyzed the day 0 samples further and used a Kruskal Wallis one-way ANOVA to identify specific features that are differentially expressed across the three groups of subjects. A large fraction (∼42% and ∼26% for positive and negative modes, respectively) of the data was found to show significant variance (false discovery rate [FDR] < 0.2). However, major clusters of variance were associated with an increase/decrease of features in non-COVID subjects, highlighting the importance of considering non-COVID infected patients as one group of controls. We did observe, however, small but distinct clusters of lipids with COVID-19 specific variation in both datasets (Supplementary Figure 6).

We next mapped our untargeted feature sets onto known literature reported lipids by mass/charge (m/z) values (55,56) and created targeted lipid datasets. We assigned 393 and 205 lipids from positive and negative mode datasets, respectively (Supplementary Table 6). We first used OPLS-DA analysis to identify multivariate clustering between the three groups of subjects. Both positive and negative mode data yielded significant models (Q2(cum) = 0.45, 0.26 for positive and negative mode data, respectively; CV-ANOVA p < 0.0001 for both) and showed significant clustering between the three groups (Figure 5A, B). Using a Kruskal Wallis test, we found 15 positive mode lipids that were significantly altered in COVID-19 patients (FDR < 0.05, pairwise p values –COVID-19 vs non-COVID-19 sepsis patients < 0.05, COVID-19 vs healthy control < 0.05, healthy control vs non-COVID-19 sepsis patients > 0.05, Figure 5C, Supplementary Figure 7). Using the same FDR and p-value cutoffs for the negative mode data, we found six features belonging to five lipids that were specifically altered in COVID subjects (Figure 5C, Supplementary Figure 8, two features were annotated as FA 18:3). Sphingomyelin levels were elevated in COVID-19 patients with sepsis compared to both non-COVID-19 sepsis patients and healthy controls, in agreement with previous observations (18), while the majority of phospholipids (phosphatidylcholines, phosphatidylinositols, ether phospholipids) were decreased. Interestingly, cholesteryl ester 18:3 (ChoE-18:3) was decreased in COVID-19 patients while the corresponding fatty acid was elevated. We then sought to determine which of these lipid signatures was also a function of COVID-19 severity by mapping ordinal scores to subjects as severe (ordinal score 5-7) and moderate (< 1-4) groups and compared the lipid levels using nonparametric tests. Three lipids, ChoE-18:3, LPC-O-16:0 and PC-O-30:0 significantly varied between the moderate and severe COVID-19 groups (FDR < 0.2, Figure 5D).

**Figure 5:**
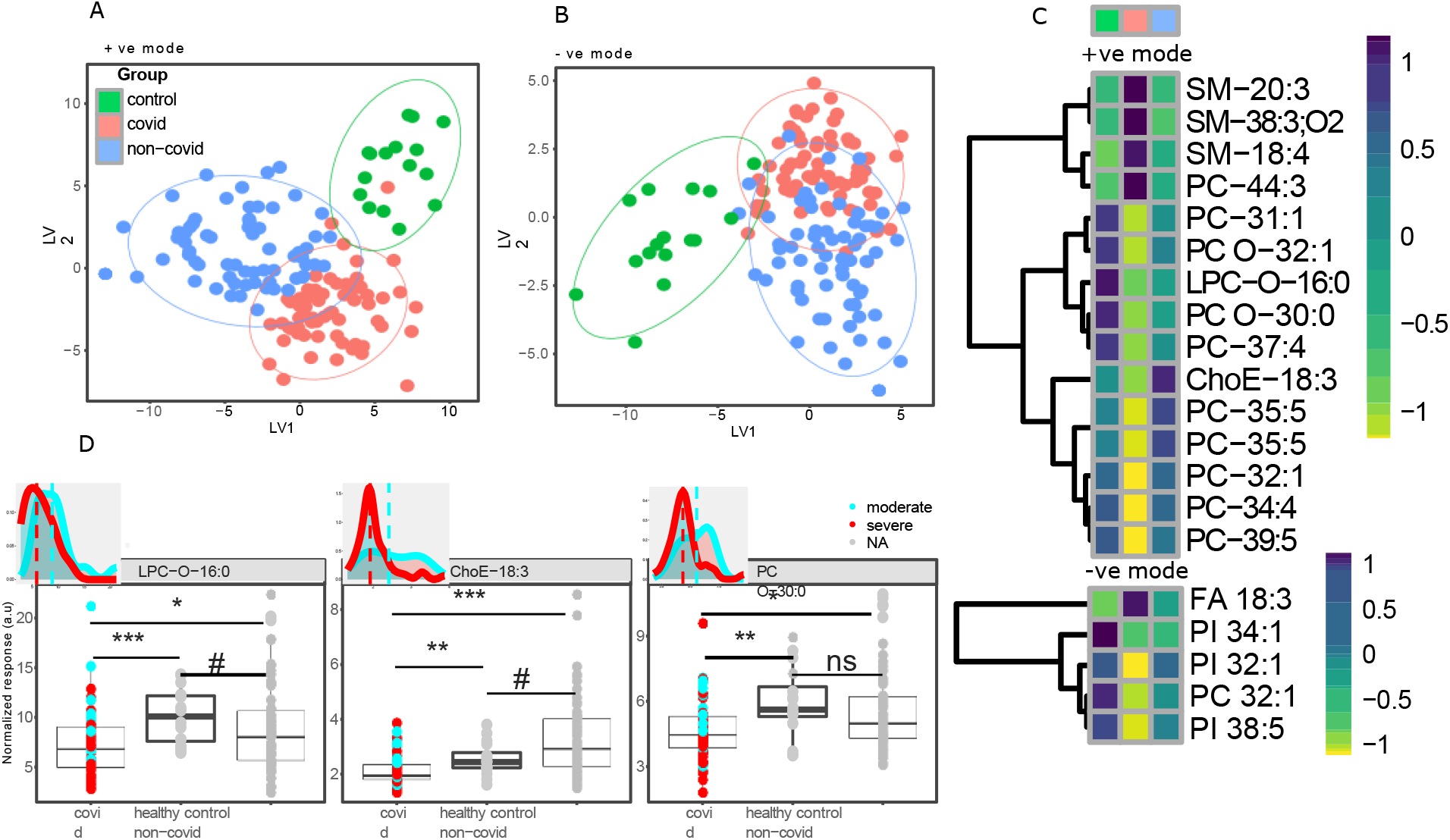
High abundance lipidomic response to Covid-19 infection. Targeted positive and negative mode lipidomics data was assessed to evaluate signatures of Covid-19 specific alterations. OPLS-DA scores plot from A) positive mode and B) negative mode lipid data. C) Lipids specific to Covid-19 assessed using Kruskal Wallis test and pairwise Mann-Whitney test (Kruskal Wallis FDR < 0.2, p(Mann Whitney) < 0.05 between covid/non-covid and covid/control; > 0.05 between non-covid/control. Median value of each lipid for each group is presented as heatmap. D) Boxplots showing three lipids that are covid specific as well as significantly different (FDR < 0.2) between moderate (ordinal score > 4) and severe (ordinal score < 4) covid subjects (# p < 0.1, * p < 0.05, ** p < 0.01, *** p < 0.001, ns = not significant). Inset density plots depict the distribution of respective lipids across the moderate and severe covid groups. Dotted line represents the median values.

#### Ether lipids are related to inflammatory pathways

To investigate the pathophysiological relevance of the observed lipids in the context of COVID-19 infection, we extracted the immunotypes, proteins, eicosanoids and clinical parameters that significantly correlated (Spearman’s rho < -0.4 or > 0.4, p < 0.05) with the level of these three lipid species in the COVID – 19 patients (Supplementary Table 7). LPC-O-16:0 was significantly correlated with measures of inflammation (PCT, CRP) and clinical severity of COVID-19 (ordinal score at day 7; the ordinal score at enrolment and day 7).

Interestingly, sPLA_2_ was also significantly anti-correlated with LPC-O-16:0. In addition, several (17 each) immune response features and proteins were associated with the circulatory levels of LPC-O-16:0. For example, this lipid is significantly negatively correlated with the previously described immunotype 2 in COVID-19 (35).

PC-O-30:0 was associated with 15 proteins and 11 immunotypes, but only to one clinical parameter, CRP (Supplementary Table 7). It is significantly negatively correlated with the previously described immunotype 1 in COVID-19 (35). Comparison of the proteins associated with the two lipids revealed largely orthogonal sets, suggesting that they are likely associated with different biological pathways. We performed functional protein enrichment analysis using string-db (Supplementary Table 8, Supplementary Figure 9) to interrogate further the degree of overlap. Several enriched pathways were significantly associated with LPC-O-16:0 (FDR < 0.001, Figure 6A, B) resulting in a strongly interconnected network, centered around the protein CXCL8 (Figure 6B). Such a well-defined network was not found from the proteins associated with PC-O-30:0 (Supplementary Figure 10). Thus, amongst the two ether lipids, LPC-O-16:0 is more closely related to the known disease pathophysiology.

**Figure 6:**
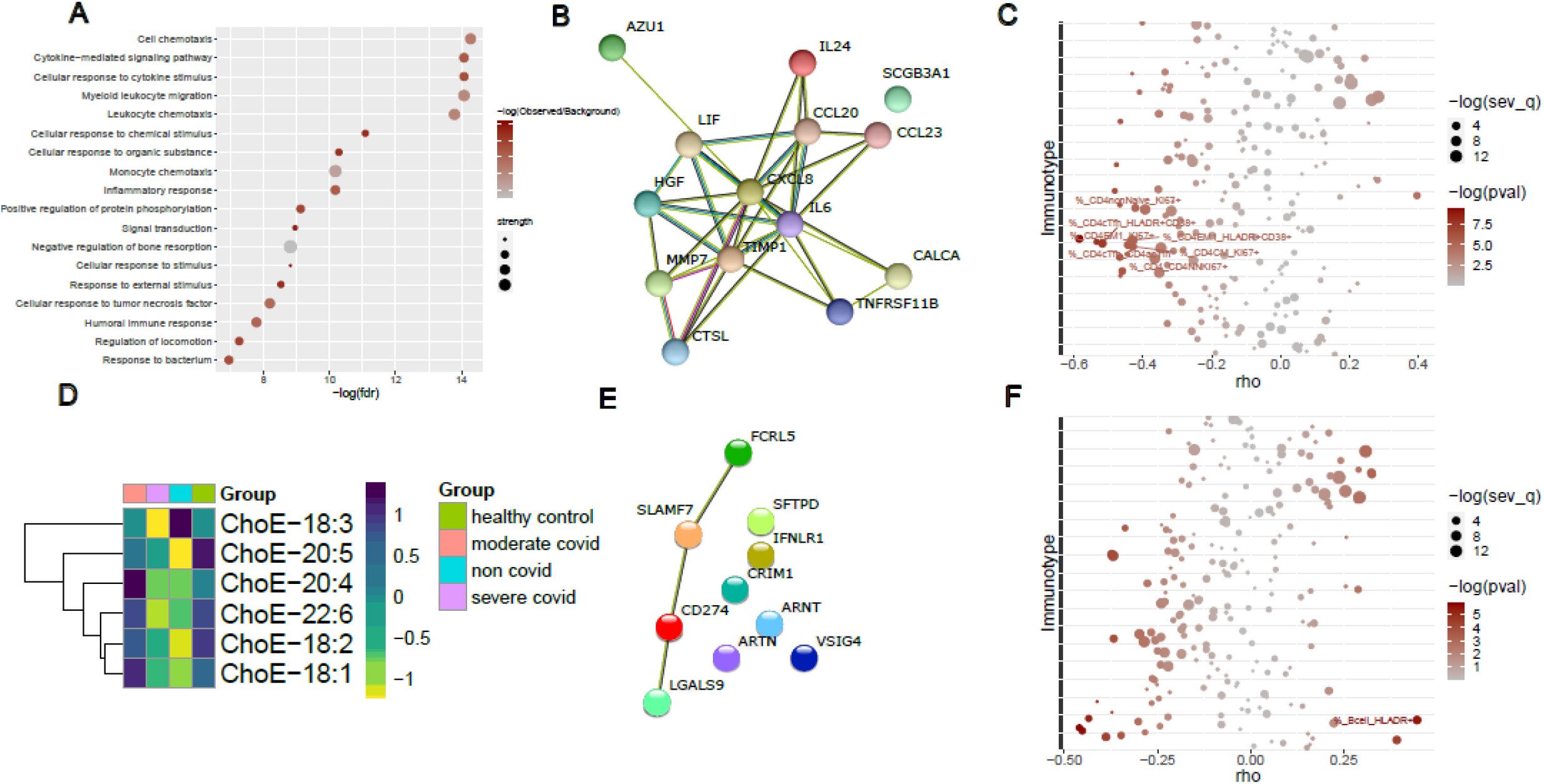
Highly abundant lipids restrain inflammation. LPC-O-16:0 is associated to inflammatory response (A – C) and ChoE-18:3 is specific to severe covid (D – F). Levels of LPC-O-16:0 from covid subjects were used for correlation analysis with protein and immunotype data to identify biological pathways associated with the lipid. A) Top functionally enriched pathways (FDR < 0.001, for all significantly enriched pathways (FDR < 0.05), see Supplementary Table 8) associated to the protein sets that are significantly associated to LPC-O-16:0 (Spearman’s rho < -0.4 and p < 0.05). B) Functional enrichment network of proteins significantly associated to LPC-O-16:0 (Spearman’s rho < -0.4 and p < 0.05). C) Immune cell types significantly correlated to LPC-O-16:0 (Spearman’s rho < -0.4 or > 0.4 and p < 0.05). The points are colored by -log (p value of correlation) and sized by -log (q value between moderate and severe covid (q_sev)). D) ChoE-18:3 is the only cholesteryl ester molecule that is depleted in the severe covid group compared to moderate covid, but elevated in the non-covid subjects. E) Functional enrichment network of proteins significantly associated (Spearman’s rho < -0.4 or > 0.4 and p < 0.05) to ChoE-18:3. F) ChoE-18:3 is significantly, and exclusively, associated to various B cell populations (Spearman’s rho < -0.4 or > 0.4 and p < 0.05)

Among the top five enriched pathways, two broad categories were apparent – cellular chemotaxis and cytokine signaling pathways. To identify the most probable tissue locations enriched by the proteins associated with LPC-O-16:0, we investigated the tissue expression enrichment of the string-db output (Supplementary Figure 11, Supplementary Table 9). Nine tissues were found to be enriched significantly (FDR < 0.05), with neutrophils being the most prominent. Given that the proteins are all negatively associated with LPC-O-16:0 that decreases with severity, it is plausible that this lipid is related to the inflammatory response towards the infection that is mediated by leucocyte chemotaxis and associated cytokine stimulation.

We identified features of CD4+ T cells (Figure 6C) that are both significantly associated with LPC-O-16:0 and significantly differ as a function of disease severity (FDR < 0.05). Activation or proliferation of CD4+ T cell subsets, including memory and T helper follicular cells inversely correlate with LPC-O-16:0, again consistent with a role as a constraint on the inflammatory response to a severe COVID-19 infection, as previously suggested (55). Moreover, this species, and ether-LPCs in general have been shown by others to be downregulated in sepsis, not just in severe COVID -19 (25, 55). Correspondingly, we found LPC-O-16:0 to be negatively correlated with sPLA_2_.

#### ChoE-18:3 is specifically related to severe COVID-19

Cholesteryl ester-18:3 (ChoE-18:3) was strikingly depleted in severe COVID-19 infection (Fig. 6D). It was decreased compared to controls and as a function of severity in COVID-19, however, unlike the ether lipids, it is elevated in the non-COVID-19 sepsis patients, compared to COVID-19 patients with severe disease. Interestingly, none of the other ChoE molecules that we profiled showed a similar pattern (Supplementary Figure 12, Figure 6D). We examined associated proteins, peripheral immune cell responses and clinical parameters to identify the biological and pathophysiological relevance context of this lipid (Spearman’s ρ< -0.4 or > 0.4, p < 0.05, Supplementary Table 7). ChoE-18:3 was significantly associated with only ten proteins (Supplementary Table 7) and, by implication, a limited number of pathways (Figure 6E). Functional enrichment analysis, however, revealed several significant pathways (Supplementary Table 8), the majority of which was related to regulation of immune response, with negative regulation of T cell proliferation being the most significant.

We next sought cellular immune responses associated with the cholesteryl esters. Interestingly, only five B cell populations were correlated with ChoE-18:3 (Supplementary Table 7, Spearman’s rho < -0.4 or > 0.4 and p < 0.05). Such exclusive association with B cells was not observed with the other cholesteryl ester species. HLADR+ B cells were positively correlated, while CD39/CD138/KI67 positive B cells were negatively correlated with ChoE-18:3(Supplementary Table 7). Among the B cell species significantly associated to ChoE-18:3, HLADR+ cells were also significantly different between moderate and severe COVID-19 patients (Figure 6F). As such, HLADR+ B cells showed positive association with two cholesteryl ester species – ChoE-18:3 and ChoE-22:6 (Supplementary Figure 13), each arising from fatty acids (FA 18:3 –LA and FA 22:6 –DHA) that bear precursor-product relationships. HLADR+ B cells are reduced in patients with severe COVID-19 (56) compared to healthy controls and patients with mild COVID independent of dexamethasone treatment. ChoE-18:3 is a potential source of the elevated ALA (alpha-linolenic acid) and its downstream products, EPA and DHA that we observed in COVID patients and was also negatively correlated with PLA_2_G2D (Supplementary Figure 14, Spearman’s ρ= -0.55, p < 0.05), a source of inflammatory eicosanoids as described above.

## Integrative correlation analysis

### COVID-19 severity correlation network

The UMAP projection of COVID-19 correlation network is shown in Figure 7A. Host response cluster 1 in this analysis contains the feature defining immunotype 1, associated with disease severity and mortality as previously described in COVID-19 (35). Most features in our host response cluster 2 are highly positively correlated with the feature defining immunotype 2 in that publication (35). The actual defining feature is located in the upper left corner of the boundary rectangle of the cluster. Our severity cluster includes severity scores such as APACHE III and the COVID-19 severity ordinal scale scores (which were evaluated four times throughout the month after hospitalization), length of hospital stay and indication of survival within one and three months after hospital admission.

**Figure 7:**
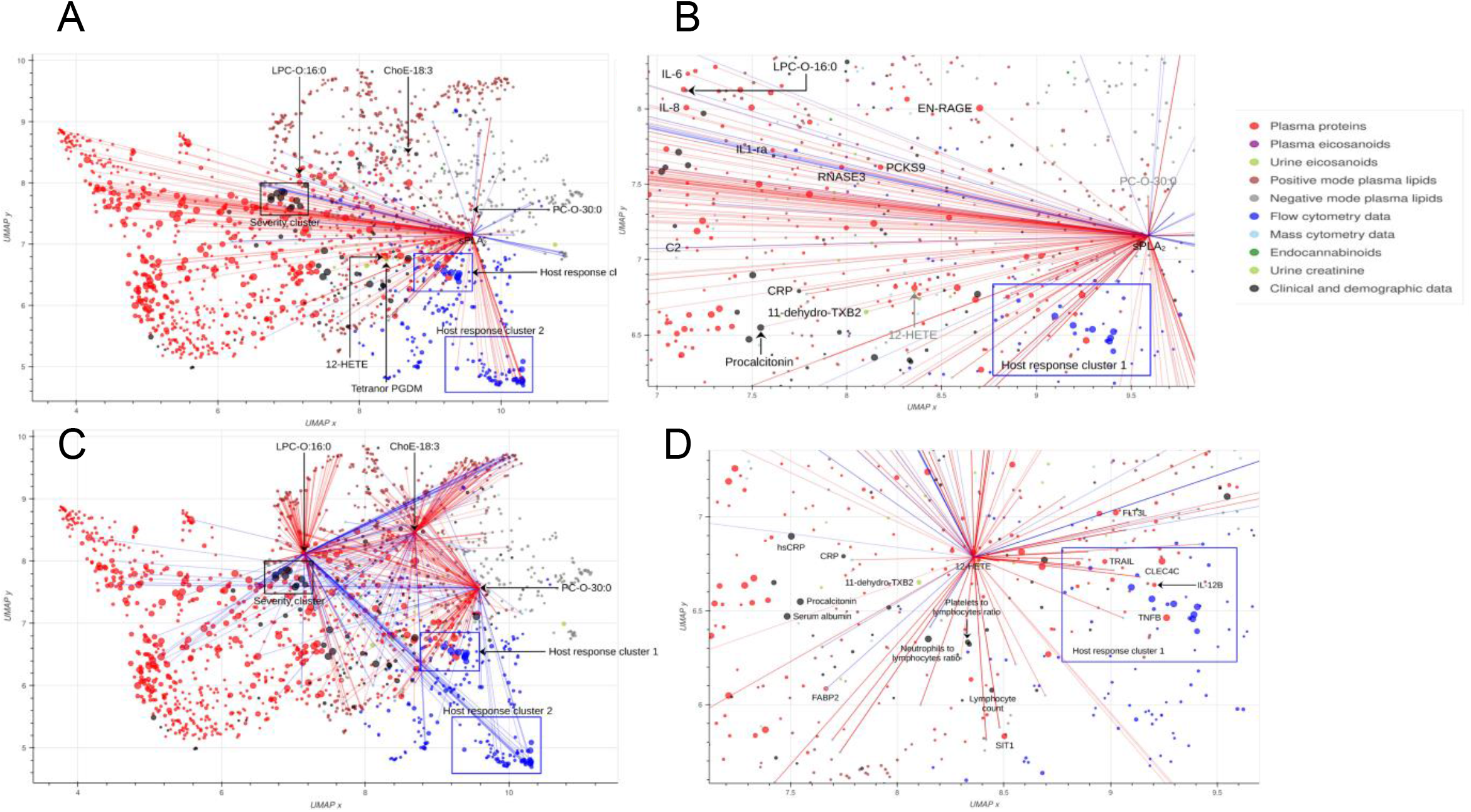
Visualization of COVID-19 correlation network. (A) UMAP projection of features measured in COVID-19 cohort Each feature is represented by a circle whose size is negatively proportional to log10 of the p-value testing difference of the feature in moderate vs. severe COVID-19. Hence, bigger circles correspond to features that differ more significantly between moderate and severe cases of COVID-19. Correlations of sPLA2 with other features are represented by lines, red lines correspond to positive correlations, with Spearman’s correlation coefficient > 0.4, blue lines correspond to negative correlations, with Spearman’s correlation coefficient < −0.4. Named features are further discussed in this paper. (B) **Region in close proximity to host response cluster 1 containing sPLA_2_.** Features printed in black show significant (p < 0.05) correlation with sPLA2, either with Spearman’s correlation coefficient > 0.4 (red lines) or < −0.4 (blue lines). (C) **Correlation subnetwork for three high abundance lipids of interest.** Red lines denote pairs of features with Spearman’s correlation coefficient > 0.4, blue lines denote pairs of features with Spearman’s correlation coefficient < −0.4. (D) **Region in close proximity to host response cluster 1 containing urinary 12-HETE and several other urinary eicosanoids.** Correlations of 12-HETE with other features are represented by red (Spearman’s correlation coefficient > 0.4) or blue (Spearman’s correlation coefficient < −0.4) lines. Compared to sPLA2, 12-HETE exhibits lower number of strong correlations with other features. However, the positioning of 12-HETE in the network is based on all correlations among all pairs of features, so its proximity to other features, such as IL-12B, TNFB, and SIT1 could be of interest.

Amongst the lipids mentioned previously, PC-O-30:0 is negatively correlated with several features in host response cluster 2 and is positioned relatively close to host response cluster 1 (Figure 7A, C). Similar to PC-O-30:0, LPC-O-16:0 is negatively correlated with host response cluster 1, while ChoE-18:3 is negatively correlated with a third, B cell related, cluster of immune response related features positioned left of the second one (Figure 7C). Urinary 12-HETE is located close to host response cluster 1 (Figure 7A, D) and so are several immune response related proteins, many of which differ significantly in their abundance in moderate vs. severe COVID-19 (such features are represented by larger circles in the figure), including IL-12B, CLEC4C, FLT3L, TRAIL, and TNFB. 12-HETE is also close to SIT1, which interacts with the ACE2 receptor for SARS-CoV-2 (57). Lymphocyte count, NLR and PLR are clinical immune related features located below 12-HETE in the region shown in Figure 7D. Many immune response related proteins correlate significantly with sPLA_2_ (Figure 7B).

### Non-COVID sepsis control vs. COVID-19 differential correlation network

The UMAP projection of non-COVID control vs. COVID-19 differential correlation network is shown in Supplementary Figure 15A. 12-HETE in plasma exhibited changes in correlation with multiple features (Supplementary Figure 15A). STRING enrichment analysis (58) of 87 proteins differentially correlated with 12-HETE was performed (Supplementary Figure 15B). The top two KEGG pathways, as ranked by FDR, were *Viral Protein Interaction with Cytokines and Cytokine Receptor* (FDR adjusted p-value 8.57·10^-12^) and *Cytokine-Cytokine Receptor Interaction* (FDR adjusted p-value 2.12·10^-18^). These 87 proteins generally exhibited loss of correlation with 12-HETE in the COVID-19 cohort (Supplementary Figure 15C). Analysis of the undirected differential correlation graph placed 12-HETE in the top 5% of features in terms of degree centrality and betweenness centrality. Proteins in the top 5% of features ranked by degree centrality (52 proteins in total) were enriched in the Neutrophil Degranulation Reactome Pathway (FDR adjusted p-value 0.00025).

## DISCUSSION

Viruses perturb the cellular lipidome, which is, in turn, crucial to their replication and propagation (1,2,59,60). Aside from these direct effects, lipids modulate powerfully the immune response to viral infection that restricts its consequences or, if unrestrained, may amplify the insult to the infected host. Here, we interrogated this interaction across the spectrum of severity of infection with COVID-19 with a particular focus on the less abundant, but biologically potent eicosanoid products of AA and LA. Indeed, the spike (S) glycoprotein on the surface of SARS-CoV-2 tightly binds LA, stabilizing it and reducing its interaction with the host angiotensin-converting enzyme 2 (ACE2) receptor that facilitates viral cell entry (61).

This study contrasts with other investigations of the lipidomic response to COVID-19 (i) by including both high and low abundance lipids; (ii) by comparing patients with sepsis due to COVID -19 with patients suffering from sepsis due to other viruses and to bacteria, as well as with healthy volunteers, (iii) by integrating lipidomics with the proteome and the peripheral cellular immune response, allowing for comparisons over time from hospital admission and with the severity of the clinical response, (iv) by tracking changes in the lipidome in response to mild infections resulting in seroconversion in healthy volunteers and (v) by providing a derived inflammatory connectome as a tool to the community for further hypothesis generation.

Eicosanoids are autacoids, not circulating hormones, but are detectable in plasma or urine, reflecting endogenous biosynthesis. They are cleared rapidly from plasma to urine where primary products or metabolites are acquired non-invasively and in greater abundance than in plasma. The cellular capacity to form eicosanoids exceeds their endogenous production in vivo (62).

Here we find that sepsis, of bacterial or viral origin, is accompanied by an eicosanoid storm with elevated levels of COX (PGE_2_, PGD_2_, TxA_2_), LOX (12-HETE and LTE_4_) and EPOX products (11(12) – 14(15)-DHETs) of AA, the endocannabinoids AEA and 2-AG, along with the LA derived DiHOMEs. Furthermore, elements of this storm reflect disease severity and are prognostic indicators in patients with sepsis due to COVID-19. This storm is accompanied by an increase in sPLA_2_ that metabolizes the AA and LA substrates to generate these bioactive lipids. Indeed, sPLA_2_ acts as a prominent immune-lipidomic hub, discriminates between moderate and severe sepsis in COVID-19 and links eicosanoids to multiple inflammatory proteins and to clinical markers of disease severity. While sPLA_2_ has been previously reported as a biomarker of sepsis (25, 61), here we expand this observation mechanistically to link sPLA_2_ to the breadth of its discrete eicosanoid products, the immune cellular responses that they influence and the consequent indices of disease severity.

In our study, the patients with sepsis due to causes other than COVID -19 had, on average, more severe disease than those with COVID-19. In the main, eicosanoids (TxA_2_, PGE_2_, AEA, 2-AG, DHETs and DiHOMEs) reflected comparative disease severity, but elevations in PGD_2_ and 12-HETE were more striking in COVID-19 despite less severe disease, reflecting a relative selectivity for SARS-CoV-2. Interestingly, bacterial, rather than viral sepsis evoked the more integrative eicosanoid response across the COX, LOX and EPOX pathways.

PGD_2_ is a potent immune modulator, acting via DPr1 and DPr2 – D Prostanoid receptors. DPr1 signaling delays migration of DCs to the lung and lymph nodes via downregulation of the chemokine CCR7 (63). PGD_2_ also contributes to the pathogenesis of respiratory syncytial virus (RSV) bronchiolitis and susceptibility to asthma via DPr2 signaling (64). In a neonatal model of severe RSV bronchiolitis, treatment with a DPr2 inhibitor decreased viral load and improved morbidity via upregulation of IFN-λ. This effect was recapitulated by treatment with a DPr1 agonist, suggesting that these two receptors for PGD_2_ have opposing roles in the regulation of the antiviral response (65). Strikingly, middle-aged mice lacking expression of DPr1 or PLA_2_G2D are protected from severe disease in a model of SARS-CoV-2 infection and a DPr1 antagonist, asapiprant, protected aged mice from lethal infection (66). Here, PLA2G2D was not elevated in patients with sepsis compared to healthy controls, suggesting that sPLA_2_ is the dominant driver of PGD_2_ formation in humans with COVID-19.

Less is known about 12-HETE and yet it emerges also as an important hub linking inflammatory proteins, peripheral immune cells, host response cluster 1 and clinical indices of disease severity. Both plasma TxB_2_ and 12-HETE are prone to platelet activation as a sampling artifact that is avoided by measuring TxM and 12-HETE in urine. 12(S) – HETE is the product of ALOX12 in platelets. 12 (R) HETE is made by ALOX12B, mostly expressed in skin and cornea or as a minor product of ALOX15 which more widely expressed, including in the vasculature. Although we did not characterize the stereoisomer, the coincident increase in the platelet COX product TxA_2_, implicates platelet 12 (S) – HETE as the form elevated in sepsis due to COVID-19. Deletion of ALOX12 results in increased platelet sensitivity to ADP induced aggregation ex vivo in mice (67) and Holinstat and colleagues (68) have shown that ALOX12 plays an essential role in regulating FcγRIIa immune mediated thrombosis. Thus, increased 12 (S) - HETE might augment the prothrombotic consequences of the increased TxA_2_ generation that we observed in sepsis and particularly in that due to COVID-19.

Increased biosynthesis of PGE_2_ is a feature of sepsis and its role as a pro-inflammatory mediator is well established (4). It can activate any of 4 EPrs that differentially regulate platelet and immune cell function. Elevated levels of PGE_2_ formation have been reported by others in patients with sepsis from COVID-19 compared to healthy controls, but here we see no distinction from the increase observed in patients with sepsis from other causes. As previously observed, PGE_2_ production by PBMCs from COVID-19 patients is elevated compared to healthy controls. In addition, PGF_2α_ was significantly increased in the supernatant of PBMCs from COVID-19 patients and its level correlated with broad populations of immune cells. This is of interest as deletion of the FPr restrains the pulmonary fibrotic response to bleomycin in mice (69) and increased biosynthesis of PGF_2α_ might contribute to the pulmonary fibrosis that can complicate long COVID-19 (70). Interestingly, Bohnacker et al reported that PGF_2α_ levels remain elevated in monocyte-derived macrophages from COVID-19 patients up to 3–5 months post infection (24). We observed a range of other eicosanoid – immune cell relationships in PBMCs from patients with COVID-19.

Our extensive analysis of highly abundant lipids revealed three ether lipids that were differentially altered in the sepsis patients due to COVID-19. All three - ChoE-18:3, LPC-O-16:0 and PC-O-30:0 - were significantly lower in patients with sepsis from COVID-19 than sepsis from other causes and, indeed, from healthy controls. All three discriminated moderate from severe COVID-19. LPC-O-16:0 is also depleted in patients with non-COVID-19 sepsis relative to healthy controls. Negative correlation of these lipids with PLA_2_s and other lipid and protein markers of the inflammatory response infer their importance as restraints on inflammation.

Previous immune profiling of patients from the MESSI cohort with COVID-19 has identified three distinct immunotypes (35). The first was associated with disease severity and showed robustly activated CD4 T cells, a paucity of circulating follicular helper cells, activated CD8 “EMRAs,” hyperactivated or exhausted CD8 T cells, and plasmablasts. Immunotype 2 was characterized by less CD4 T cell activation, Tbet+ effector CD4 and CD8 T cells, and proliferating memory B cells and was not associated with disease severity. Immunotype 3, which negatively correlated with disease severity and lacked obvious activated T and B cell responses, was also identified. Here, interrogation of the connectome reveals significant correlations between both PC O:30-0 and LPC-O-16:0 with host response cluster 2 and between ChoE-18:3 and the third host response cluster.

We anticipated that a milder lipidomic inflammatory phenotype might accompany seroconversion after infection by SARS-CoV-2 in healthy marines, perhaps most evident in those with accompanying symptoms. Such a mild lipidomic response was, indeed, evident. There was an elevation of LA and its DiHOME metabolites (but not of AA derived eicosanoids) as we had seen in patients with sepsis, but surprisingly, only in the marines who seroconverted asymptomatically.

These studies deepen our understanding of how the lipidome drives the immune response across the spectrum of severity of sepsis. Besides expanding the breath of the eicosanoid storm that characterizes sepsis, we identify immuno-lipidic hubs that promote inflammation – sPLA_2_, PGD_2_ and 12-HETE – and exhibit relative specificity for COVID-19. Additionally, we identify hubs amongst the more abundant lipids, ChoE-18:3, LPC-O-16:0 and PC-O-30:0, that restrain inflammation. These abundant lipids are depressed in patients with sepsis due to COVID-19 compared to those with sepsis from other causes and with healthy controls. Finally, we provide a network analysis tool that displays the topical relationship of these hubs with previously described COVID-19 immunotypes and allows for interrogation by the community to generate novel mechanistic and therapeutic hypotheses.

## Support

This work was supported by grants from the NIH U54TR001878 (GAF), AI105343, AI082630, AI108545, AI155577, AI149680, U19AI082630 (EJW), HL142981 and NR018836-01 (AMW) and HL161196 (NJM). Additional support was provided from Defense Advanced Research Projects Agency contract number N6600119C4022 (S.C.S.) and Defense Health Agency grant 9700130 through the Naval Medical Research Center (A.G.L.).

DM and EJW are supported by the Parker Institute for Immunotherapy. EJW is also supported by the Perelman School of Medicine COVID Fund. NJM reports funding to her institution for unrelated work from Quantum Leap Healthcare Collaborative and BioMarck, Inc, and consulting fees from AstraZeneca Inc and Endpoint Health Inc. GAF is the McNeil Professor of Translational Medicine and Therapeutics and holds a Merit Award from the American Heart Association. He is an advisor to Calico Laboratories.

A.G.L is a military service member. This work was prepared as part of his official duties. Title 17, US Code §105 provides that copyright protection under this title is not available for any work of the US Government. Title 17, US code §101 defines a US Government work as a work prepared by a military service member or employee of the US Government as part of that person’s official duties. The views expressed in the article are those of the authors and do not necessarily express the official policy and position of the US Navy, the Department of Defense, the US Government or the institutions affiliated with the authors.

## Supporting information

All Supplementary Figures

All Supplementary Tables

