## Supplementary material for "Deep Phenotyping of the Lipidomic Response in COVID and non-COVID Sepsis": All Supplementary Figures

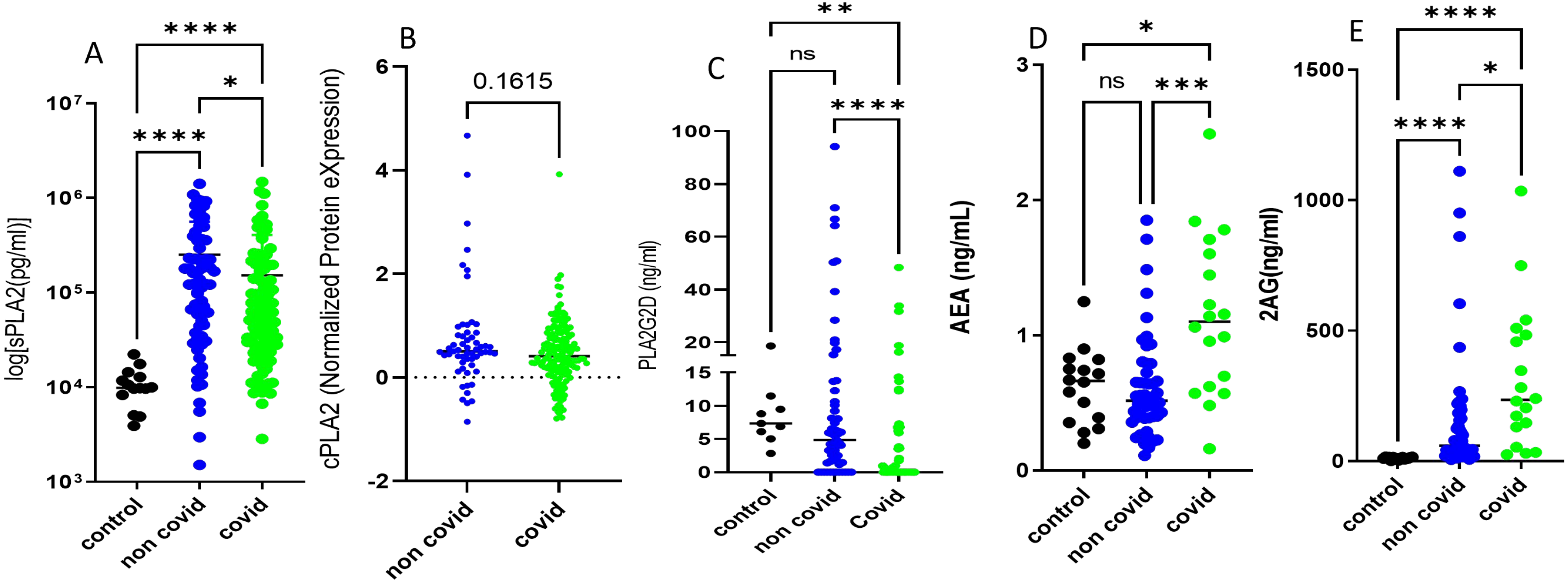

### Supplementary Figure 1.

**Plasma PLAs and endocannabinoids in control, non-covid and covid-19 patients. A) sPLA2 B) cPlA2 C) PLA2G2D D) AEA E) 2AG**

* P<0.05, ** P<0.01, *** P<0.001, **** P<0.0001.

**Suppl Fig. 2**

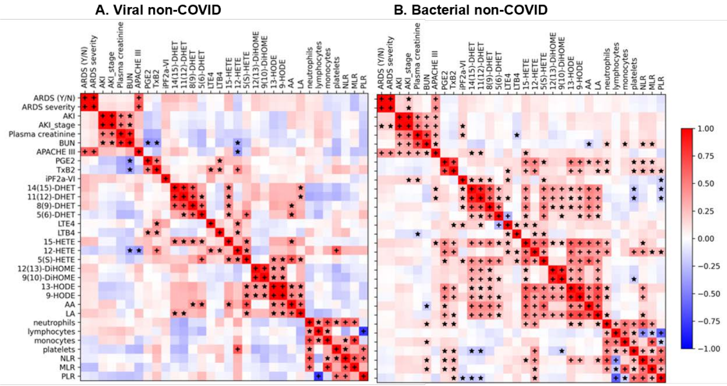

**Suppl Figure 2. Plasma lipidomes of critically ill patients with viral or bacterial infections.** Spearman's rank correlation analysis *FDR<0.05; ⁺FDR <0.001

Supplementary Figure 3

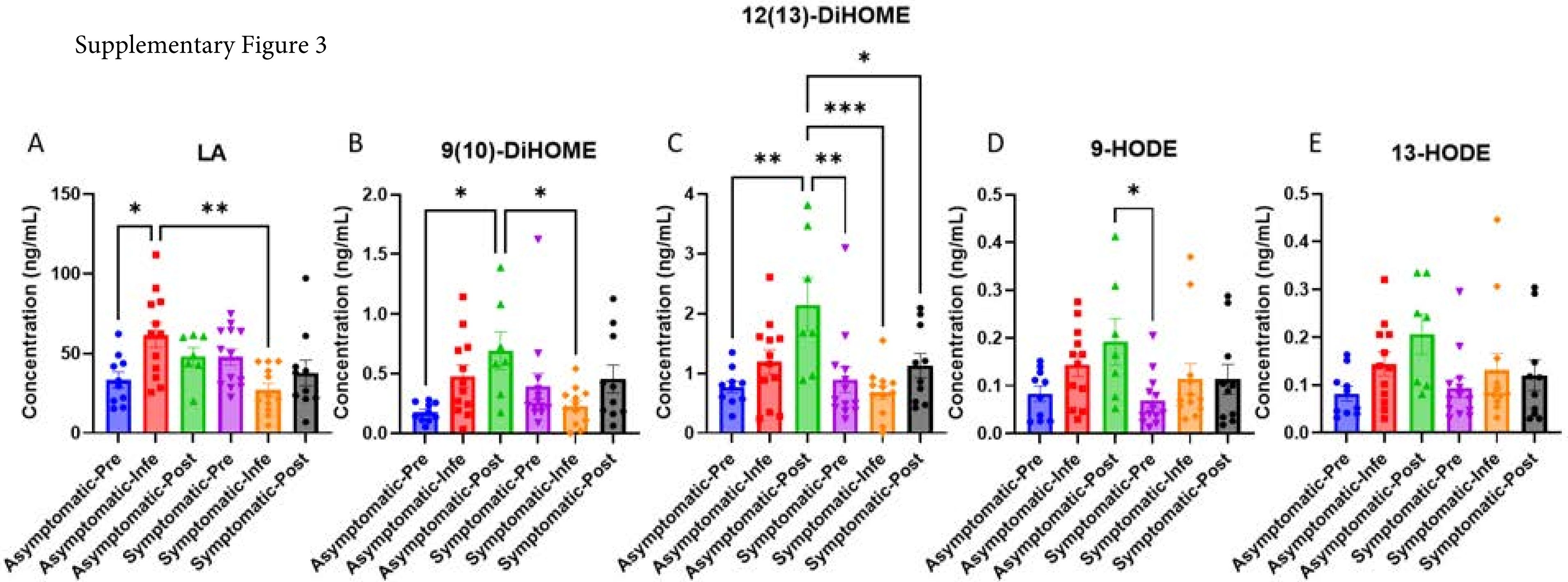

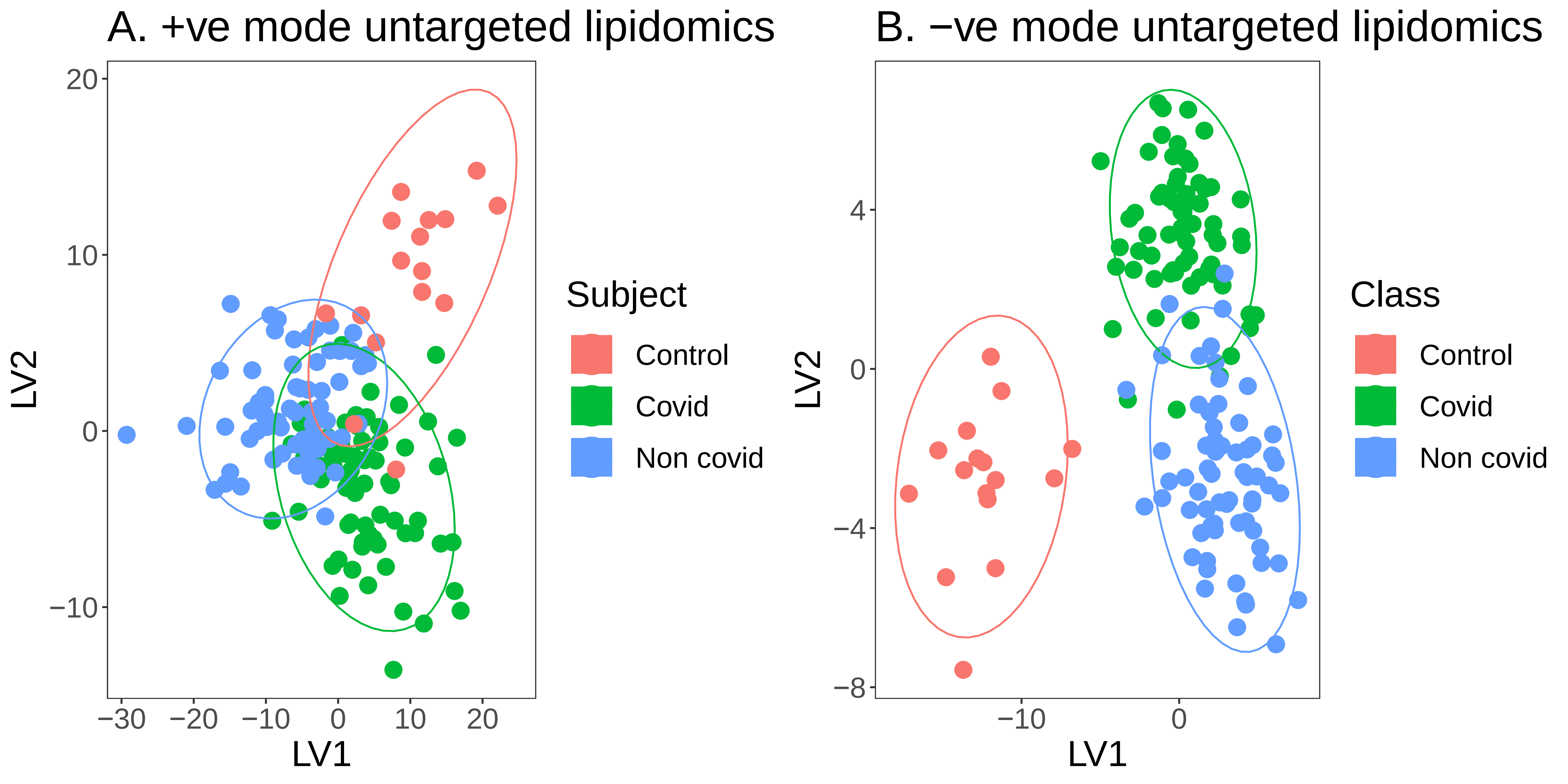
A. +ve mode untargeted lipidomics
B. −ve mode untargeted lipidomics
LV1
LV2
20
10
0
−10
−30
−20
−10
0
10
20
Subject
Control
Covid
Non covid
4
0
−4
−8
−10
0
Class
Control
Covid
Non covid

Supplementar Figure 4: Orthogonal partial least square discriminant analysis (OPLS-DA) scores plot using untargeted positive and negative mode lipidomics data from covid-19, non-covid and heatlhy control subjects

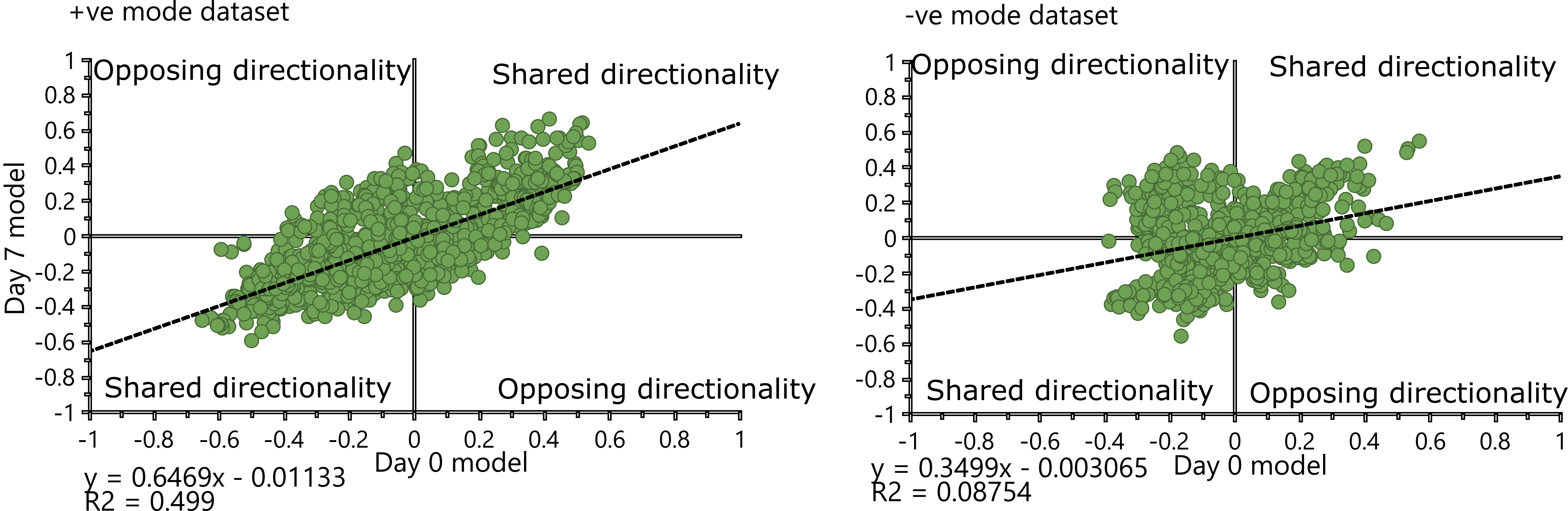

+ve mode dataset
Opposing directionality
Shared directionality
Shared directionality
Opposing directionality
Day 7 model
Day 0 model
y = 0.6469x - 0.01133
R2 = 0.499
-ve mode dataset
Opposing directionality
Shared directionality
Shared directionality
Opposing directionality
Day 0 model
y = 0.3499x - 0.003065
R2 = 0.08754

Supplementary Figure 5: Shared and unique structure plot depicting differential response of lipid features from covid, non-covid and healthy subjects from day 0 and day 7 samples

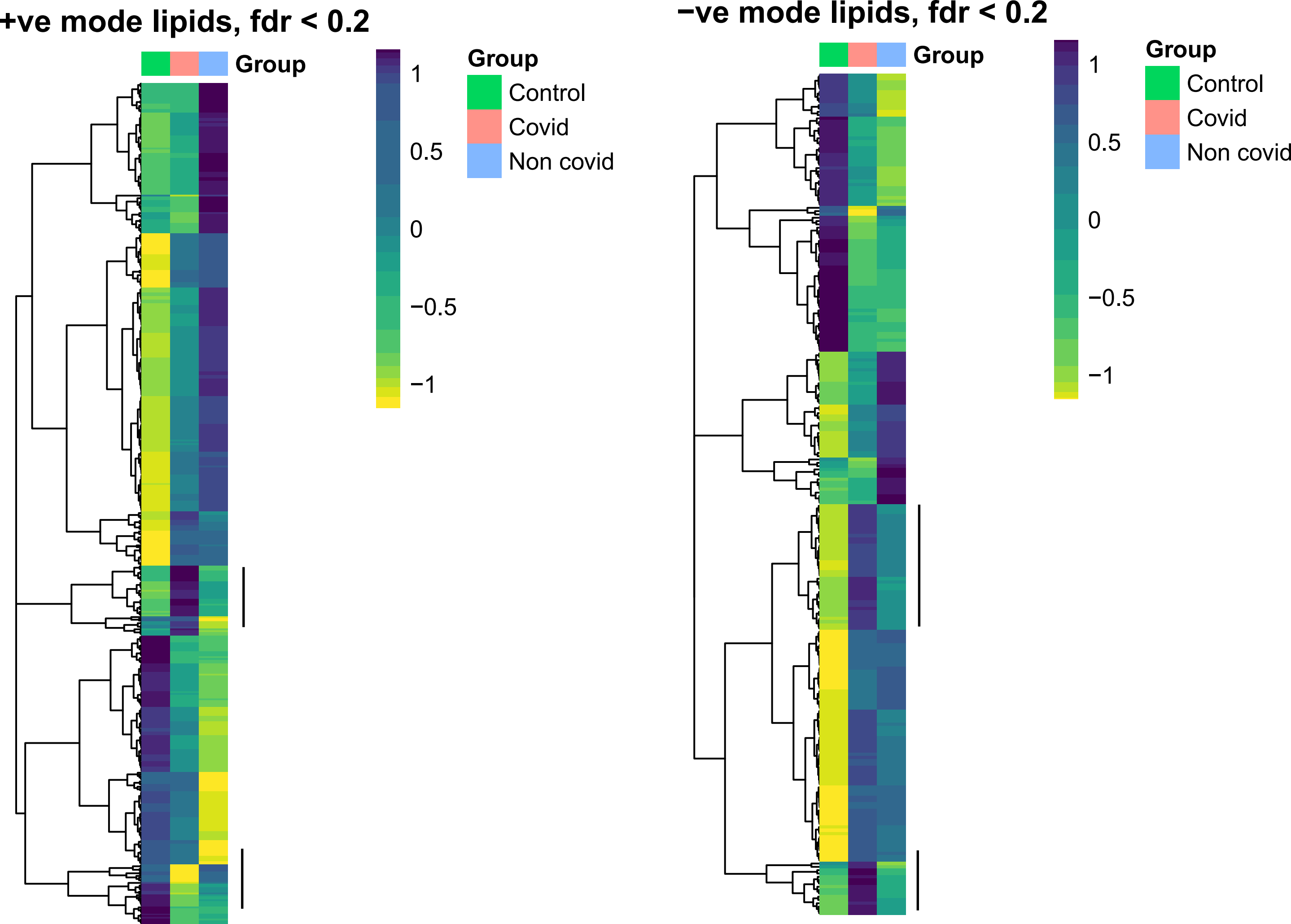

+ve mode lipids, fdr < 0.2
Group
1
0.5
0
−0.5
−1
Group
Control
Covid
Non covid
−ve mode lipids, fdr < 0.2
Group
1
0.5
0
−0.5
−1
Group
Control
Covid
Non covid

Supplementary Figure 6: Untargeted lipid features showing significant variance by Kruskal Wallis test between covid, non-covid and healthy subjects at day 0 (fdr < 0.2). Blocks labelled by vertical lines are cluster of features that changed similarly in covid compared to both non-covid ad controls.

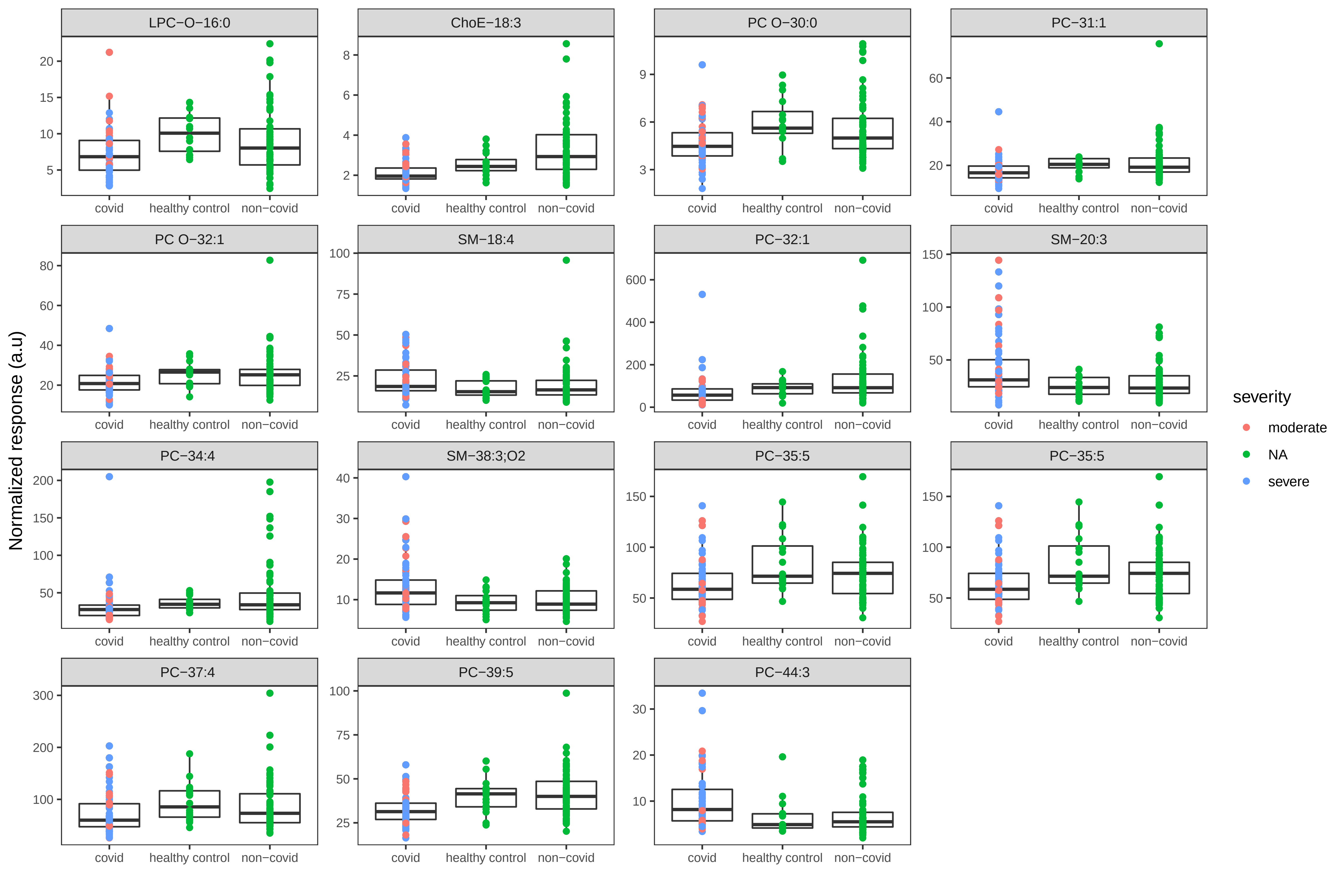
LPC−O−16:0
ChoE−18:3
PC O−30:0
PC−31:1
PC O−32:1
SM−18:4
PC−32:1
SM−20:3
PC−34:4
SM−38:3;O2
PC−35:5
PC−35:5
PC−37:4
PC−39:5
PC−44:3
covid
healthy control
non−covid
Normalized response (a.u)
severity
moderate
NA
severe

Supplementary Figure 7: Boxplots depicting positive mode covid specific lipids from day 0. The criterions used was same as that used in Figure 1C. Pink dots - moderate covid, blue dots - severe covid, green dots - healthy controls or non-covid.

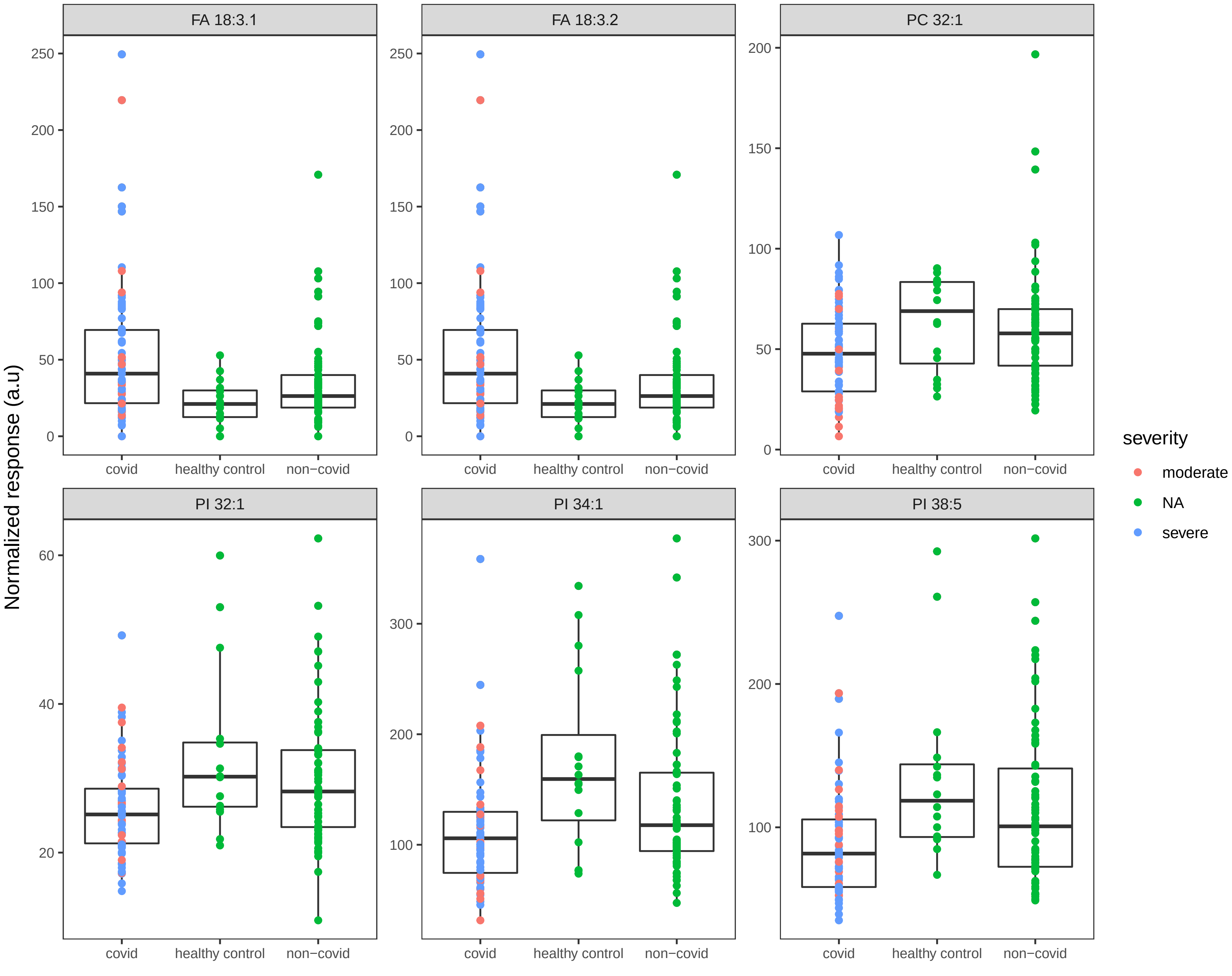

FA 18:3.1
FA 18:3.2
PC 32:1
PI 32:1
PI 34:1
PI 38:5
Normalized response (a.u)
covid
healthy control
non−covid
severity
moderate
NA
severe

Supplementary Figure 8: Boxplots depicting negative mode covid specific lipids from day 0. The criterions used was same as that used in Figure 1C. Pink dots - moderate covid, blue dots - severe covid, green dots - healthy controls or non-covid.

Biological processses enriched from proteins associated with LPC-O-16:0

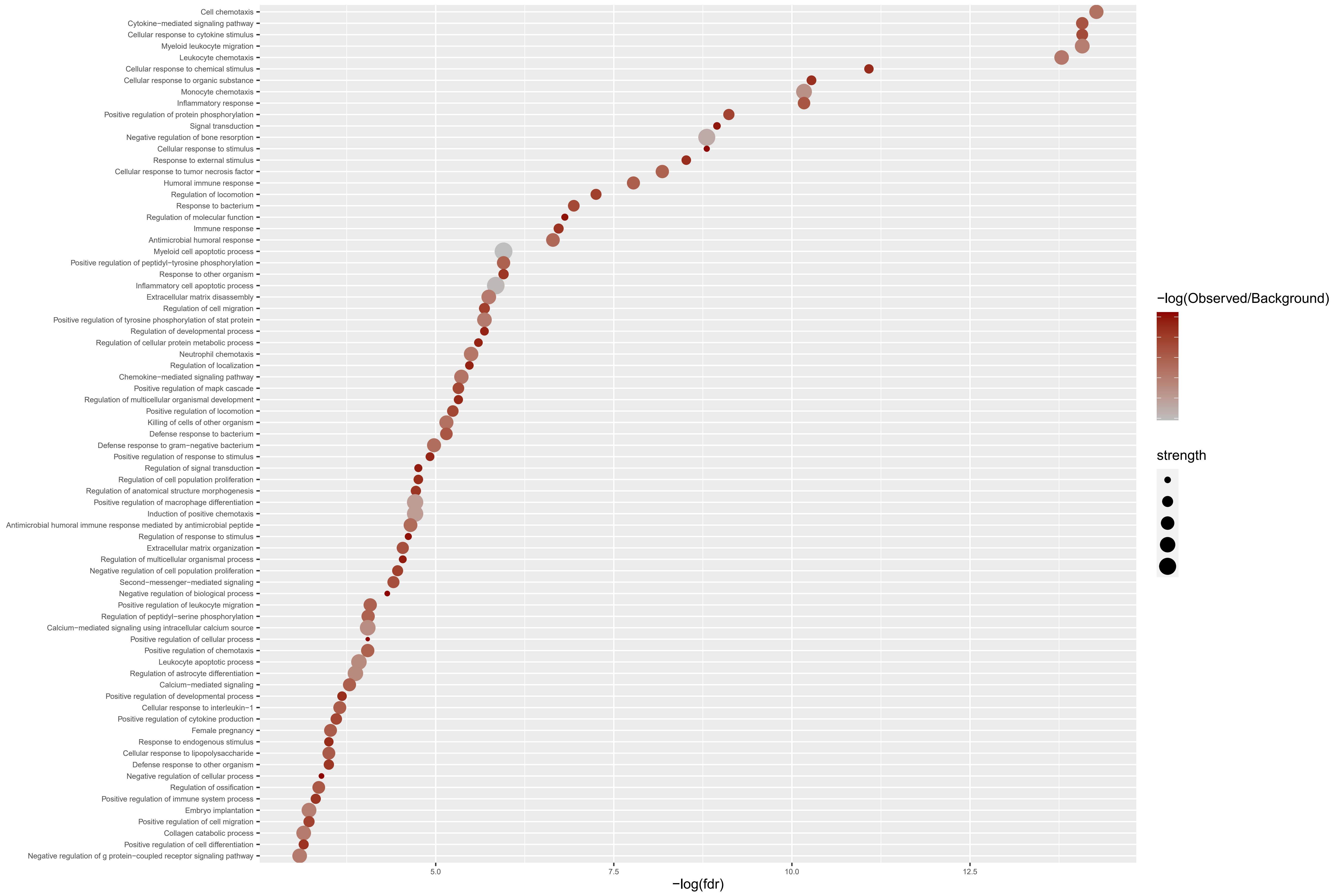

Biological processses enriched from proteins associated with PC-O-30:0

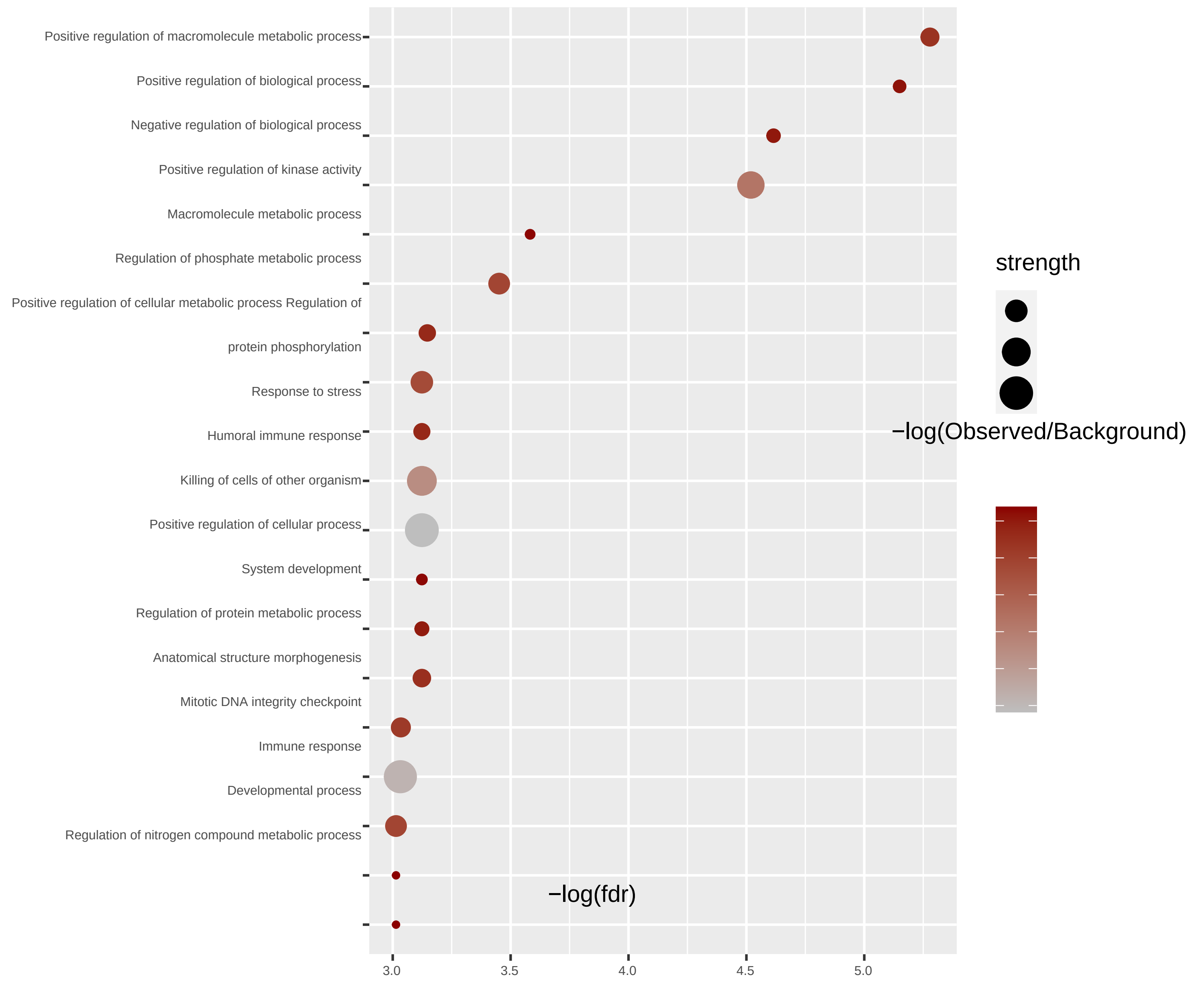

Supplementary Figure 9: Functionally enriched biological pathways (fdr < 0.05) obtained from proteins significantly correlated (Spearman's rho < -0.4 or > 0.4, and p < 0.05) to LPC-O-16:0 or PC-O-30:0

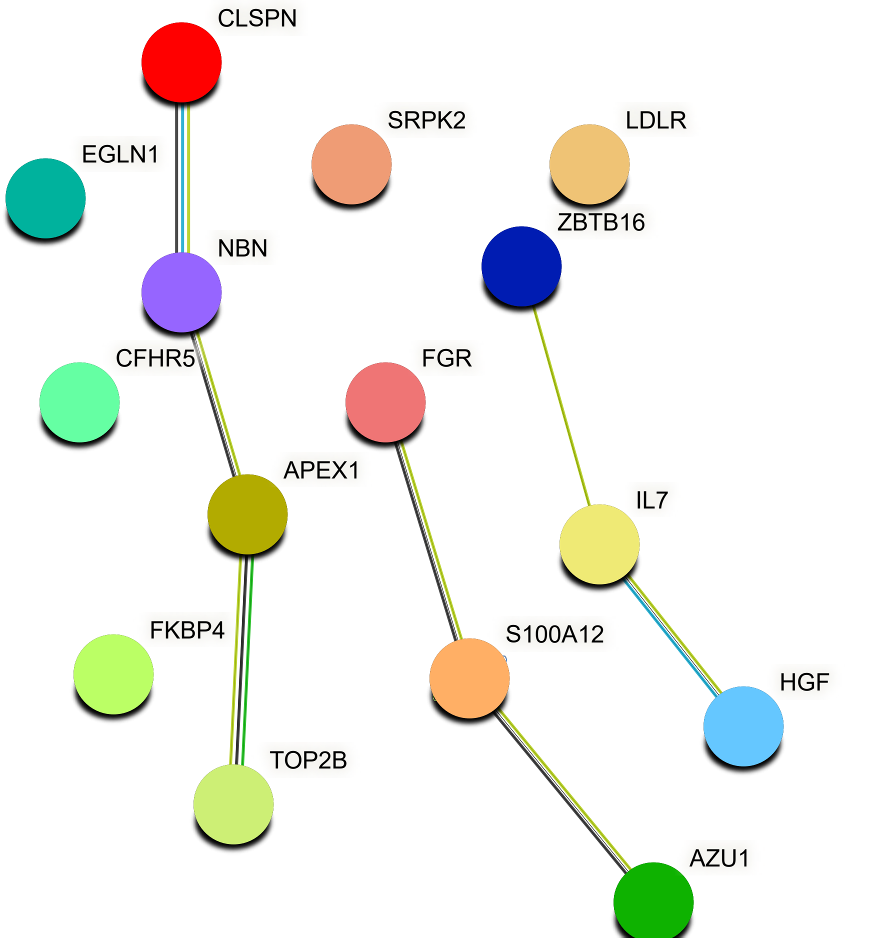
CLSPN
SRPK2
LDLR
EGLN1
ZBTB16
NBN
CFHR5
FGR
APEX1
IL7
FKBP4
S100A12
HGF
TOP2B
AZU1

Supplementary Figure 10:
Functional enrichment network of proteins significantly (refer to Supplementary Fiure S6) associated to PC-O-30:0.

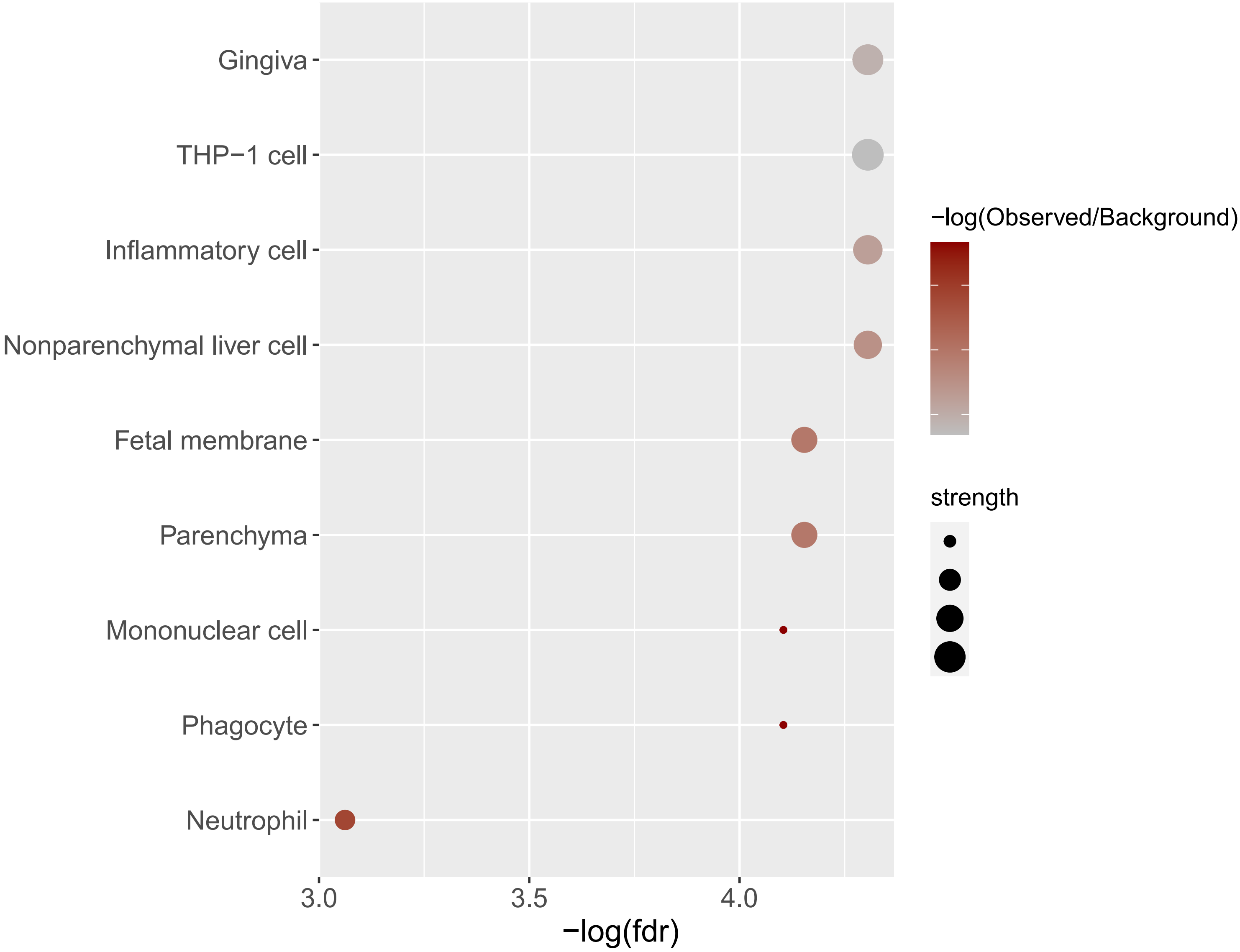

Supplementary Figure 11: Top enriched tissue locations of proteins significantly associated to LPC-O-16:0 (Spearman's rho < -0.4 and p < 0.05).

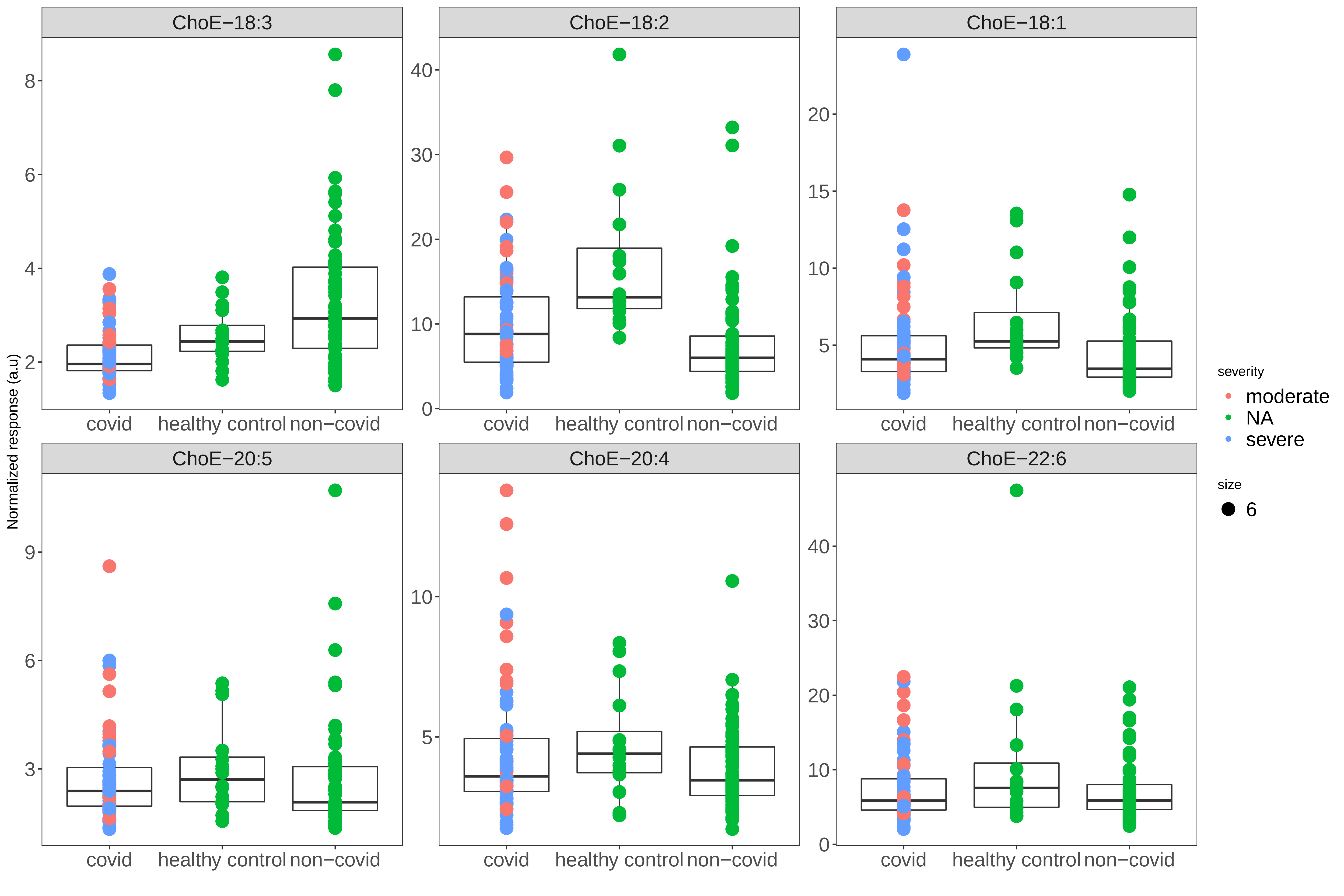

ChoE−18:3
ChoE−18:2
ChoE−18:1
ChoE−20:5
ChoE−20:4
ChoE−22:6
Normalized response (a.u)
covid
healthy control
non−covid
severity
moderate
NA
severe
size
6

Supplementary Figure 12: Levels of choletesteryl esters between covid (severe and moderate groups are labeled by different colors) , control and non-covid subjects

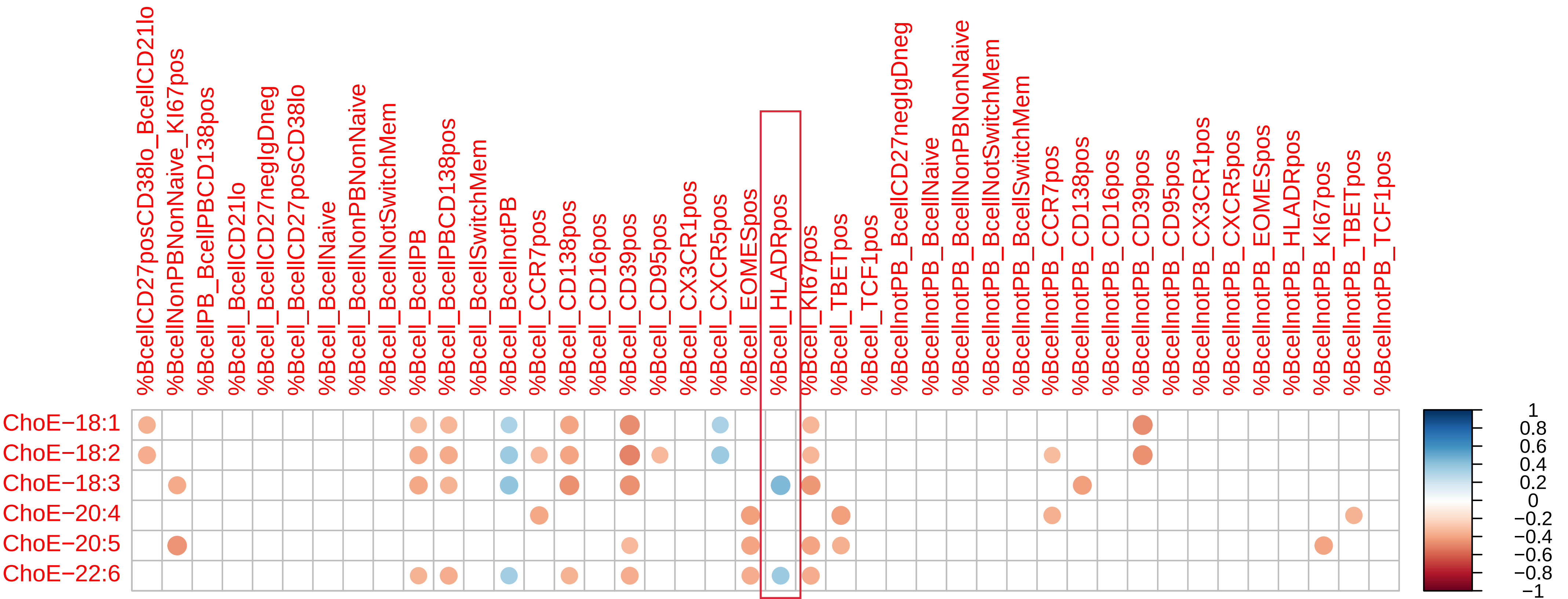
%BcellCD27posCD38lo_BcellCD21lo
%BcellNonPBNonNaive_KI67pos
%BcellPB_BcellPBCD138pos
%Bcell_BcellCD21lo
%Bcell_BcellCD27negIgDneg
%Bcell_BcellCD27posCD38lo
%Bcell_BcellNaive
%Bcell_BcellNonPBNonNaive
%Bcell_BcellNotSwitchMem
%Bcell_BcellPB
%Bcell_BcellPBCD138pos
%Bcell_BcellSwitchMem
%Bcell_BcellnotPB
%Bcell_CCR7pos
%Bcell_CD138pos
%Bcell_CD16pos
%Bcell_CD39pos
%Bcell_CD95pos
%Bcell_CX3CR1pos
%Bcell_CXCR5pos
%Bcell_EOMESpos
%Bcell_HLADRpos
%Bcell_KI67pos
%Bcell_TBETpos
%Bcell_TCF1pos
%BcellnotPB_BcellCD27negIgDneg
%BcellnotPB_BcellNaive
%BcellnotPB_BcellNonPBNonNaive
%BcellnotPB_BcellNotSwitchMem
%BcellnotPB_BcellSwitchMem
%BcellnotPB_CCR7pos
%BcellnotPB_CD138pos
%BcellnotPB_CD16pos
%BcellnotPB_CD39pos
%BcellnotPB_CD95pos
%BcellnotPB_CX3CR1pos
%BcellnotPB_CXCR5pos
%BcellnotPB_EOMESpos
%BcellnotPB_HLADRpos
%BcellnotPB_KI67pos
%BcellnotPB_TBETpos
%BcellnotPB_TCF1pos
ChoE−18:1
ChoE−18:2
ChoE−18:3
ChoE−20:4
ChoE−20:5
ChoE−22:6
1
0.8
0.6
0.4
0.2
0
−0.2
−0.4
−0.6
−0.8
−1

Supplementary Figure 13: Spearman's correlation of B cell populations of Covid-19 subjects with different cholesteryl ester levels. Only significant ($p < 0.05$) correlations are shown. HLADR positive B cell fraction is highlighted as it shows positive association with cholesteryl esters of FA 18:3 and 22:6

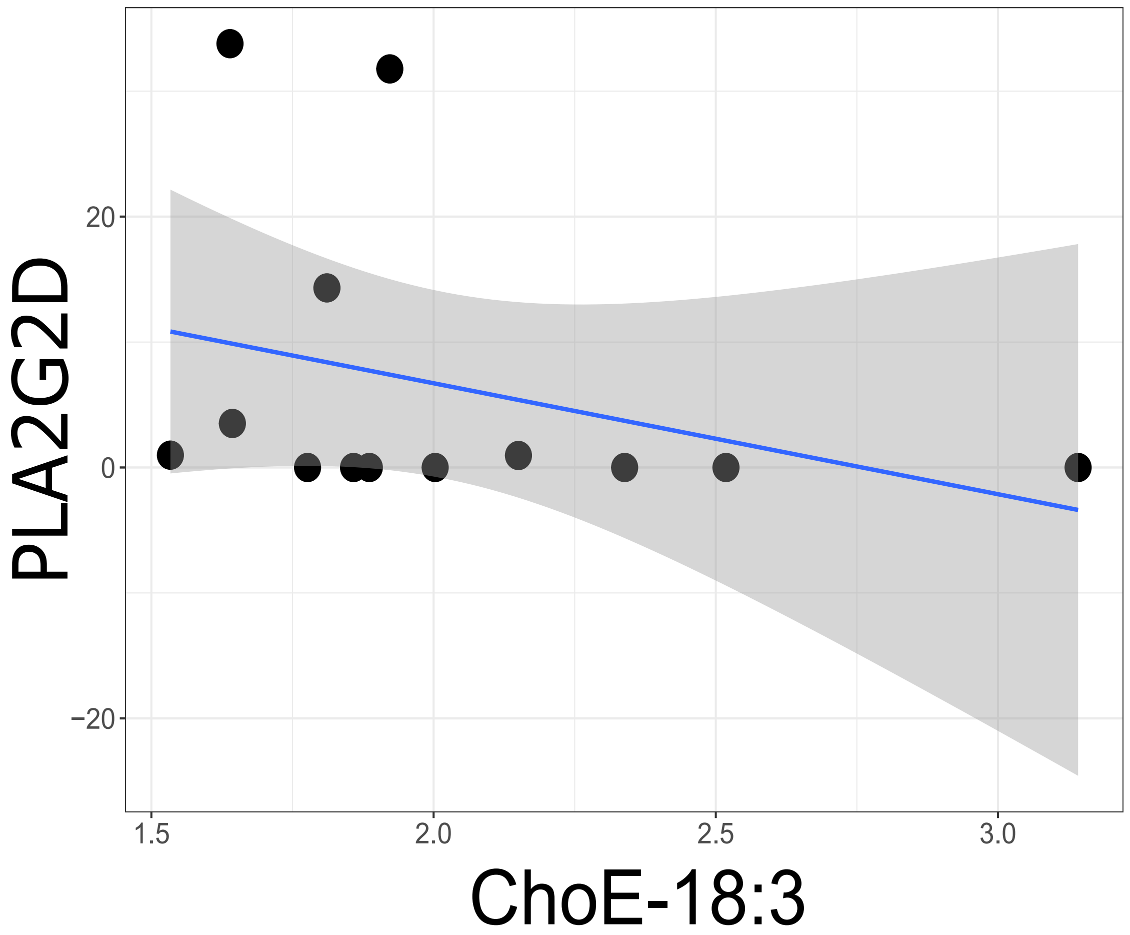

Supplementary Figure 14: Strong linear association of ChoE-18:3 with PLA2G2D level

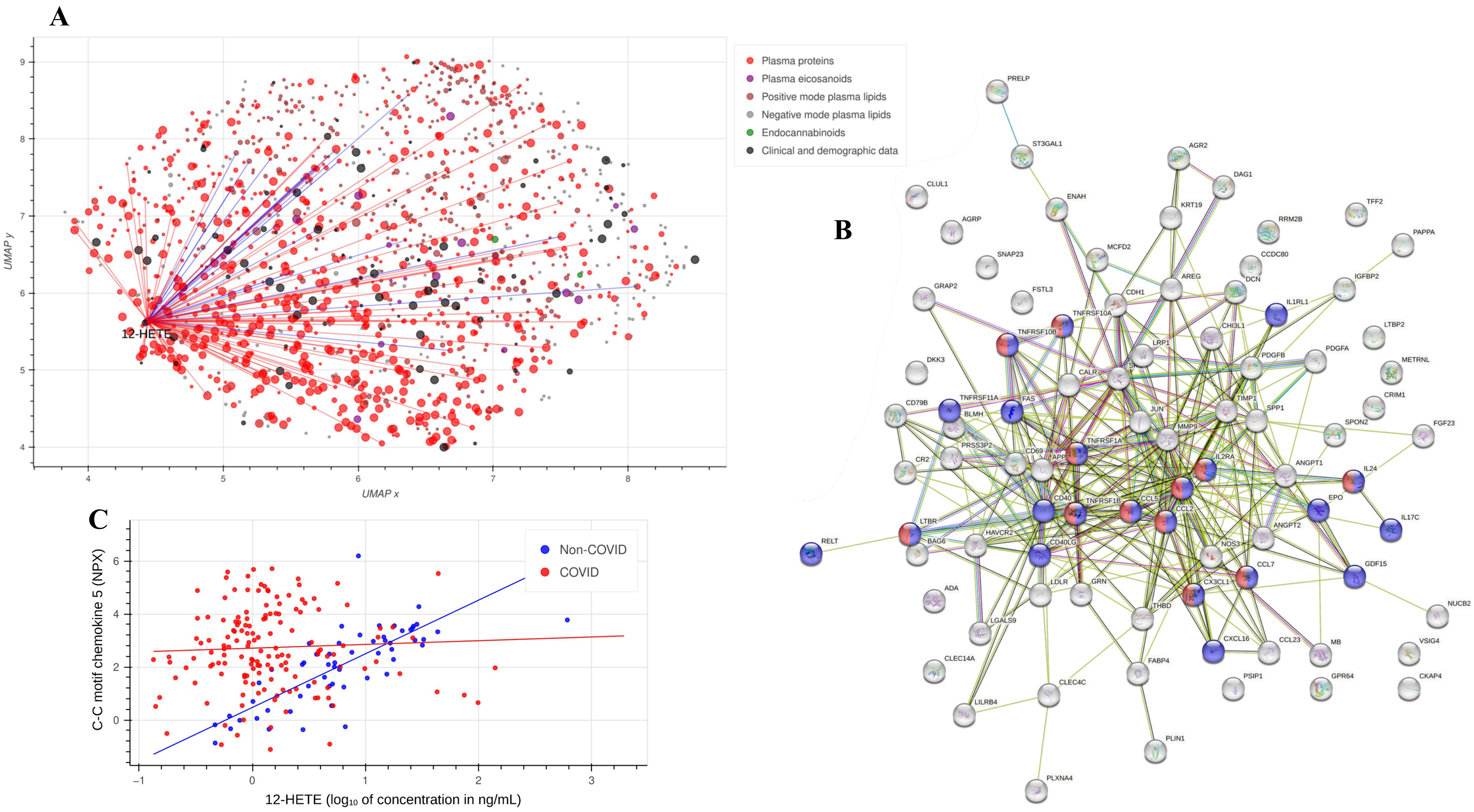
A
B
C
Plasma proteins
Plasma eicosanoids
Positive mode plasma lipids
Negative mode plasma lipids
Endocannabinoids
Clinical and demographic data
12-HETE
UMAP x
UMAP y
PRELP
ST3GAL1
AGR2
DAG1
CLUL1
ENAH
KRT19
TFF2
AGRP
RRM2B
PAPPA
MCFD2
SNAP23
CCDC80
AREG
DCN
IGFBP2
GRAP2
FSTL3
CDH1
TNFRSF10A
IL1RL1
TNFRSF10B
CHI3L1
LTBP2
LRP1
PDGFB
PDGFA
DKK3
METRNL
CALR
CD79B
TNFRSF11A
FAS
TIMP1
CRIM1
BLMH
JUN
SPP1
SPON2
MMP9
FGF23
PRSS3P2
TNFRSF1A
CR2
CD69
APP
IL2RA
ANGPT1
IL24
CD40
TNFRSF1B
CCL5
EPO
CCL2
IL17C
LTBR
HAVCR2
ANGPT2
RELT
BAG6
CD40LG
NOS3
GDF15
CCL7
ADA
LDLR
GRN
CX3CL1
NUCB2
THBD
LGALS9
CXCL16
CCL23
MB
VSIG4
CLEC14A
FABP4
PSIP1
GPR64
CKAP4
CLEC4C
LILRB4
PLIN1
PLXNA4
Non-COVID
COVID
C-C motif chemokine 5 (NPX)
12-HETE (log10 of concentration in ng/mL)

**Supplementary Figure 15: Analysis of non-COVID ICU control vs. COVID-19 differential correlation network.** (A) UMAP projection of features measured in both non-COVID ICU control and COVID-19 cohorts. Each feature is represented by a circle whose size is negatively proportional to $\log_{10}$ of the p-value for difference of the feature in non-COVID control vs. COVID-19. Therefore, bigger circles correspond to features that differ more significantly between non-COVID control and COVID-19 cases. Differential correlations of eicosanoid 12-HETE (measured in plasma) with other features are represented by lines, red lines correspond to the increase of Spearman's correlation coefficient in COVID-19 when compared to non-COVID control by 0.4 or more, blue lines correspond to the decrease of Spearman's correlation coefficient in COVID-19 by 0.4 or more. (B) STRING-DB enrichment analysis of 87 proteins differentially correlated with plasma 12-HETE (difference in correlation coefficients at least 0.4 with p-value for the difference at most 0.01). Proteins colored red belong to the *Viral protein interaction with cytokine and cytokine receptor* KEGG pathway while proteins colored blue belong to the *Cytokine-cytokine receptor interaction* KEGG pathway. FDR adjusted p-values for the two pathways were $8.57 \cdot 10^{-12}$ and $2.12 \cdot 10^{-18}$ respectively. (C) Loss of correlation between C-C motif chemokine (CCL5) and plasma 12-HETE in COVID-19, FDR adjusted p-value for difference in correlation is $2.46 \cdot 10^{-7}$. Theil-Sen estimator was used to fit lines.

**Supplementary Table 1: Characteristic COVID-19 patients.**

| Subject ID | Age | Sex | Race | Specimen collected | ICU | Time of collection |
|---|---|---|---|---|---|---|
| 5 | 19 | M | Unknown | Serum | N | Within 24 hr |
| 34 | 32 | F | Black | Serum | N | Within 24 hr |
| 42 | 30 | F | Black | Serum | N | Within 24 hr |
| 44 | 22 | M | Hispanic Latino/White | Serum | N | Within 24 hr |
| 49 | 38 | M | Black | Serum | N | Within 24 hr |
| 61 | 44 | M | Black | Serum | N | Within 24 hr |
| 62 | 29 | M | White | Serum | N | Within 24 hr |
| 69 | 43 | F | Black | Serum | N | Within 24 hr |
| 72 | 27 | F | Black | Serum | N | Within 24 hr |
| 73 | 30 | M | Hispanic Latino/White | Serum | N | Within 24 hr |
| 75 | 31 | M | Black | Serum | N | Within 24 hr |
| 10 | 27 | M | Unknown | Serum | Y | Within 24 hr |
| 39 | 41 | F | Black | Serum | Y | Within 24 hr |
| 51 | 19 | F | Unknown | Serum | Y | Within 24 hr |
| 64 | 27 | M | Black | Serum | Y | Within 24 hr |
| 71 | 40 | M | Black | Serum | Y | Within 24 hr |
| 77 | 30 | M | White | Serum | Y | Within 24 hr |
| 29 | 59 | F | Black | PBMCs | N | Within 24 hr |
| 38 | 61 | F | Hispanic Latino/White | PBMCs | N | Within 24 hr |
| 88 | 63 | F | Black | PBMCs | Y | Within 24 hr |
| 87 | 65 | F | Black | PBMCs | Y | Within 24 hr |
| 91 | 51 | F | Asian | PBMCs | Y | Within 24 hr |
| 103 | 73 | F | White | PBMCs | Y | Within 24 hr |
| 35 | 46 | M | Hispanic Latino/White | PBMCs | Y | Within 24 hr |
| 59 | 46 | M | Black | PBMCs | N | Within 24 hr |
| 76 | 60 | M | White | PBMCs | N | Within 24 hr |
| 82 | 58 | M | White | PBMCs | Y | Within 24 hr |
| 90 | 61 | M | Hispanic Latino/White | PBMCs | N | Within 24 hr |

**Supplementary Table 2: Characteristics of healthy individuals for PBMC study**

| Subject ID | Age | Sex | Race | Specimen collected |
|---|---|---|---|---|
| F1 | 75 | F | White | Plasma, Serum, PBMCs, urine |
| F2 | 64 | F | White | Plasma, Serum, PBMCs, urine |
| F3 | 59 | F | Hispanic Latino/White | Plasma, Serum, PBMCs, urine |
| F4 | 61 | F | Black | Plasma, Serum, PBMCs, urine |
| F5 | 37 | F | Black | Plasma, Serum, PBMCs, urine |
| F6 | 60 | F | Mix | Plasma, Serum, PBMCs, urine |
| F7 | 55 | F | Mix | Plasma, Serum, PBMCs, urine |
| F8 | 54 | F | White | Plasma, Serum, PBMCs, urine |
| M1 | 31 | M | White | Plasma, Serum, PBMCs, urine |
| M2 | 45 | M | Black | Plasma, Serum, PBMCs, urine |
| M3 | 29 | M | White | Plasma, Serum, PBMCs, urine |
| M4 | 48 | M | Asian | Plasma, Serum, PBMCs, urine |
| M5 | 62 | M | White | Plasma, Serum, PBMCs, urine |
| M6 | 69 | M | White | Plasma, Serum, PBMCs, urine |
| M7 | 51 | M | White | Plasma, Serum, PBMCs, urine |
| M8 | 58 | M | Hispanic Latino/White | Plasma, Serum, PBMCs, urine |

**Supplementary Table 3: Surface Antibody Staining Cocktail for PBMCs**

| Antibody | Company | Clone | Catalog |
|---|---|---|---|
| Brilliant Violet 421 anti-human PD1 | Biolegend | EH12.2H7 | 329920 |
| GhostDye Violet 510 | Tonbo | | 13-0870-T500 |
| Brilliant Violet 570 anti-human CD16 | Biolegend | 3G8 | 302036 |
| Super Bright 600 anti-human CD56 | ThermoFisher | TULY56 | 63-0566-42 |
| Brilliant Violet 650 anti-human CD45RA | Biolegend | HI100 | 304136 |
| Super Bright 702 anti-human CD8 | ThermoFisher | OKT8 | 67-0086-42 |
| Brilliant Violet 785 anti-human CD4 | Biolegend | OKT4 | 317442 |
| FITC anti-human CD88 | Biolegend | S5/1 | 344305 |
| FITC anti-human CD89 | Biolegend | A59 | 354114 |
| BB700 Mouse Anti-Human CD19 | BD | SJ25C1 | 566396 |
| PE anti-human CD39 | ThermoFisher | eBioA1 | 12-0399-41 |
| PE/Cyanine5 anti-human HLA-DR | Biolegend | L243 | 307608 |
| PE/Cy7 anti-human CD27 | BD | M-T271 | 560609 |
| Alexa Fluor647 Anti-Human CXCR5 | BD | RF8B2 | 558113 |
| AF700 anti-human CD3 | Biolegend | HIT3a | 300324 |
| APC-H7 Mouse anti-Human CD14 | BD | MφP9 | 560180 |
| Brilliant Stain Buffer | BD | | 566349 |
| Human TruStain FcX | Biolegend | | 422302 |

| | Healthy control | Mild COVID-19 | Severe COVID-19 | All COVID-19 | Non-COVID control |
|---|---|---|---|---|---|
| Plasma proteins | 0 | 76 | 91 | 167 | 57 |
| Plasma eicosanoids | 16 | 67 | 86 | 153 | 249 |
| Urine eicosanoids | 26 | 105 | 71 | 176 | 11 |
| Plasma lipids | 16 | 16 | 51 | 67 | 66 |
| Flow cytometry data | 0 | 60 | 60 | 120 | 2 |
| Mass cytometry data | 0 | 5 | 7 | 12 | 0 |
| Endocannabinoids | 16 | 6 | 8 | 14 | 48 |
| **Total samples** | **26** | **142** | **127** | **269** | **263** |

**Supplementary Table 4: Sample sizes for various analyses.** Each cell represents the number of assayed samples in a specific cohort. Urine eicosanoids were measured on 11 samples from non-COVID cohort, but no other measurements were performed on these samples, making them unsuitable for integration analysis.

**Supplementary Table 5**: Features signficantly correlated to sPLA2 levels in covid subjects

| Data type | Correlated feature | Spearmans's rho | p | n |
|---|---|---|---|---|
| Clinical and demographic data | ards_ever | 0.254401674 | 0.039271716 | 66 |
| | ast | 0.335493972 | 0.013137471 | 54 |
| | bmi | 0.261287966 | 0.034080357 | 66 |
| | crp | 0.676550552 | 0.0003931 | 23 |
| | daysaliveventfree_in28 | -0.381841573 | 0.00569387 | 51 |
| | ino_yn | 0.287568187 | 0.024626412 | 61 |
| | lymphocytes | -0.362081387 | 0.005645654 | 57 |
| | monocytes | -0.262306222 | 0.04870571 | 57 |
| | neutrophils_to_lymphocytes_ratio | 0.385283161 | 0.00336475 | 56 |
| | ordinal_d7 | 0.311160481 | 0.010988888 | 66 |
| | pct | 0.411440765 | 0.010277505 | 38 |
| | platelets_to_lymphocytes_ratio | 0.379835954 | 0.003884506 | 56 |
| | weight | 0.283566979 | 0.021039577 | 66 |
| Flow cytometry data | percOf_Bcell_BcellPB | 0.297624921 | 0.049750438 | 44 |

| | | | | |
|---|---|---|---|---|
| | percOf_Bcell_CD39pos | 0.296839212 | 0.050385528 | 44 |
| | percOf_Bcell_CD95pos | 0.303030303 | 0.045554579 | 44 |
| | percOf_Bcell_EOMESpos | 0.302406709 | 0.046023454 | 44 |
| | percOf_Bcell_KI67pos | 0.351258018 | 0.019389821 | 44 |
| | percOf_Bcell_TBETpos | 0.335318369 | 0.026076153 | 44 |
| | percOf_Bcell_TCF1pos | 0.340122631 | 0.023884288 | 44 |
| | percOf_BcellnotPB_TBETpos | 0.340662438 | 0.02364793 | 44 |
| | percOf_CD4_CD38pos | 0.388033405 | 0.009249681 | 44 |
| | percOf_CD4_CD4NNHLADRposCD38pos | 0.429799835 | 0.003594601 | 44 |
| | percOf_CD4_CD4NNKI67pos | 0.397603946 | 0.007525269 | 44 |
| | percOf_CD4_HLADRposCD38pos | 0.429120124 | 0.003653821 | 44 |
| | percOf_CD4_KI67pos | 0.393093728 | 0.008299796 | 44 |
| | percOf_CD4_TBETpos | 0.362508809 | 0.015594182 | 44 |
| | percOf_CD4acTfh_HLADRposCD38pos | 0.406074915 | 0.006238131 | 44 |
| | percOf_CD4CM_HLADRposCD38pos | 0.422510396 | 0.00427576 | 44 |
| | percOf_CD4CM_KI67pos | 0.405073996 | 0.006379519 | 44 |

| | | | | |
|---|---|---|---|---|
| | percOf_CD4cTfh_HLADRposCD38pos | 0.412544045 | 0.005388178 | 44 |
| | percOf_CD4cTfh_KI67pos | 0.369133192 | 0.013669244 | 44 |
| | percOf_CD4EM1_HLADRposCD38pos | 0.445399768 | 0.002447736 | 44 |
| | percOf_CD4EM1_KI67pos | 0.492547306 | 0.000682778 | 44 |
| | percOf_CD4EM2_HLADRposCD38pos | 0.350951374 | 0.019503293 | 44 |
| | percOf_CD4EM2_KI67pos | 0.314164905 | 0.037815452 | 44 |
| | percOf_CD4EM3_HLADRposCD38pos | 0.486821705 | 0.000805185 | 44 |
| | percOf_CD4EM3_KI67pos | 0.347145877 | 0.020958635 | 44 |
| | percOf_CD4nonNaive_CD38pos | 0.480620155 | 0.000959568 | 44 |
| | percOf_CD4nonNaive_HLADRposCD38pos | 0.437506607 | 0.002979712 | 44 |
| | percOf_CD4nonNaive_KI67pos | 0.413107822 | 0.005319122 | 44 |
| | percOf_CD4nonNaive_TBETpos | 0.355826492 | 0.017764023 | 44 |
| | percOf_CD8_CD38pos | 0.450051094 | 0.002175122 | 44 |
| | percOf_CD8_CD95pos | 0.331594898 | 0.027887978 | 44 |
| | percOf_CD8_HLADRpos | 0.340662438 | 0.02364793 | 44 |
| | percOf_CD8_HLADRposCD38pos | 0.414179499 | 0.005189976 | 44 |

| | | | | |
|---|---|---|---|---|
| | percOf_CD8_KI67pos | 0.420295983 | 0.004503865 | 44 |
| | percOf_CD8_TBETpos | 0.345172657 | 0.021748472 | 44 |
| | percOf_CD8CM_HLADRposCD38pos | 0.444115574 | 0.002528121 | 44 |
| | percOf_CD8CM_KI67pos | 0.52080764 | 0.000289645 | 44 |
| | percOf_CD8EM1_HLADRposCD38pos | 0.35503876 | 0.018035846 | 44 |
| | percOf_CD8EM2_HLADRposCD38pos | 0.429051464 | 0.00365985 | 44 |
| | percOf_CD8EM3_HLADRposCD38pos | 0.365186751 | 0.014790011 | 44 |
| | percOf_CD8EMRA_HLADRposCD38pos | 0.361746362 | 0.015829809 | 44 |
| | percOf_CD8Ex_HLADRposCD38pos | 0.375757576 | 0.011950362 | 44 |
| | percOf_CD8Ex_KI67pos | 0.516701903 | 0.000329609 | 44 |
| | percOf_CD8nonNaive_CD38pos | 0.389711064 | 0.008924906 | 44 |
| | percOf_CD8nonNaive_CD95pos | 0.387342308 | 0.009386396 | 44 |
| | percOf_CD8nonNaive_HLADRposCD38pos | 0.36124031 | 0.015987859 | 44 |
| | percOf_CD8RAposCD27posR7neg_HLADRposCD38pos | 0.4389161 | 0.00287786 | 44 |
| | percOf_CD8RAposCD27posR7neg_KI67pos | 0.303566397 | 0.045154602 | 44 |
| | percOf_CD8RAposCD27posR7posCD95pos_HLADRposCD38pos | 0.315796892 | 0.036777575 | 44 |

| | | | | |
|---|---|---|---|---|
| | percOf_CD8RAposCD27posR7posCD95pos_KI67pos | 0.360958421 | 0.016076477 | 44 |
| | percOf_Live_CD4EM1 | -0.32346723 | 0.032207893 | 44 |
| | percOf_Live_CD4HLADRposCD38pos | 0.402354363 | 0.006777804 | 44 |
| | percOf_Live_CD4NNHLADRposCD38pos | 0.410769669 | 0.005610632 | 44 |
| | percOf_Live_CD4nonNaive | -0.31516262 | 0.037178133 | 44 |
| | percOf_Live_CD8CM | -0.33894841 | 0.024405252 | 44 |
| | percOf_Live_CD8HLADRposCD38pos | 0.304027626 | 0.044812768 | 44 |
| | umap_component2 | 0.433121917 | 0.003317101 | 44 |
| Mass cytometry data | %CM in CD8 | 0.9 | 0.037386073 | 5 |
| | %DCs in Intact | 0.948683298 | 0.013846833 | 5 |
| | %EM in CD8 | 0.9 | 0.037386073 | 5 |
| | %NaÃ¯ve in CD4 | -1 | 1.40E-24 | 5 |
| | %pDCs in DCs | 0.894427191 | 0.040519326 | 5 |
| | %Th1-like in CD4 | 0.9 | 0.037386073 | 5 |
| | Early NK | 0.9 | 0.037386073 | 5 |
| | mDC | 0.9 | 0.03738607 | 5 |

| | | | | |
|---|---|---|---|---|
| | | | 3 | |
| | pDC | 0.894427191 | 0.040519326 | 5 |
| | Th1-like | 0.9 | 0.037386073 | 5 |
| | Treg | 0.9 | 0.037386073 | 5 |
| Negative mode plasma lipids | LPC 16:0.1 | -0.457400257 | 0.005035639 | 36 |
| | LPC 16:0.2 | -0.457400257 | 0.005035639 | 36 |
| | LPC 17:0 | -0.423680824 | 0.010025978 | 36 |
| | LPC 18:0 | -0.375289575 | 0.024107004 | 36 |
| | LPC 18:0 | -0.375289575 | 0.024107004 | 36 |
| | LPC 18:1 | -0.590733591 | 0.000148807 | 36 |
| | LPC 18:1 | -0.590733591 | 0.000148807 | 36 |
| | LPC 18:2 | -0.40978121 | 0.013064816 | 36 |
| | LPC 18:2 | -0.413642214 | 0.012151503 | 36 |
| | LPC 22:5 .1 | -0.381467181 | 0.021698633 | 36 |
| | LPC 22:5 .2 | -0.381467181 | 0.021698633 | 36 |

| | | | | |
|---|---|---|---|---|
| | LPE 18:0 .1 | -0.434603516 | 0.008080801 | 36 |
| | LPE 18:0 .2 | -0.434603516 | 0.008080801 | 36 |
| | PC 36:3.2 | -0.32998713 | 0.04935705 | 36 |
| | PC 37:6 | -0.461003861 | 0.004659197 | 36 |
| | PC 39:6 | -0.4996139 | 0.001919866 | 36 |
| | PC O-34:2.1 | -0.372972973 | 0.025065379 | 36 |
| | PC O-34:2.2 | -0.372972973 | 0.025065379 | 36 |
| | PC O-36:3.1 | -0.462548263 | 0.004505467 | 36 |
| | PC O-36:3.2 | -0.462548263 | 0.004505467 | 36 |
| | PC O-36:3.3 | -0.462548263 | 0.004505467 | 36 |
| | PC O-38:6 | -0.437323037 | 0.007650053 | 36 |
| | PC O-40:5 | -0.33024453 | 0.049169412 | 36 |
| | PC O-40:6 | -0.455083655 | 0.005291258 | 36 |
| | PE 40:5 | -0.349292149 | 0.036791828 | 36 |
| | PE O-38:4 | - | 0.015 | 36 |

| | | | | |
|---|---|---|---|---|
| | | 0.402059202 | 066094 | |
| | PI 34:1 | -0.364736165 | 0.028731468 | 36 |
| | PI 36:1 | -0.449420849 | 0.005963673 | 36 |
| | PI 36:3 | -0.431660232 | 0.008570181 | 36 |
| | PI 36:4 | -0.441698842 | 0.00699821 | 36 |
| | PI 38:3 | -0.407979408 | 0.013510469 | 36 |
| | PI 38:4 | -0.353153153 | 0.034622226 | 36 |
| | PI 38:5 | -0.384555985 | 0.020571568 | 36 |
| Plasma eicosanoids | LTE4 | 0.302203877 | 0.013656782 | 66 |
| Plasma proteins | CAM_O00533 Neural cell adhesion molecule L1-like protein (CHL1) | -0.411877675 | 0.000590261 | 66 |
| | CAM_O15031 Plexin-B2 (PLXNB2) | 0.263458929 | 0.032567626 | 66 |
| | CAM_O95445 Apolipoprotein M (APOM) | -0.333472498 | 0.006215763 | 66 |
| | CAM_P01033 Metalloproteinase inhibitor 1 (TIMP1) | 0.405573531 | 0.000729138 | 66 |
| | CAM_P05154 Plasma serine protease inhibitor (SERPINA5) | -0.374637303 | 0.001940925 | 66 |
| | CAM_P06681 Complement C2 (C2) | 0.401774345 | 0.00082652 | 66 |

| | | | | |
|---|---|---|---|---|
| | CAM_P07478 Trypsin-2 (PRSS2) | 0.346874022 | 0.004326191 | 66 |
| | CAM_P10721 Mast/stem cell growth factor receptor Kit (KIT) | -0.242542532 | 0.049741186 | 66 |
| | CAM_P14543 Nidogen-1 (NID1) | 0.528483457 | 5.09045E-06 | 66 |
| | CAM_P15907 Beta-galactoside alpha-2,6-sialyltransferase 1 (ST6GAL1) | 0.574908673 | 4.45E-07 | 66 |
| | CAM_P20062 Transcobalamin-2 (TCN2) | 0.347041019 | 0.004306276 | 66 |
| | CAM_P22749 Granulysin (GNLY) | 0.465149776 | 8.31645E-05 | 66 |
| | CAM_P23141 Liver carboxylesterase 1 (CES1) | 0.349838221 | 0.003984538 | 66 |
| | CAM_P42785 Lysosomal Pro-X carboxypeptidase (PRCP) | 0.282496608 | 0.021550941 | 66 |
| | CAM_Q14767 Latent-transforming growth factor beta-binding protein 2 (LTBP2) | 0.29360192 | 0.016727166 | 66 |
| | CAM_Q15582 Transforming growth factor-beta-induced protein ig-h3 (TGFBI) | 0.412086421 | 0.000586104 | 66 |
| | CAM_Q8N423 Leukocyte immunoglobulin-like receptor subfamily B member 2 (LILRB2) | 0.275399228 | 0.025216034 | 66 |
| | CAM_Q8NHL6 Leukocyte immunoglobulin-like receptor subfamily B member 1 (LILRB1) | 0.298236092 | 0.015006699 | 66 |
| | CAM_Q99650 Oncostatin-M-specific receptor subunit beta (OSMR) | 0.271266047 | 0.027583594 | 66 |
| | CAM_Q9BXJ1 Complement C1q tumor necrosis factor-related protein 1 (C1QTNF1) | 0.326876109 | 0.007387873 | 66 |
| | CAM_Q9BXR6 Complement factor H-related protein 5 (CFHR5) | 0.35706085 | 0.00325017 | 66 |
| | CAM_Q9Y5C1 Angiopoietin-related protein 3 (ANGPTL3) | 0.294687402 | 0.016309704 | 66 |

| | | | | |
|---|---|---|---|---|
| | CVD2_O00182 Galectin-9 (Gal-9) | 0.295188394 | 0.016120063 | 66 |
| | CVD2_O00220 Tumor necrosis factor receptor superfamily member 10A (TNFRSF10A) | 0.25331385 | 0.040148845 | 66 |
| | CVD2_P00797 Renin (REN) | 0.279490659 | 0.023043845 | 66 |
| | CVD2_P01730 T-cell surface glycoprotein CD4 (CD4) | 0.282997599 | 0.021310295 | 66 |
| | CVD2_P04792 Heat shock 27 kDa protein (HSP 27) | 0.364366976 | 0.002632304 | 66 |
| | CVD2_P05231 Interleukin-6 (IL6) | 0.396680931 | 0.000975542 | 66 |
| | CVD2_P07711 Cathepsin L1 (CTSL1) | 0.417054587 | 0.000494695 | 66 |
| | CVD2_P09341 C-X-C motif chemokine 1 (CXCL1) | 0.455839683 | 0.000119937 | 66 |
| | CVD2_P12931 Proto-oncogene tyrosine-protein kinase Src (SRC) | 0.342866089 | 0.004829275 | 66 |
| | CVD2_P18510 Interleukin-1 receptor antagonist protein (IL-1ra) | 0.423275232 | 0.000398593 | 66 |
| | CVD2_P19883 Follistatin (FS) | 0.298653585 | 0.014859458 | 66 |
| | CVD2_P21583 Stem cell factor (SCF) | -0.326124622 | 0.007532966 | 66 |
| | CVD2_P26022 Pentraxin-related protein PTX3 (PTX3) | 0.319653481 | 0.008889121 | 66 |
| | CVD2_P35218 Carbonic anhydrase 5A, mitochondrial (CA5A) | 0.387704832 | 0.001298317 | 66 |
| | CVD2_Q13043 Serine/threonine-protein kinase 4 (STK4) | 0.281077132 | 0.022245351 | 66 |
| | CVD2_Q16698 2,4-dienoyl-CoA reductase, | 0.3301743 | 0.00677962 | 66 |

| | | | | |
|---|---|---|---|---|
| | mitochondrial (DECR1) | 03 | 6 | |
| | CVD2_Q99523 Sortilin (SORT1) | 0.2898862 33 | 0.018 22622 2 | 66 |
| | CVD2_Q9BQR3 Serine protease 27 (PRSS27) | - 0.3740945 62 | 0.001 97291 6 | 66 |
| | CVD2_Q9BUD6 Spondin-2 (SPON2) | 0.2750652 33 | 0.025 40075 1 | 66 |
| | CVD2_Q9BWV1 Brother of CDO (Protein BOC) | - 0.2871307 8 | 0.019 41072 3 | 66 |
| | CVD2_Q9UJM8 Hydroxyacid oxidase 1 (HAOX1) | 0.3497547 23 | 0.003 99382 5 | 66 |
| | CVD2_Q9UKP3 Melusin (ITGB1BP2) | 0.4257801 9 | 0.000 36495 7 | 66 |
| | CVD2_Q9Y6K9 NF-kappa-B essential modulator (NEMO) | 0.3705041 23 | 0.002 19672 2 | 66 |
| | CVD3_O00300 Osteoprotegerin (OPG) | 0.2834353 15 | 0.022 14175 | 65 |
| | CVD3_P00749 Urokinase-type plasminogen activator (uPA) | 0.4037150 35 | 0.000 85341 2 | 65 |
| | CVD3_P01130 Low-density lipoprotein receptor (LDL receptor) | 0.2895541 96 | 0.019 30833 8 | 65 |
| | CVD3_P04275 von Willebrand factor (vWF) | 0.2571241 26 | 0.038 66993 4 | 65 |
| | CVD3_P05121 Plasminogen activator inhibitor 1 (PAI) | 0.3820367 13 | 0.001 68741 5 | 65 |
| | CVD3_P05164 Myeloperoxidase (MPO) | 0.3772727 27 | 0.001 94829 3 | 65 |
| | CVD3_P10451 Osteopontin (OPN) | 0.3648601 4 | 0.002 80560 2 | 65 |
| | CVD3_P10646 Tissue factor pathway inhibitor (TFPI) | 0.2613636 36 | 0.035 46575 1 | 65 |
| | CVD3_P13500 Monocyte chemotactic protein 1 | 0.4062062 | 0.000 | 65 |

| | | | | |
|---|---|---|---|---|
| | (MCP-1) | 94 | 786763 | |
| | CVD3_P15085 Carboxypeptidase A1 (CPA1) | 0.35034965 | 0.00422192 | 65 |
| | CVD3_P20160 Azurocidin (AZU1 | 0.308566434 | 0.012390502 | 65 |
| | CVD3_P24158 Myeloblastin (PRTN3) | 0.495148601 | 2.75043E-05 | 65 |
| | CVD3_P28799 Granulins (GRN) | 0.447333916 | 0.000187 | 65 |
| | CVD3_P42574 Caspase-3 (CASP-3) | 0.336013986 | 0.0062092 | 65 |
| | CVD3_Q12860 Contactin-1 (CNTN1) | -0.39375 | 0.001174157 | 65 |
| | CVD3_Q13867 Bleomycin hydrolase (BLM hydrolase) | 0.389816434 | 0.00132818 | 65 |
| | CVD3_Q15166 Paraoxonase (PON3) | -0.277229021 | 0.025368573 | 65 |
| | CVD3_Q8NBP7 Proprotein convertase subtilisin/kexin type 9 (PCSK9) | 0.421284965 | 0.000474459 | 65 |
| | CVD3_Q92876 Kallikrein-6 (KLK6 ) | -0.384090909 | 0.001584946 | 65 |
| | CVD3_Q99969 Retinoic acid receptor responder protein 2 (RARRES2) | 0.248208042 | 0.046195164 | 65 |
| | CVD3_Q99988 Growth/differentiation factor 15 (GDF-15) | 0.29527972 | 0.01694311 | 65 |
| | CVD3_Q9H2A7 C-X-C motif chemokine 16 (CXCL16) | 0.298513986 | 0.015720132 | 65 |
| | CVD3_Q9Y275 Tumor necrosis factor ligand superfamily member 13B (TNFSF13B) | 0.379326923 | 0.001831671 | 65 |
| | DEV_O00585 C-C motif chemokine 21 (CCL21) | 0.281870368 | 0.021854994 | 66 |
| | DEV_O14773 Tripeptidyl-peptidase 1 (TPP1) | 0.422356748 | 0.000411617 | 66 |

| | | | | |
|---|---|---|---|---|
| | DEV_O43464 Serine protease HTRA2, mitochondrial (HTRA2) | 0.356601607 | 0.003293 | 66 |
| | DEV_O95721 Synaptosomal-associated protein 29 (SNAP29) | 0.397056675 | 0.000963771 | 66 |
| | DEV_P00995 Serine protease inhibitor Kazal-type 1 (SPINK1) | 0.299154577 | 0.014684411 | 66 |
| | DEV_P04233 HLA class II histocompatibility antigen gamma chain (CD74) | 0.314727064 | 0.010059785 | 66 |
| | DEV_P07237 Protein disulfide-isomerase (P4HB) | 0.421939255 | 0.000417665 | 66 |
| | DEV_P14174 Macrophage migration inhibitory factor (MIF) | 0.327711095 | 0.007229534 | 66 |
| | DEV_P15289 Arylsulfatase A (ARSA) | 0.386911596 | 0.001331022 | 66 |
| | DEV_P15291 Beta-1,4-galactosyltransferase 1 (B4GALT1) | 0.444233379 | 0.000186589 | 66 |
| | DEV_P19256 Lymphocyte function-associated antigen 3 (CD58) | -0.28708903 | 0.019429163 | 66 |
| | DEV_P23280 Carbonic anhydrase 6 (CA6) | -0.325665379 | 0.007622857 | 66 |
| | DEV_P23284 Peptidyl-prolyl cis-trans isomerase B (PPIB) | 0.384114393 | 0.001452354 | 66 |
| | DEV_P30086 Phosphatidylethanolamine-binding protein 1 (PEBP1) | 0.249347667 | 0.04348533 | 66 |
| | DEV_P31948 Stress-induced-phosphoprotein 1 (STIP1) | 0.323035174 | 0.008155988 | 66 |
| | DEV_P50895 Basal cell adhesion molecule (BCAM) | 0.293894166 | 0.016613881 | 66 |
| | DEV_P55103 Inhibin beta C chain (INHBC) | 0.320237971 | 0.008758443 | 66 |
| | DEV_P63313 Thymosin beta-10 (TMSB10) | 0.262874439 | 0.032969268 | 66 |

| | | | | |
|---|---|---|---|---|
| | DEV_P78552 Interleukin-13 receptor subunit alpha-1 (IL13RA1) | 0.432668824 | 0.000285374 | 66 |
| | DEV_Q07108 Early activation antigen CD69 (CD69) | 0.390710782 | 0.001180851 | 66 |
| | DEV_Q11128,P21217 Galactoside 3(4)-L-fucosyltransferase,Alpha-(1,3)-fucosyltransferase 3/5 (FUT3/5) | 0.332345267 | 0.006403657 | 66 |
| | DEV_Q14112 Nidogen-2 (NID2) | 0.301116794 | 0.014015786 | 66 |
| | DEV_Q14118 Dystroglycan (DAG1) | 0.288091013 | 0.018990698 | 66 |
| | DEV_Q14696 LDLR chaperone MESD (MESDC2) | 0.350088717 | 0.003956793 | 66 |
| | DEV_Q4KMG0 Cell adhesion molecule-related/down-regulated by oncogenes (CDON) | -0.305082977 | 0.01274404 | 66 |
| | DEV_Q8IWV2 Contactin-4 (CNTN4) | -0.254399332 | 0.039273587 | 66 |
| | DEV_Q8TEU8 WAP, Kazal, immunoglobulin, Kunitz and NTR domain-containing protein 2 (WFIKKN2) | -0.279240163 | 0.023172118 | 66 |
| | DEV_Q96QR1 Secretoglobin family 3A member 1 (SCGB3A1) | -0.359565807 | 0.003025155 | 66 |
| | DEV_Q99497 Protein deglycase DJ-1 (PARK7) | 0.310426887 | 0.011188959 | 66 |
| | DEV_Q99538 Legumain (LGMN) | 0.263709425 | 0.03239675 | 66 |
| | DEV_Q9BY76 Angiopoietin-related protein 4 (ANGPTL4) | 0.268301847 | 0.029394509 | 66 |
| | DEV_Q9NQ38 Serine protease inhibitor Kazal-type 5 (SPINK5) | -0.300949796 | 0.01407165 | 66 |
| | DEV_Q9NZV1 Cysteine-rich motor neuron 1 protein (CRIM1) | 0.327961591 | 0.007182616 | 66 |
| | DEV_Q9UBX1 Cathepsin F (CTSF) | - | 0.002 | 66 |

| | | | | |
|---|---|---|---|---|
| | | 0.365786452 | 525223 | |
| | DEV_Q9UKK9 ADP-sugar pyrophosphatase (NUDT5) | 0.382277424 | 0.001537358 | 66 |
| | DEV_Q9Y240 C-type lectin domain family 11 member A (CLEC11A) | 0.372382841 | 0.002076931 | 66 |
| | DEV_Q9Y279 V-set and immunoglobulin domain-containing protein 4 (VSIG4) | 0.27468949 | 0.025609922 | 66 |
| | INF_O00300 Osteoprotegerin (OPG) | 0.342949588 | 0.004818289 | 66 |
| | INF_O14625 C-X-C motif chemokine 11 (CXCL11) | 0.388581568 | 0.001263013 | 66 |
| | INF_O14788 TNF-related activation-induced cytokine (TRANCE) | -0.327168354 | 0.007332113 | 66 |
| | INF_O15169 Axin-1 (AXIN1) | 0.342156351 | 0.004923558 | 66 |
| | INF_O43557 Tumor necrosis factor ligand superfamily member 14 (TNFSF14 ) | 0.368583655 | 0.002325528 | 66 |
| | INF_O95630 STAM-binding protein (STAMPB) | 0.261204467 | 0.034139691 | 66 |
| | INF_P00749 Urokinase-type plasminogen activator (uPA) | 0.396847928 | 0.000970295 | 66 |
| | INF_P00813 Adenosine Deaminase (ADA) | 0.295063146 | 0.016167295 | 66 |
| | INF_P01137 Latency-associated peptide transforming growth factor beta-1 (LAP TGF-beta-1) | 0.300407056 | 0.014254537 | 66 |
| | INF_P01579 Interferon gamma (IFN-gamma) | 0.27468949 | 0.025609922 | 66 |
| | INF_P02778 C-X-C motif chemokine 10 (CXCL10 ) | 0.452374491 | 0.000137081 | 66 |
| | INF_P05231 Interleukin-6 (IL6) | 0.424360714 | 0.00038368 | 66 |

| | | | | |
|---|---|---|---|---|
| | | | 5 | |
| | INF_P09341 C-X-C motif chemokine 1 (CXCL1) | 0.4477403 19 | 0.000 16353 7 | 66 |
| | INF_P09603 Macrophage colony-stimulating factor 1 (CSF-1) | 0.4290783 84 | 0.000 32462 2 | 66 |
| | INF_P10145 Interleukin-8 (IL-8) | 0.4652750 23 | 8.274 99E-05 | 66 |
| | INF_P13232 Interleukin-7 (IL-7) | 0.4025675 82 | 0.000 80526 5 | 66 |
| | INF_P13500 Monocyte chemotactic protein 1 (MCP-1) | 0.4422711 62 | 0.000 20075 3 | 66 |
| | INF_P13725 Oncostatin-M (OSM) | 0.2915979 54 | 0.017 52195 7 | 66 |
| | INF_P14210 Hepatocyte growth factor (HGF) | 0.3231604 22 | 0.008 12988 1 | 66 |
| | INF_P15018 Leukemia inhibitory factor (LIF) | 0.2769857 01 | 0.024 35405 6 | 66 |
| | INF_P21583 Stem cell factor (SCF) | - 0.2723932 78 | 0.026 92004 6 | 66 |
| | INF_P22301 Interleukin-10 (IL10) | 0.4745851 16 | 5.675 98E-05 | 66 |
| | INF_P42830 C-X-C motif chemokine 5 (CXCL5 ) | 0.2477611 94 | 0.044 88244 | 66 |
| | INF_P50225 Sulfotransferase 1A1 (ST1A1) | 0.2848763 18 | 0.020 42811 5 | 66 |
| | INF_P55773 C-C motif chemokine 23 (CCL23) | 0.3524684 27 | 0.003 70163 5 | 66 |
| | INF_P78556 C-C motif chemokine 20 (CCL20) | 0.3458720 38 | 0.004 44740 7 | 66 |
| | INF_P80075 Monocyte chemotactic protein 2 (MCP-2) | 0.4418536 69 | 0.000 20389 1 | 66 |
| | INF_P80098 Monocyte chemotactic protein 3 | 0.5497338 | 1.746 | 66 |

| | | | | |
|---|---|---|---|---|
| | (MCP-3) | 48 | 48E-06 | |
| | INF_P80162 C-X-C motif chemokine 6 (CXCL6) | 0.31731552 | 0.009428985 | 66 |
| | INF_P80511 Protein S100-A12 (EN-RAGE ) | 0.431207598 | 0.000300791 | 66 |
| | INF_Q13478 Interleukin-18 receptor 1 (IL-18R1) | 0.534328358 | 3.82041E-06 | 66 |
| | INF_Q14790 Caspase-8 (CASP-8 ) | 0.377392756 | 0.001785568 | 66 |
| | INF_Q8IXJ6 SIR2-like protein 2 (SIRT2) | 0.276526459 | 0.024600973 | 66 |
| | INF_Q8NFT8 Delta and Notch-like epidermal growth factor-related receptor (DNER) | -0.389374804 | 0.00123182 | 66 |
| | INF_Q99731 C-C motif chemokine 19 (CCL19) | 0.27761194 | 0.024020733 | 66 |
| | INF_Q9H5V8 CUB domain-containing protein 1 (CDCP1) | 0.285293811 | 0.020236355 | 66 |
| | INF_Q9NZQ7 Programmed cell death 1 ligand 1 (PD-L1) | 0.284375326 | 0.020660267 | 66 |
| | IRE_O00273 DNA fragmentation factor subunit alpha (DFFA) | 0.42160526 | 0.000422561 | 66 |
| | IRE_O14867 Transcription regulator protein BACH1 (BACH1) | 0.474919111 | 5.5986E-05 | 66 |
| | IRE_O43597 Protein sprouty homolog 2 (SPRY2) | 0.341237867 | 0.005047981 | 66 |
| | IRE_O75475 PC4 and SFRS1-interacting protein (PSIP1) | 0.402818077 | 0.000798657 | 66 |
| | IRE_O94992 Protein HEXIM1 (HEXIM1) | 0.458010646 | 0.000110225 | 66 |
| | IRE_O95786 Probable ATP-dependent RNA helicase DDX58 (DDX58) | 0.391044776 | 0.001168409 | 66 |

| | | | | |
|---|---|---|---|---|
| | IRE_P05231 Interleukin-6 (IL6) | 0.445986849 | 0.000174714 | 66 |
| | IRE_P08727 Keratin, type I cytoskeletal 19 (KRT19) | 0.338816407 | 0.005389403 | 66 |
| | IRE_P09038 Fibroblast growth factor 2 (FGF2) | 0.288132763 | 0.018972613 | 66 |
| | IRE_P14317 Hematopoietic lineage cell-specific protein (HCLS1) | 0.39676443 | 0.000972915 | 66 |
| | IRE_P16278 Beta-galactosidase (GLB1) | 0.445819852 | 0.000175815 | 66 |
| | IRE_P16455 Methylated-DNA--protein-cysteine methyltransferase (MGMT) | 0.336603695 | 0.005719024 | 66 |
| | IRE_P18627 Lymphocyte activation gene 3 protein (LAG3) | 0.310468636 | 0.011177489 | 66 |
| | IRE_P19474 E3 ubiquitin-protein ligase TRIM21 (TRIM21) | 0.429579376 | 0.000318864 | 66 |
| | IRE_P22301 Interleukin-10 (IL10) | 0.496670494 | 2.21909E-05 | 66 |
| | IRE_P30044 Peroxiredoxin-5, mitochondrial (PRDX5) | 0.406700762 | 0.000702316 | 66 |
| | IRE_P30048 Thioredoxin-dependent peroxide reductase, mitochondrial (PRDX3) | 0.463438054 | 8.90263E-05 | 66 |
| | IRE_P50135 Histamine N-methyltransferase (HNMT) | 0.253981839 | 0.039608334 | 66 |
| | IRE_P51617 Interleukin-1 receptor-associated kinase 1 (IRAK1) | 0.425320948 | 0.000370923 | 66 |
| | IRE_P78362 SRSF protein kinase 2 (SRPK2) | 0.406492015 | 0.000707214 | 66 |
| | IRE_P78410 Butyrophilin subfamily 3 member A2 (BTN3A2) | 0.412879658 | 0.000570549 | 66 |
| | IRE_Q00978 Interferon regulatory factor 9 (IRF9) | 0.454503705 | 0.00012629 | 66 |

| | | | | |
|---|---|---|---|---|
| | | | 8 | |
| | IRE_Q04637 Eukaryotic translation initiation factor 4 gamma 1 (EIF4G1) | 0.4130884 04 | 0.000 56651 9 | 66 |
| | IRE_Q05084 Islet cell autoantigen 1 (ICA1) | 0.4230247 36 | 0.000 40210 7 | 66 |
| | IRE_Q05516 Zinc finger and BTB domain-containing protein 16 (ZBTB16) | 0.3987266 46 | 0.000 91299 3 | 66 |
| | IRE_Q06830 Peroxiredoxin-1 (PRDX1) | 0.3397766 41 | 0.005 25164 9 | 66 |
| | IRE_Q07065 Cytoskeleton-associated protein 4 (CKAP4) | 0.3820686 78 | 0.001 54729 5 | 66 |
| | IRE_Q12968 Nuclear factor of activated T-cells, cytoplasmic 3 (NFATC3) | 0.3018682 81 | 0.013 76675 5 | 66 |
| | IRE_Q13490 Baculoviral IAP repeat-containing protein 2 (BIRC2) | 0.4211042 69 | 0.000 43000 4 | 66 |
| | IRE_Q14203 Dynactin subunit 1 (DCTN1) | 0.3425738 44 | 0.004 86790 2 | 66 |
| | IRE_Q8IU57 Interferon lambda receptor 1 (IFNLR1) | 0.3952614 55 | 0.001 02119 3 | 66 |
| | IRE_Q8NHJ6 Leukocyte immunoglobulin-like receptor subfamily B member 4 (LILRB4) | 0.4183070 66 | 0.000 4738 | 66 |
| | IRE_Q96DB9 FXYD domain-containing ion transport regulator 5 (FXYD5) | 0.3097171 49 | 0.011 38552 3 | 66 |
| | IRE_Q96SB3 Neurabin-2 (PPP1R9B) | 0.3392756 5 | 0.005 32312 8 | 66 |
| | IRE_Q9C035 Tripartite motif-containing protein 5 (TRIM5) | 0.5104060 12 | 1.196 59E-05 | 66 |
| | IRE_Q9GZT9 Egl nine homolog 1 (EGLN1) | 0.2525206 14 | 0.040 79862 | 66 |
| | IRE_Q9HCM2 Plexin-A4 (PLXNA4) | 0.2741467 49 | 0.025 91462 5 | 66 |
| | IRE_Q9NWZ3 Interleukin-1 receptor-associated kinase 4 (IRAK4) | 0.2681348 5 | 0.029 49943 | 66 |

| | | | | |
|---|---|---|---|---|
| | | | 5 | |
| | IRE_Q9UKX5 Integrin alpha-11 (ITGA11) | -0.251059388 | 0.042018348 | 66 |
| | IRE_Q9UQQ2 SH2B adapter protein 3 (SH2B3) | 0.288174512 | 0.018954543 | 66 |
| | IRE_Q9UQV4 Lysosome-associated membrane glycoprotein 3 (LAMP3) | 0.333889991 | 0.006147413 | 66 |
| | IRE_Q9Y2J8 Protein-arginine deiminase type-2 (PADI2) | 0.275440977 | 0.025193024 | 66 |
| | IRE_Q9Y3P8 Signaling threshold-regulating transmembrane adapter 1 (SIT1) | -0.364032982 | 0.002658081 | 66 |
| | MET_O00161 Synaptosomal-associated protein 23 (SNAP23) | 0.269512577 | 0.028643127 | 66 |
| | MET_O15123 Angiopoietin-2 (ANGPT2) | 0.399185889 | 0.00089946 | 66 |
| | MET_O75356 Ectonucleoside triphosphate diphosphohydrolase 5 (ENTPD5) | 0.383362906 | 0.001486604 | 66 |
| | MET_O75791 GRB2-related adapter protein 2 (GRAP2) | 0.36950214 | 0.002263106 | 66 |
| | MET_O95544 NAD kinase (NADK) | 0.490700344 | 2.8788E-05 | 66 |
| | MET_O95841 Angiopoietin-related protein 1 (ANGPTL1) | 0.344744807 | 0.004587358 | 66 |
| | MET_P09104 Gamma-enolase (ENO2) | 0.294102912 | 0.016533367 | 66 |
| | MET_P09417 Dihydropteridine reductase (QDPR) | 0.45717566 | 0.000113871 | 66 |
| | MET_P09668 Pro-cathepsin H (CTSH) | 0.274522492 | 0.025703353 | 66 |
| | MET_P12724 Eosinophil cationic protein (RNASE3) | 0.416804091 | 0.000498974 | 66 |
| | MET_P16083 Ribosyldihydronicotinamide | 0.2589917 | 0.03574371 | 66 |

| | | | | |
|---|---|---|---|---|
| | dehydrogenase [quinone] (NQO2) | 55 | 9 | |
| | MET_P19022 Cadherin-2 (CDH2) | 0.3585220 75 | 0.003 11716 8 | 66 |
| | MET_P19971 Thymidine phosphorylase (TYMP) | 0.4679195 8 | 8.482 05E-05 | 65 |
| | MET_P21964 Catechol O-methyltransferase (COMT) | 0.4450683 64 | 0.000 18084 5 | 66 |
| | MET_P23526 Adenosylhomocysteinase (AHCY) | 0.2921824 44 | 0.017 28687 2 | 66 |
| | MET_P27695 DNA-(apurinic or apyrimidinic site) lyase (APEX1) | 0.4377622 38 | 0.000 23711 2 | 66 |
| | MET_P31431 Syndecan-4 (SDC4) | 0.2753992 28 | 0.025 21603 4 | 66 |
| | MET_P35754 Glutaredoxin-1 (GLRX) | 0.3526771 74 | 0.003 67996 5 | 66 |
| | MET_P41236 Protein phosphatase inhibitor 2 (PPP1R2) | 0.4550464 46 | 0.000 12367 7 | 66 |
| | MET_P43234 Cathepsin O (CTSO) | 0.4273249 14 | 0.000 34552 8 | 66 |
| | MET_P46109 Crk-like protein (CRKL) | 0.4275754 1 | 0.000 34246 8 | 66 |
| | MET_P50452 Serpin B8 (SERPINB8) | 0.4089552 24 | 0.000 65133 9 | 66 |
| | MET_P51693 Amyloid-like protein 1 (APLP1) | - 0.3261246 22 | 0.007 53296 6 | 66 |
| | MET_P52888 Thimet oligopeptidase (THOP1) | 0.3225759 32 | 0.008 25234 2 | 66 |
| | MET_P98082 Disabled homolog 2 (DAB2) | 0.4286191 42 | 0.000 32998 2 | 66 |
| | MET_Q02790 Peptidyl-prolyl cis-trans isomerase FKBP4 (FKBP4) | 0.4178060 75 | 0.000 48206 | 66 |
| | MET_Q13275 Semaphorin-3F (SEMA3F) | 0.3038304 | 0.013 | 66 |

| | | | | |
|---|---|---|---|---|
| | | 98 | 134415 | |
| | MET_Q15155 Nodal modulator 1 (NOMO1) | 0.339117133 | 0.005720281 | 65 |
| | MET_Q16773 Kynurenine--oxoglutarate transaminase 1 (KYAT1) | 0.331051039 | 0.006625522 | 66 |
| | MET_Q76M96 Coiled-coil domain-containing protein 80 (CCDC80) | 0.248136938 | 0.04454825 | 66 |
| | MET_Q8N1Q1 Carbonic anhydrase 13 (CA13) | 0.353929652 | 0.003552296 | 66 |
| | MET_Q8NI22 Multiple coagulation factor deficiency protein 2 (MCFD2) | 0.434756288 | 0.000264601 | 66 |
| | MET_Q8WVQ1 Soluble calcium-activated nucleotidase 1 (CANT1) | 0.296190377 | 0.015746453 | 66 |
| | MET_Q96JA1 Leucine-rich repeats and immunoglobulin-like domains protein 1 (LRIG1) | 0.490074105 | 2.95765E-05 | 66 |
| | MET_Q9GZM7 Tubulointerstitial nephritis antigen-like (TINAGL1) | 0.336645444 | 0.005712644 | 66 |
| | MET_Q9NY25 C-type lectin domain family 5 member A (CLEC5A) | 0.271934036 | 0.027188738 | 66 |
| | ODA_O15357 Phosphatidylinositol 3,4,5-trisphosphate 5-phosphatase 2 (INPPL1) | 0.395303204 | 0.001019823 | 66 |
| | ODA_O60240 Perilipin-1 (PLIN1) | 0.28771527 | 0.019154122 | 66 |
| | ODA_O60934 Nibrin (NBN) | 0.373468323 | 0.002010416 | 66 |
| | ODA_O75354 Ectonucleoside triphosphate diphosphohydrolase 6 (ENTPD6) | 0.288592005 | 0.018774651 | 66 |
| | ODA_O95994 Anterior gradient protein 2 homolog (AGR2) | 0.250516648 | 0.042478996 | 66 |
| | ODA_P01258 Calcitonin (CALCA) | 0.316188289 | 0.009699324 | 66 |

| | | | | |
|---|---|---|---|---|
| | ODA_P07947 Tyrosine-protein kinase Yes (YES1) | 0.349963469 | 0.003970644 | 66 |
| | ODA_P09769 Tyrosine-protein kinase Fgr (FGR) | 0.344327314 | 0.004640171 | 66 |
| | ODA_P09960 Leukotriene A-4 hydrolase (LTA4H) | 0.360776537 | 0.002921457 | 66 |
| | ODA_P40121 Macrophage-capping protein (CAPG) | 0.287757019 | 0.019135905 | 66 |
| | ODA_P42658 Dipeptidyl aminopeptidase-like protein 6 (DPP6) | -0.384114393 | 0.001452354 | 66 |
| | ODA_P49023 Paxillin (PXN) | 0.376933514 | 0.001810659 | 66 |
| | ODA_P53539 Protein fosB (FOSB) | 0.391838013 | 0.001139334 | 66 |
| | ODA_P55957 BH3-interacting domain death agonist (BID) | 0.260077236 | 0.034949166 | 66 |
| | ODA_P61244 Protein max (MAX) | 0.401273354 | 0.000840205 | 66 |
| | ODA_P80303 Nucleobindin-2 (NUCB2) | 0.318108757 | 0.009242701 | 66 |
| | ODA_P98073 Enteropeptidase (TMPRSS15) | -0.285672536 | 0.02006373 | 66 |
| | ODA_Q02246 Contactin-2 (CNTN2) | -0.369877883 | 0.002238005 | 66 |
| | ODA_Q02880 DNA topoisomerase 2-beta (TOP2B) | 0.420728525 | 0.000435664 | 66 |
| | ODA_Q07954 Prolow-density lipoprotein receptor-related protein 1 (LRP1) | 0.354597641 | 0.003485827 | 66 |
| | ODA_Q11201 CMP-N-acetylneuraminate-beta-galactosamide-alpha-2,3-sialyltransferase 1 (ST3GAL1) | 0.47379188 | 5.86375E-05 | 66 |

| | | | | |
|---|---|---|---|---|
| | ODA_Q12778 Forkhead box protein O1 (FOXO1) | 0.256946039 | 0.037282042 | 66 |
| | ODA_Q12913 Receptor-type tyrosine-protein phosphatase eta (PTPRJ) | 0.415635111 | 0.000519389 | 66 |
| | ODA_Q15165 Serum paraoxonase/arylesterase 2 (PON2) | 0.316063041 | 0.009729774 | 66 |
| | ODA_Q15797 Mothers against decapentaplegic homolog 1 (SMAD1) | 0.382653168 | 0.001519615 | 66 |
| | ODA_Q7L5Y9 Macrophage erythroblast attacher (MAEA) | 0.250057405 | 0.042872024 | 66 |
| | ODA_Q7LG56 Ribonucleoside-diphosphate reductase subunit M2 B (RRM2B) | 0.407452249 | 0.000684935 | 66 |
| | ODA_Q86SJ6 Desmoglein-4 (DSG4) | -0.377685002 | 0.001769763 | 66 |
| | ODA_Q86SR1 Polypeptide N-acetylgalactosaminyltransferase 10 (GALNT10) | 0.2474272 | 0.045181228 | 66 |
| | ODA_Q8N8S7 Protein enabled homolog (ENAH) | 0.314560067 | 0.010101713 | 66 |
| | ODA_Q8NDB2 B-cell scaffold protein with ankyrin repeats (BANK1) | 0.272435028 | 0.02689573 | 66 |
| | ODA_Q96RT1 Erbin (ERBIN) | 0.319528233 | 0.008917344 | 66 |
| | ODA_Q9GZY6 Linker for activation of T-cells family member 2 (LAT2) | 0.249013673 | 0.043776439 | 66 |
| | ODA_Q9NQ88 Fructose-2,6-bisphosphatase TIGAR (TIGAR) | 0.331009289 | 0.00663279 | 66 |
| | ODA_Q9NRA1 Platelet-derived growth factor C (PDGFC) | 0.253105104 | 0.040319002 | 66 |
| | ODA_Q9UKL0 REST corepressor 1 (RCOR1) | 0.346498278 | 0.0043713 | 66 |
| | ODA_Q9ULX7 Carbonic anhydrase 14 (CA14) | -0.313474585 | 0.010377965 | 66 |

| | | | | |
|---|---|---|---|---|
| | ODA_Q9UNK0 Syntaxin-8 (STX8) | 0.251894374 | 0.041317725 | 66 |
| | ODA_Q9Y478 5'-AMP-activated protein kinase subunit beta-1 (PRKAB1) | 0.249431166 | 0.043412803 | 66 |
| | ODA_Q9Y4K4 Mitogen-activated protein kinase kinase kinase kinase 5 (MAP4K5) | 0.296482622 | 0.015638894 | 66 |
| | ODA_Q9Y5A7 NEDD8 ultimate buster 1 (NUB1) | 0.310760881 | 0.011097482 | 66 |
| Positive mode plasma lipids | ChoE-18:1 | -0.337194337 | 0.044315496 | 36 |
| | ChoE-18:2 | -0.353410553 | 0.034481379 | 36 |
| | LPC-14:0 | -0.49034749 | 0.002397472 | 36 |
| | LPC-15:0 | -0.58018018 | 0.000208086 | 36 |
| | LPC-16:0 | -0.41956242 | 0.010856246 | 36 |
| | LPC-17:0 | -0.494465894 | 0.002173715 | 36 |
| | LPC-18:0 | -0.368339768 | 0.027076719 | 36 |
| | LPC-18:1 | -0.35006435 | 0.036349286 | 36 |
| | LPC-18:3 | -0.347490347 | 0.037841481 | 36 |
| | LPC-19:3 | -0.329472329 | 0.049734049 | 36 |
| | LPC-20:5 | -0.3747747 | 0.02431731 | 36 |

| | | | | |
|---|---|---|---|---|
| | | 75 | 1 | |
| | LPC-22:6 | -0.356756757 | 0.032692522 | 36 |
| | LPC-O-16:0 | -0.433719434 | 0.008225222 | 36 |
| | LPC-O-22:0 | -0.385070785 | 0.020388549 | 36 |
| | SM-17:0 | -0.337194337 | 0.044315496 | 36 |
| | SM-20:1 | -0.361389961 | 0.030341608 | 36 |
| | SM-26:5 | -0.333848134 | 0.046601989 | 36 |
| | SM-31:1;O2 17:1;O2/14:0 | -0.423423423 | 0.010076239 | 36 |
| | SM-32:0;O2 | -0.389189189 | 0.018972645 | 36 |
| | SM-33:2;O2 | -0.361647362 | 0.03021518 | 36 |
| | SM-35:1;O2 18:1;O2/17:0 | -0.337194337 | 0.044315496 | 36 |
| | SM-35:1;O2 21:1;O2/14:0 | -0.337194337 | 0.044315496 | 36 |
| | SM-38:2;O2 18:2;O2/20:0 | -0.361389961 | 0.030341608 | 36 |
| Urine eicosanoids | 11-dehydro TxB2 | 0.448739496 | 0.006854441 | 35 |
| | iPF2a-III | 0.335392535 | 0.045535178 | 36 |

Supplementary table 6
Positive mode targeted lipid list

| Index | exact.mass | ID | Duplicate p | Class |
|---|---|---|---|---|
| 1 | 468.3079 | LPC-14:0 | | LPC |
| 2 | 480.3087 | LPE-18:1 | | LPE |
| 3 | 480.3436 | LPC-15:1 | | LPC |
| 4 | 482.3245 | LPC-15:0 | | LPC |
| 5 | 482.3606 | LPC-O-16:0 | | LPC-O |
| 6 | 494.3243 | LPC-16:1 | | LPC |
| 7 | 496.3399 | LPC-16:0 | | LPC |
| 8 | 502.2926 | LPE-20:4 | | LPE |
| 9 | 508.3758 | LPC-17:1 | | LPC |
| 10 | 510.3558 | LPC-17:0 | | LPC |
| 11 | 518.3246 | LPC-18:3 | | LPC |
| 12 | 520.3394 | LPC-18:2 | 2 | LPC |
| 13 | 520.3397 | LPC-18:2 | 1 | LPC |
| 14 | 522.3538 | LPC-18:1 | | LPC |
| 15 | 524.3726 | LPC-18:0 | | LPC |
| 16 | 528.3084 | LPE-22:5 | | LPE |
| 17 | 532.3395 | LPC-19:3 | | LPC |
| 18 | 534.3914 | LPC-O-20:2 | | LPC-O |
| 19 | 536.3705 | LPC-19:1 | | LPC |
| 20 | 536.4075 | LPC-O-20:1 | | LPC-O |
| 21 | 538.3873 | LPC-19:0 | 2 | LPC |
| 22 | 538.3885 | LPC-19:0 | 1 | LPC |
| 23 | 542.3225 | LPC-20:5 | | LPC |
| 24 | 544.3421 | LPC-20:4 | | LPC |
| 25 | 546.3555 | LPC-20:3 | | LPC |
| 26 | 550.3869 | LPC-20:1 | | LPC |
| 27 | 552.4025 | LPC-20:0 | | LPC |
| 28 | 558.5094 | DG-30:0 | | DG |
| 29 | 566.4539 | LPC-O-22:0 | | LPC-O |
| 30 | 568.3428 | LPC-22:6 | | LPC |
| 31 | 578.418 | LPC-22:1 | | LPC |
| 32 | 580.4336 | LPC-22:0 | | LPC |
| 33 | 582.5093 | DG-32:2 | | DG |
| 34 | 584.5251 | DG-32:1 | | DG |
| 35 | 586.5408 | DG-32:0 | | DG |
| 36 | 594.5819 | Cer-38:1 | | Cer |
| 37 | 606.4501 | LPC-24:1 | | LPC |
| 38 | 606.5095 | DG-34:4 | | DG |
| 39 | 608.5248 | DG-34:3 | | DG |
| 40 | 610.5403 | DG-34:2 | | DG |
| 41 | 612.5562 | DG-34:1 | | DG |
| 42 | 614.5723 | DG-34:0 | | DG |
| 43 | 620.5978 | Cer-40:2 | | Cer |
| 44 | 622.6133 | Cer-40:1 | 3 | Cer |
| 45 | 632.5257 | DG-36:5 | | DG |
| 46 | 634.5405 | DG-36:4 | 2 | DG |

| | | | | |
|---|---|---|---|---|
| 47 | 634.5409 | DG-36:4 | 1 | DG |
| 48 | 634.6133 | Cer-41:2 | | Cer |
| 49 | 636.556 | DG-36:3 | | DG |
| 50 | 636.6295 | Cer-41:1 | | Cer |
| 51 | 638.5712 | DG-36:2 | | DG |
| 52 | 638.645 | Cer-41:0 | | Cer |
| 53 | 640.5872 | DG-36:1 | | DG |
| 54 | 642.6035 | DG-36:0 | | DG |
| 55 | 646.6137 | Cer-40:1 | 1 | Cer |
| 56 | 647.5124 | SM-30:1;O2 18:1;O2/1 | | SM |
| 57 | 648.6292 | Cer-40:1 | 2 | Cer |
| 58 | 648.6295 | Cer-42:2 | | Cer |
| 59 | 650.6445 | Cer-42:1 | | Cer |
| 60 | 656.5248 | DG-38:7 | | DG |
| 61 | 658.5405 | DG-38:6 | | DG |
| 62 | 660.5562 | DG-38:5 | | DG |
| 63 | 661.5275 | SM-31:1;O2 17:1;O2/1 | | SM |
| 64 | 663.4521 | SM-14:6 | | SM |
| 65 | 664.4935 | PC-28:7 | | PC |
| 66 | 664.6057 | ChoE-18:3 | | ChoE |
| 67 | 666.6193 | ChoE-18:2 | | ChoE |
| 68 | 668.6337 | ChoE-18:1 | | ChoE |
| 69 | 673.528 | SM-32:2;O2 18:2;O2/1 | | SM |
| 70 | 675.5436 | SM-32:1;O2 16:1;O2/1 | | SM |
| 71 | 675.544 | SM-14:0 | | SM |
| 72 | 677.5595 | SM-32:0;O2 | | SM |
| 73 | 687.5435 | SM-33:2;O2 | | SM |
| 74 | 688.6016 | ChoE-20:5 | | ChoE |
| 75 | 689.5591 | SM-33:1;O2 17:1;O2/1 | | SM |
| 76 | 689.5617 | SM-15:0 | | SM |
| 77 | 690.6189 | ChoE-20:4 | | ChoE |
| 78 | 692.559 | PC O-30:0 | | PC-O |
| 79 | 701.5564 | SM-16:1 | | SM |
| 80 | 701.5591 | SM-34:2;O2 18:2;O2/1 | | SM |
| 81 | 702.5056 | PC-30:2 | | PC |
| 82 | 703.5747 | SM-34:1;O2 | | SM |
| 83 | 703.5762 | SM-34:1;O2 18:1;O2/1 | | SM |
| 84 | 703.5767 | SM-16:0 | | SM |
| 85 | 704.5582 | PE O-34:1 | | PE-O |
| 86 | 705.5909 | SM-34:0;O2 18:0;O2/1 | | SM |
| 87 | 711.5465 | SM-17:3 | | SM |
| 88 | 714.5433 | PE O-35:3 | | PE-O |
| 89 | 714.6185 | ChoE-22:6 | | ChoE |
| 90 | 715.5753 | SM-35:2;O2 21:2;O2/1 | | SM |
| 91 | 716.5593 | PC O-32:2 | | PC-O |
| 92 | 717.5898 | SM-17:0 | | SM |
| 93 | 717.5908 | SM-35:1;O2 18:1;O2/1 | | SM |

| | | | | |
|---|---|---|---|---|
| 94 | 717.5911 | SM-35:1;O2 21:1;O2/1 | | SM |
| 95 | 718.5394 | PC-31:1 | | PC |
| 96 | 718.5748 | PC O-32:1 | | PC-O |
| 97 | 720.5544 | PC-31:0 | | PC |
| 98 | 720.5545 | PC-31:0 | | PC |
| 99 | 723.5447 | SM-18:4 | | SM |
| 100 | 725.56 | SM-18:3 | | SM |
| 101 | 727.5752 | SM-36:3;O2 18:1;O2/1 | | SM |
| 102 | 728.5228 | PC-32:3 | | PC |
| 103 | 729.5915 | SM-36:2;O2 18:2;O2/1 | | SM |
| 104 | 729.5917 | SM-18:1 | | SM |
| 105 | 730.5387 | PC-32:2 | | PC |
| 106 | 731.6069 | SM-36:1;O2 16:1;O2/2 | | SM |
| 107 | 731.6089 | SM-18:0 | | SM |
| 108 | 732.5576 | PC-32:1 | | PC |
| 109 | 733.6226 | SM-36:0;O2 | | SM |
| 110 | 734.5718 | PC-32:0 | | PC |
| 111 | 734.6068 | PC O-33:0 | | PC-O |
| 112 | 739.5718 | SM-37:4;O2 | | SM |
| 113 | 740.6752 | TG-42:0 | | TG |
| 114 | 742.5386 | PC-33:3 | | PC |
| 115 | 742.5741 | PC O-34:3 | 1 | PC-O |
| 116 | 742.575 | PC O-34:3 | 2 | PC-O |
| 117 | 745.6218 | SM-37:1;O2 18:1;O2/1 | | SM |
| 118 | 746.605 | PC O-34:1 | | PC-O |
| 119 | 748.5283 | PE O-38:7 | | PE-O |
| 120 | 748.5856 | PC-33:0 | | PC |
| 121 | 748.6213 | PC O-34:0 | | PC-O |
| 122 | 751.5771 | SM-20:4 | | SM |
| 123 | 752.522 | PC-34:5 | | PC |
| 124 | 753.5944 | SM-20:3 | | SM |
| 125 | 754.5378 | PC-34:4 | | PC |
| 126 | 754.5747 | PE O-38:4 | | PE-O |
| 127 | 754.5748 | PC O-35:4 | 1 | PC-O |
| 128 | 754.5754 | PC O-35:4 | 2 | PC-O |
| 129 | 754.6968 | TG-43:0 | | TG |
| 130 | 755.6072 | SM-38:3;O2 | | SM |
| 131 | 756.5535 | PC-34:3 | | PC |
| 132 | 756.5908 | PC O-35:3 | 1 | PC-O |
| 133 | 756.5909 | PC O-35:3 | 2 | PC-O |
| 134 | 757.6225 | SM-20:1 | | SM |
| 135 | 757.6226 | SM-38:2;O2 18:2;O2/2 | | SM |
| 136 | 758.5718 | PC-34:2 | | PC |
| 137 | 758.6063 | PC O-35:2 | | PC-O |
| 138 | 759.6376 | SM-38:1;O2 18:1;O2/2 | | SM |
| 139 | 759.6407 | SM-20:0 | | SM |
| 140 | 760.5883 | PC-34:1 | | PC |

| | | | | |
|---|---|---|---|---|
| 141 | 761.652 | SM-38:0;O2 | | SM |
| 142 | 762.6032 | PC-34:0 | | PC |
| 143 | 764.5231 | PC-35:6 | | PC |
| 144 | 764.6807 | TG-44:2 | | TG |
| 145 | 766.538 | PC-35:5 | | PC |
| 146 | 766.5385 | PC-35:5 | | PC |
| 147 | 766.575 | PC O-36:5 | 2 | PC-O |
| 148 | 766.5752 | PC O-36:5 | 1 | PC-O |
| 149 | 766.6937 | TG-44:1 | | TG |
| 150 | 768.5568 | PC-35:4 | | PC |
| 151 | 768.5771 | PC O-36:4 | | PC-O |
| 152 | 770.5698 | PC-35:3 | | PC |
| 153 | 772.5274 | PE O-40:9 | | PE-O |
| 154 | 772.5858 | PC-35:2 | 2 | PC |
| 155 | 772.5861 | PC-35:2 | 1 | PC |
| 156 | 772.6209 | PC O-36:2 | 1 | PC-O |
| 157 | 772.6215 | PC O-36:2 | 2 | PC-O |
| 158 | 773.6527 | SM-39:1;O2 16:1;O2/2 | | SM |
| 159 | 773.6563 | SM-21:0 | | SM |
| 160 | 774.6008 | PC-35:1 | | PC |
| 161 | 778.5362 | PC-36:6 | 1 | PC |
| 162 | 778.5386 | PC-36:6 | 2 | PC |
| 163 | 780.5514 | PC-36:5 | 1 | PC |
| 164 | 780.5523 | PC-36:5 | 2 | PC |
| 165 | 780.5905 | PC O-37:5 | | PC-O |
| 166 | 782.5695 | PC-36:4 | 4 | PC |
| 167 | 782.5715 | PC-36:4 | 1 | PC |
| 168 | 782.5718 | PC-36:4 | 2 | PC |
| 169 | 782.5728 | PC-36:4 | 3 | PC |
| 170 | 782.593 | PC O-37:4 | | PC-O |
| 171 | 782.7242 | TG-45:0 | | TG |
| 172 | 783.6379 | SM-40:3;O2 | | SM |
| 173 | 784.5768 | PC-36:3 | 2 | PC |
| 174 | 784.5831 | PC-36:3 | 1 | PC |
| 175 | 785.653 | SM-40:2;O2 18:2;O2/2 | | SM |
| 176 | 785.6532 | SM-40:2;O2 16:1;O2/2 | | SM |
| 177 | 785.6536 | SM-22:1 | 1 | SM |
| 178 | 785.657 | SM-22:1 | 2 | SM |
| 179 | 786.6012 | PC-36:2 | | PC |
| 180 | 787.6685 | SM-40:1;O2 18:1;O2/2 | | SM |
| 181 | 787.6719 | SM-22:0 | | SM |
| 182 | 788.6165 | PC-36:1 | | PC |
| 183 | 789.684 | SM-40:0;O2 | | SM |
| 184 | 790.5381 | PC-37:7 | | PC |
| 185 | 790.5723 | PC O-38:7 | 1 | PC-O |
| 186 | 790.5743 | PC O-38:7 | 3 | PC-O |
| 187 | 790.5756 | PC O-38:7 | 2 | PC-O |

| | | | | |
|---|---|---|---|---|
| 188 | 790.6325 | PC-36:0 | | PC |
| 189 | 790.6967 | TG-46:3 | | TG |
| 190 | 792.5908 | PC O-38:6 | | PC-O |
| 191 | 792.7114 | TG-46:2 | | TG |
| 192 | 794.569 | PC-37:5 | | PC |
| 193 | 794.6059 | PC O-38:5 | 1 | PC-O |
| 194 | 794.6062 | PC O-38:5 | 2 | PC-O |
| 195 | 794.722 | TG-46:1 | | TG |
| 196 | 796.5855 | PC-37:4 | | PC |
| 197 | 796.7368 | TG-46:0 | | TG |
| 198 | 798.6011 | PC-37:3 | | PC |
| 199 | 798.6368 | PC O-38:3 | | PC-O |
| 200 | 799.6657 | SM-23:1 | 1 | SM |
| 201 | 799.6688 | SM-41:2;O2 17:1;O2/2 | | SM |
| 202 | 799.6701 | SM-23:1 | 2 | SM |
| 203 | 801.6845 | SM-41:1;O2 18:1;O2/2 | | SM |
| 204 | 801.6848 | SM-41:1;O2 | | SM |
| 205 | 801.6851 | SM-23:0 | 2 | SM |
| 206 | 801.6882 | SM-23:0 | 1 | SM |
| 207 | 802.5403 | PC-38:8 | | PC |
| 208 | 802.6321 | PC-37:1 | | PC |
| 209 | 804.5527 | PC-37:0 | | PC |
| 210 | 804.5535 | PC-38:7 | | PC |
| 211 | 804.5905 | PE O-42:7 | | PE-O |
| 212 | 804.7063 | TG-47:3 | | TG |
| 213 | 806.5669 | PC-38:6 | 1 | PC |
| 214 | 806.5672 | PC-38:6 | 2 | PC |
| 215 | 806.7217 | TG-47:2 | | TG |
| 216 | 807.6346 | SM-24:4 | 1 | SM |
| 217 | 807.6374 | SM-42:5;O2 | | SM |
| 218 | 807.6377 | SM-24:4 | 2 | SM |
| 219 | 808.5831 | PC-38:5 | | PC |
| 220 | 808.739 | TG-47:1 | | TG |
| 221 | 809.6502 | SM-42:4;O2 18:1;O2/2 | | SM |
| 222 | 809.6506 | SM-24:3 | | SM |
| 223 | 809.6533 | SM-42:4;O2 18:2;O2/2 | | SM |
| 224 | 810.5992 | PC-38:4 | 4 | PC |
| 225 | 810.6016 | PC-38:4 | 2 | PC |
| 226 | 810.6037 | PC-38:4 | 3 | PC |
| 227 | 810.6044 | PC-38:4 | 1 | PC |
| 228 | 811.6653 | SM-24:2 | | SM |
| 229 | 811.6691 | SM-42:3;O2 18:2;O2/2 | | SM |
| 230 | 812.6152 | PC-38:3 | | PC |
| 231 | 813.6838 | SM-24:1 | 1 | SM |
| 232 | 813.685 | SM-42:2;O2 18:1;O2/2 | | SM |
| 233 | 813.6871 | SM-24:1 | 2 | SM |
| 234 | 814.5734 | PC O-40:9 | | PC-O |

| | | | | |
|---|---|---|---|---|
| 235 | 814.6338 | PC-38:2 | | PC |
| 236 | 815.6998 | SM-42:1;O2 | | SM |
| 237 | 815.7006 | SM-42:1;O2 18:1;O2/2 | | SM |
| 238 | 815.7028 | SM-24:0 | | SM |
| 239 | 816.5884 | PC O-40:8 | 1 | PC-O |
| 240 | 816.5898 | PC O-40:8 | 2 | PC-O |
| 241 | 816.5906 | PC O-40:8 | 3 | PC-O |
| 242 | 816.6472 | PC-38:1 | | PC |
| 243 | 818.5699 | PC-39:7 | | PC |
| 244 | 818.6038 | PC O-40:7 | 3 | PC-O |
| 245 | 818.606 | PC O-40:7 | 2 | PC-O |
| 246 | 818.6063 | PC O-40:7 | 4 | PC-O |
| 247 | 818.6064 | PC O-40:7 | 1 | PC-O |
| 248 | 818.6649 | PC-38:0 | | PC |
| 249 | 818.7239 | TG-48:3 | 2 | TG |
| 250 | 818.7251 | TG-48:3 | 1 | TG |
| 251 | 820.5859 | PC-39:6 | | PC |
| 252 | 820.6201 | PC O-40:6 | 2 | PC-O |
| 253 | 820.6208 | PC O-40:6 | 1 | PC-O |
| 254 | 820.6224 | PC O-40:6 | | PC-O |
| 255 | 820.7375 | TG-48:2 | | TG |
| 256 | 822.6011 | PC-39:5 | | PC |
| 257 | 822.6373 | PC O-40:5 | 2 | PC-O |
| 258 | 822.638 | PC O-40:5 | 1 | PC-O |
| 259 | 822.7565 | TG-48:1 | | TG |
| 260 | 823.6712 | SM-25:3 | | SM |
| 261 | 824.6165 | PC-39:4 | | PC |
| 262 | 824.6531 | PC O-40:4 | 2 | PC-O |
| 263 | 824.6534 | PC O-40:4 | 1 | PC-O |
| 264 | 824.77 | TG-48:0 | 2 | TG |
| 265 | 824.7726 | TG-48:0 | 1 | TG |
| 266 | 826.5356 | PC-40:10 | | PC |
| 267 | 826.6689 | PC O-40:3 | 1 | PC-O |
| 268 | 826.6738 | PC O-40:3 | 2 | PC-O |
| 269 | 827.7 | SM-43:2;O2 18:2;O2/2 | | SM |
| 270 | 827.7007 | SM-43:2;O | 1 | SM |
| 271 | 827.7011 | SM-43:2;O | 2 | SM |
| 272 | 827.7049 | SM-25:1 | | SM |
| 273 | 828.5542 | PC-40:9 | | PC |
| 274 | 829.7124 | SM-25:0 | | SM |
| 275 | 829.7154 | SM-43:1;O2 | | SM |
| 276 | 829.7158 | SM-43:1;O2 18:1;O2/2 | | SM |
| 277 | 832.5808 | PC-39:0 | | PC |
| 278 | 832.583 | PC-40:7 | | PC |
| 279 | 832.5853 | PC-40:7 | | PC |
| 280 | 832.744 | TG-49:3 | | TG |
| 281 | 833.6518 | SM-26:5 | | SM |

| | | | | |
|---|---|---|---|---|
| 282 | 834.5983 | PC-40:6 | | PC |
| 283 | 834.6 | PC-40:6 | | PC |
| 284 | 834.7592 | TG-49:2 | | TG |
| 285 | 835.6672 | SM-26:4 | 2 | SM |
| 286 | 835.6685 | SM-26:4 | 1 | SM |
| 287 | 836.6119 | PC-40:5 | | PC |
| 288 | 837.6869 | SM-26:3 | | SM |
| 289 | 839.7038 | SM-26:2 | | SM |
| 290 | 840.6478 | PC-40:3 | 2 | PC |
| 291 | 840.6486 | PC-40:3 | 1 | PC |
| 292 | 841.7189 | SM-26:1 | | SM |
| 293 | 842.6638 | PC-40:2 | | PC |
| 294 | 842.7266 | TG-50:5 | | TG |
| 295 | 844.6195 | PC O-42:8 | 1 | PC-O |
| 296 | 844.6223 | PC O-42:8 | 2 | PC-O |
| 297 | 844.7397 | TG-50:4 | | TG |
| 298 | 846.6364 | PC O-42:7 | 1 | PC-O |
| 299 | 846.6367 | PC O-42:7 | 3 | PC-O |
| 300 | 846.6382 | PC O-42:7 | 2 | PC-O |
| 301 | 846.7554 | TG-50:3 | 2 | TG |
| 302 | 846.7567 | TG-50:3 | 1 | TG |
| 303 | 848.6528 | PC O-42:6 | 1 | PC-O |
| 304 | 848.653 | PC O-42:6 | 2 | PC-O |
| 305 | 848.7715 | TG-50:2 | | TG |
| 306 | 850.6685 | PC O-42:5 | 1 | PC-O |
| 307 | 850.6687 | PC O-42:5 | 2 | PC-O |
| 308 | 850.7871 | TG-50:1 | | TG |
| 309 | 852.6838 | PC O-42:4 | 1 | PC-O |
| 310 | 852.6847 | PC O-42:4 | 2 | PC-O |
| 311 | 854.7004 | PC O-42:3 | | PC-O |
| 312 | 855.7324 | SM-27:1 | | SM |
| 313 | 856.585 | PC-42:9 | | PC |
| 314 | 856.5852 | PC-42:9 | | PC |
| 315 | 856.7147 | PC O-42:2 | | PC-O |
| 316 | 856.7151 | PC O-42:2 | | PC-O |
| 317 | 858.6001 | PC-42:8 | | PC |
| 318 | 858.7546 | TG-51:4 | | TG |
| 319 | 860.6141 | PC-42:7 | | PC |
| 320 | 860.6174 | PC-42:7 | | PC |
| 321 | 860.7722 | TG-51:3 | | TG |
| 322 | 862.6323 | PC-42:6 | 1 | PC |
| 323 | 862.6334 | PC-42:6 | 2 | PC |
| 324 | 862.7913 | TG-51:2 | | TG |
| 325 | 864.6472 | PC-42:5 | | PC |
| 326 | 864.8056 | TG-51:1 | 1 | TG |
| 327 | 864.8061 | TG-51:1 | 2 | TG |
| 328 | 866.6629 | PC-42:4 | | PC |

| | | | | |
|---|---|---|---|---|
| 329 | 866.6647 | PC-42:4 | | PC |
| 330 | 866.8201 | TG-51:0 | | TG |
| 331 | 868.74 | TG-52:6 | | TG |
| 332 | 870.6345 | PC O-44:9 | | PC-O |
| 333 | 870.6355 | PC O-44:9 | | PC-O |
| 334 | 870.6952 | PC-42:2 | | PC |
| 335 | 870.7558 | TG-52:5 | | TG |
| 336 | 872.6536 | PC O-44:8 | | PC-O |
| 337 | 872.772 | TG-52:4 | | TG |
| 338 | 874.6682 | PC O-44:7 | | PC-O |
| 339 | 874.6685 | PC O-44:7 | | PC-O |
| 340 | 874.7885 | TG-52:3 | | TG |
| 341 | 876.8063 | TG-52:2 | | TG |
| 342 | 878.6985 | PC O-44:5 | | PC-O |
| 343 | 878.8224 | TG-52:1 | | TG |
| 344 | 880.715 | PC O-44:4 | 1 | PC-O |
| 345 | 880.7159 | PC O-44:4 | 2 | PC-O |
| 346 | 880.8375 | TG-52:0 | | TG |
| 347 | 885.7817 | SM-29:0 | | SM |
| 348 | 890.8201 | TG-53:2 | | TG |
| 349 | 892.8337 | TG-53:1 | | TG |
| 350 | 893.751 | SM-30:3 | | SM |
| 351 | 894.6971 | PC-44:4 | | PC |
| 352 | 894.7515 | TG-54:7 | | TG |
| 353 | 896.6501 | PC O-46:10 | | PC-O |
| 354 | 896.7118 | PC-44:3 | | PC |
| 355 | 896.772 | TG-54:6 | 2 | TG |
| 356 | 896.774 | TG-54:6 | 1 | TG |
| 357 | 898.7225 | PC-44:2 | | PC |
| 358 | 898.7861 | TG-54:5 | | TG |
| 359 | 900.6853 | PC O-46:8 | | PC-O |
| 360 | 900.7373 | PC-44:1 | | PC |
| 361 | 900.8053 | TG-54:4 | | TG |
| 362 | 902.8207 | TG-54:3 | | TG |
| 363 | 904.7165 | PC O-46:6 | | PC-O |
| 364 | 904.8358 | TG-54:2 | | TG |
| 365 | 906.8511 | TG-54:1 | | TG |
| 366 | 908.8682 | TG-54:0 | | TG |
| 367 | 916.8366 | TG-55:3 | | TG |
| 368 | 918.8494 | TG-55:2 | | TG |
| 369 | 920.7708 | TG-56:8 | | TG |
| 370 | 920.869 | TG-55:1 | | TG |
| 371 | 922.7908 | TG-56:7 | | TG |
| 372 | 924.7462 | PC-46:3 | | PC |
| 373 | 924.8026 | TG-56:6 | | TG |
| 374 | 926.8182 | TG-56:5 | | TG |
| 375 | 928.769 | PC-46:1 | | PC |

| | | | |
|---|---|---|---|
| 376 | 928.8317 | TG-56:4 | TG |
| 377 | 930.8494 | TG-56:3 | TG |
| 378 | 932.86 | TG-56:2 | TG |
| 379 | 934.8801 | TG-56:1 | TG |
| 380 | 944.7735 | TG-58:10 | TG |
| 381 | 946.7879 | TG-58:9 | TG |
| 382 | 946.8836 | TG-57:2 | TG |
| 383 | 948.9012 | TG-57:1 | TG |
| 384 | 952.8344 | TG-58:6 | TG |
| 385 | 956.8646 | TG-58:4 | TG |
| 386 | 958.8865 | TG-58:3 | TG |
| 387 | 960.8916 | TG-58:2 | TG |
| 388 | 976.9298 | TG-59:1 | TG |
| 389 | 978.9465 | TG-59:0 | TG |
| 390 | 988.9258 | TG-60:2 | TG |
| 391 | 990.9436 | TG-60:1 | TG |
| 392 | 1014.942 | TG-62:3 | TG |
| 393 | 1016.962 | TG-62:2 | TG |

Negative mode targeted lipid list

| Index | exact.mass | ID | peak | Class |
|---|---|---|---|---|
| 1 | 526.3521 | LPC O-16:1 | | LPC-O |
| 2 | 554.3834 | LPC O-18:0 | | LPC-O |
| 3 | 552.3669 | LPC O-18:1 | 1 | LPC-O |
| 4 | 552.3671 | LPC O-18:1 | 2 | LPC-O |
| 5 | 582.4139 | LPC O-20:0 | | LPC-O |
| 6 | 636.4618 | LPC O-24:1 | | LPC-O |
| 7 | 634.4457 | LPC O-24:2 | | LPC-O |
| 8 | 436.2837 | LPE O-16:1 | | LPE-O |
| 9 | 464.3146 | LPE O-18:1 | | LPE-O |
| 10 | 462.2985 | LPE O-18:2 | | LPE-O |
| 11 | 492.3449 | LPE O-20:1 | | LPE-O |
| 12 | 764.5797 | PC O-32:0 | | PC-O |
| 13 | 762.5646 | PC O-32:1 | | PC-O |
| 14 | 790.5949 | PC O-34:1 | | PC-O |
| 15 | 788.5808 | PC O-34:2 | 1 | PC-O |
| 16 | 788.5813 | PC O-34:2 | 2 | PC-O |
| 17 | 786.5646 | PC O-34:3 | | PC-O |
| 18 | 816.6114 | PC O-36:2 | | PC-O |
| 19 | 814.5961 | PC O-36:3 | 1 | PC-O |
| 20 | 814.5947 | PC O-36:3 | 2 | PC-O |
| 21 | 814.5949 | PC O-36:3 | 3 | PC-O |
| 22 | 812.5806 | PC O-36:4 | | PC-O |
| 23 | 810.5651 | PC O-36:5 | | PC-O |
| 24 | 840.611 | PC O-38:4 | 1 | PC-O |
| 25 | 840.6113 | PC O-38:4 | 2 | PC-O |
| 26 | 838.5955 | PC O-38:5 | 1 | PC-O |
| 27 | 838.5972 | PC O-38:5 | 2 | PC-O |
| 28 | 836.5804 | PC O-38:6 | | PC-O |
| 29 | 834.5637 | PC O-38:7 | | PC-O |
| 30 | 868.6436 | PC O-40:4 | | PC-O |
| 31 | 866.6277 | PC O-40:5 | | PC-O |
| 32 | 864.6101 | PC O-40:6 | | PC-O |
| 33 | 894.6594 | PC O-42:5 | | PC-O |
| 34 | 922.6912 | PC O-44:5 | | PC-O |
| 35 | 920.6752 | PC O-44:6 | | PC-O |
| 36 | 700.5289 | PE O-34:2 | | PE-O |
| 37 | 698.5128 | PE O-34:3 | | PE-O |
| 38 | 728.5596 | PE O-36:2 | | PE-O |
| 39 | 726.5438 | PE O-36:3 | | PE-O |
| 40 | 724.5276 | PE O-36:4 | | PE-O |
| 41 | 722.5123 | PE O-36:5 | | PE-O |
| 42 | 720.4961 | PE O-36:6 | | PE-O |
| 43 | 736.527 | PE O-37:5 | | PE-O |
| 44 | 754.5761 | PE O-38:3 | | PE-O |
| 45 | 752.559 | PE O-38:4 | | PE-O |
| 46 | 750.5428 | PE O-38:5 | 1 | PE-O |

| | | | | |
|---|---|---|---|---|
| 47 | 750.5429 | PE O-38:5 | 3 | PE-O |
| 48 | 750.5438 | PE O-38:5 | 2 | PE-O |
| 49 | 748.5265 | PE O-38:6 | 1 | PE-O |
| 50 | 748.527 | PE O-38:6 | 2 | PE-O |
| 51 | 746.5131 | PE O-38:7 | | PE-O |
| 52 | 778.5747 | PE O-40:5 | | PE-O |
| 53 | 776.5593 | PE O-40:6 | | PE-O |
| 54 | 774.5435 | PE O-40:7 | | PE-O |
| 55 | 772.5282 | PE O-40:8 | | PE-O |
| 56 | 227.2018 | FA 14:0 | | FA |
| 57 | 255.2332 | FA 16:0 | | FA |
| 58 | 253.2174 | FA 16:1 | | FA |
| 59 | 269.2488 | FA 17:0 | | FA |
| 60 | 283.2643 | FA 18:0 | | FA |
| 61 | 281.2494 | FA 18:1 | | FA |
| 62 | 279.2334 | FA 18:2 | | FA |
| 63 | 277.2172 | FA 18:3 | 2 | FA |
| 64 | 277.2174 | FA 18:3 | 1 | FA |
| 65 | 311.2958 | FA 20:0 | | FA |
| 66 | 309.2804 | FA 20:1 | | FA |
| 67 | 303.2331 | FA 20:4 | | FA |
| 68 | 301.2177 | FA 20:5 | | FA |
| 69 | 339.3271 | FA 22:0 | | FA |
| 70 | 331.2643 | FA 22:4 | | FA |
| 71 | 329.2486 | FA 22:5 | 1 | FA |
| 72 | 329.2485 | FA 22:5 | 2 | FA |
| 73 | 327.233 | FA 22:6 | | FA |
| 74 | 365.3425 | FA 24:1 | | FA |
| 75 | 744.5613 | HexCer 34:1-O2 | | HexCer |
| 76 | 800.6254 | HexCer 38:1-O2 | | HexCer |
| 77 | 828.6566 | HexCer 40:1-O2 | | HexCer |
| 78 | 409.2351 | LPA 16:0 | | LPA |
| 79 | 437.2681 | LPA 18:0 | | LPA |
| 80 | 435.252 | LPA 18:1 | | LPA |
| 81 | 433.2363 | LPA 18:2 | | LPA |
| 82 | 459.2506 | LPA 20:3 | | LPA |
| 83 | 457.2363 | LPA 20:4 | | LPA |
| 84 | 481.2359 | LPA 22:6 | | LPA |
| 85 | 512.2994 | LPC 14:0 | | LPC |
| 86 | 526.3149 | LPC 15:0 | | LPC |
| 87 | 540.3308 | LPC 16:0 | 2 | LPC |
| 88 | 540.3309 | LPC 16:0 | 1 | LPC |
| 89 | 538.3154 | LPC 16:1 | | LPC |
| 90 | 554.3463 | LPC 17:0 | | LPC |
| 91 | 552.3314 | LPC 17:1 | | LPC |
| 92 | 568.3622 | LPC 18:0 | | LPC |
| 93 | 568.3621 | LPC 18:0 | | LPC |

| | | | | |
|---|---|---|---|---|
| 94 | 566.3463 | LPC 18:1 | | LPC |
| 95 | 566.3466 | LPC 18:1 | | LPC |
| 96 | 564.3309 | LPC 18:2 | | LPC |
| 97 | 564.3304 | LPC 18:2 | | LPC |
| 98 | 582.376 | LPC 19:0 | | LPC |
| 99 | 582.3782 | LPC 19:0 | | LPC |
| 100 | 596.3932 | LPC 20:0 | | LPC |
| 101 | 594.3771 | LPC 20:1 | | LPC |
| 102 | 592.3622 | LPC 20:2 | | LPC |
| 103 | 590.3467 | LPC 20:3 | | LPC |
| 104 | 588.331 | LPC 20:4 | 2 | LPC |
| 105 | 588.33 | LPC 20:4 | 1 | LPC |
| 106 | 616.3627 | LPC 22:4 | | LPC |
| 107 | 614.346 | LPC 22:5 | 1 | LPC |
| 108 | 614.3459 | LPC 22:5 | 2 | LPC |
| 109 | 612.331 | LPC 22:6 | | LPC |
| 110 | 652.4555 | LPC 24:0 | | LPC |
| 111 | 452.2782 | LPE 16:0 | 2 | LPE |
| 112 | 452.2779 | LPE 16:0 | 1 | LPE |
| 113 | 480.3091 | LPE 18:0 | 2 | LPE |
| 114 | 480.3095 | LPE 18:0 | 1 | LPE |
| 115 | 478.2938 | LPE 18:1 | | LPE |
| 116 | 476.2788 | LPE 18:2 | 2 | LPE |
| 117 | 476.2785 | LPE 18:2 | 1 | LPE |
| 118 | 502.2933 | LPE 20:3 | | LPE |
| 119 | 500.2786 | LPE 20:4 | | LPE |
| 120 | 528.3096 | LPE 22:4 | | LPE |
| 121 | 526.2947 | LPE 22:5 | | LPE |
| 122 | 509.2889 | LPG 18:1 | | LPG |
| 123 | 507.272 | LPG 18:2 | | LPG |
| 124 | 571.2899 | LPI 16:0 | | LPI |
| 125 | 599.3203 | LPI 18:0 | | LPI |
| 126 | 597.304 | LPI 18:1 | | LPI |
| 127 | 524.2999 | LPS 18:0 | | LPS |
| 128 | 522.2827 | LPS 18:1 | | LPS |
| 129 | 750.5283 | PC 30:0 | | PC |
| 130 | 778.5603 | PC 32:0 | | PC |
| 131 | 776.5436 | PC 32:1 | | PC |
| 132 | 774.5281 | PC 32:2 | | PC |
| 133 | 788.5441 | PC 33:2 | | PC |
| 134 | 806.5911 | PC 34:0 | | PC |
| 135 | 804.5755 | PC 34:1 | | PC |
| 136 | 802.5609 | PC 34:2 | | PC |
| 137 | 800.5433 | PC 34:3 | | PC |
| 138 | 800.5443 | PC 34:3 | | PC |
| 139 | 798.5273 | PC 34:4 | | PC |
| 140 | 816.5751 | PC 35:2 | 2 | PC |

| | | | | |
|---|---|---|---|---|
| 141 | 816.5756 | PC 35:2 | 1 | PC |
| 142 | 812.5436 | PC 35:4 | | PC |
| 143 | 832.6069 | PC 36:1 | | PC |
| 144 | 830.5914 | PC 36:2 | | PC |
| 145 | 828.5754 | PC 36:3 | 1 | PC |
| 146 | 828.5762 | PC 36:3 | 2 | PC |
| 147 | 826.5604 | PC 36:4 | 1 | PC |
| 148 | 826.5587 | PC 36:4 | 2 | PC |
| 149 | 824.5441 | PC 36:5 | 1 | PC |
| 150 | 824.5447 | PC 36:5 | 2 | PC |
| 151 | 824.5436 | PC 36:5 | 3 | PC |
| 152 | 822.5289 | PC 36:6 | | PC |
| 153 | 844.6072 | PC 37:2 | | PC |
| 154 | 842.5912 | PC 37:3 | | PC |
| 155 | 836.5436 | PC 37:6 | | PC |
| 156 | 858.622 | PC 38:2 | | PC |
| 157 | 856.6069 | PC 38:3 | | PC |
| 158 | 854.5911 | PC 38:4 | 1 | PC |
| 159 | 854.5914 | PC 38:4 | 2 | PC |
| 160 | 854.5904 | PC 38:4 | 3 | PC |
| 161 | 852.5763 | PC 38:5 | 1 | PC |
| 162 | 852.5759 | PC 38:5 | 2 | PC |
| 163 | 850.5598 | PC 38:6 | 1 | PC |
| 164 | 850.5593 | PC 38:6 | 2 | PC |
| 165 | 848.5442 | PC 38:7 | 1 | PC |
| 166 | 848.5437 | PC 38:7 | 2 | PC |
| 167 | 864.575 | PC 39:6 | | PC |
| 168 | 882.6227 | PC 40:4 | 1 | PC |
| 169 | 882.623 | PC 40:4 | 2 | PC |
| 170 | 880.6063 | PC 40:5 | 1 | PC |
| 171 | 880.607 | PC 40:5 | 2 | PC |
| 172 | 878.5912 | PC 40:6 | 1 | PC |
| 173 | 878.5897 | PC 40:6 | 2 | PC |
| 174 | 876.5743 | PC 40:7 | | PC |
| 175 | 874.5613 | PC 40:8 | | PC |
| 176 | 898.5605 | PC 42:10 | | PC |
| 177 | 716.5233 | PE 34:1 | | PE |
| 178 | 714.5075 | PE 34:2 | | PE |
| 179 | 744.5549 | PE 36:1 | | PE |
| 180 | 742.5383 | PE 36:2 | | PE |
| 181 | 740.5228 | PE 36:3 | | PE |
| 182 | 738.5077 | PE 36:4 | 1 | PE |
| 183 | 738.5073 | PE 36:4 | 2 | PE |
| 184 | 768.5548 | PE 38:3 | | PE |
| 185 | 766.5382 | PE 38:4 | | PE |
| 186 | 764.5226 | PE 38:5 | 1 | PE |
| 187 | 764.5237 | PE 38:5 | 2 | PE |

| | | | |
|---|---|---|---|
| 188 | 762.507 | PE 38:6 | PE |
| 189 | 794.5702 | PE 40:4 | PE |
| 190 | 792.5533 | PE 40:5 | PE |
| 191 | 790.5383 | PE 40:6 | PE |
| 192 | 807.5031 | PI 32:1 | PI |
| 193 | 835.5353 | PI 34:1 | PI |
| 194 | 833.5187 | PI 34:2 | PI |
| 195 | 863.5644 | PI 36:1 | PI |
| 196 | 861.5504 | PI 36:2 | PI |
| 197 | 859.5335 | PI 36:3 | PI |
| 198 | 857.5181 | PI 36:4 | PI |
| 199 | 887.5651 | PI 38:3 | PI |
| 200 | 885.5505 | PI 38:4 | PI |
| 201 | 883.5335 | PI 38:5 | PI |
| 202 | 881.5185 | PI 38:6 | PI |
| 203 | 909.5477 | PI 40:6 | PI |
| 204 | 788.5445 | PS 36:1 | PS |
| 205 | 810.5275 | PS 38:4 | PS |

**Supplementary Table 7:** Features (Clinical and demograhic, immunotypes and proteins) assoiated with LPC-O-16:0, PC-O-30:0 and ChoE-18:3 (Spearman's rho < -0.4 or > 0.4, p < 0.05)

| Dataset | Correlated feature | Spearman's rho | p | n | Moderate vs severe covid q value |
|---|---|---|---|---|---|
| LPC O -16:0 | | | | | |
| Clinical and demographic | ards_ever | -0.4 | 0.00087 | 67 | 7.56E-24 |
| | daysaliveventfree_in28 | 0.51 | 7.77E-05 | 55 | 2.56E-18 |
| | hsCRP | -0.58 | 0.011084 | 18 | 1.15E-05 |
| | intubated_yn | -0.43 | 0.000468 | 63 | 4.12E-22 |
| | ordinal_d7 | -0.4 | 0.000887 | 67 | 4.82E-26 |
| | ordinal_enrollment | -0.4 | 0.000803 | 67 | 2.58E-46 |
| | pct | -0.44 | 0.004874 | 40 | 1.10E-05 |
| Immunotype | percOf_CD4_CD4EMRA | -0.47 | 0.002467 | 40 | 0.751329993 |
| | percOf_CD4_CD4NNKI67pos | -0.46 | 0.002831 | 40 | 0.046842496 |
| | percOf_CD4_KI67pos | -0.47 | 0.002373 | 40 | 0.08582355 |
| | percOf_CD4_TBETpos | -0.45 | 0.00401 | 40 | 0.548651911 |
| | percOf_CD4CM_KI67pos | -0.43 | 0.005638 | 40 | 0.012293841 |
| | percOf_CD4cTfh_CD4acTfh | -0.44 | 0.004168 | 40 | 0.025195859 |
| | percOf_CD4cTfh_HLADRposCD38pos | -0.52 | 0.00 | 40 | 0.012241891 |

| | | | | | |
|---|---|---|---|---|---|
| | | | 0621 | | |
| | percOf_CD4cTfh_KI67pos | -0.53 | 0.000379 | 40 | 0.301400751 |
| | percOf_CD4EM1_HLADRposCD38pos | -0.43 | 0.005523 | 40 | 0.000783423 |
| | percOf_CD4EM1_KI67pos | -0.58 | 7.40E-05 | 40 | 0.026755803 |
| | percOf_CD4nonNaive_KI67pos | -0.42 | 0.006672 | 40 | 0.031992847 |
| | percOf_CD4nonNaive_TBETpos | -0.47 | 0.002388 | 40 | 0.544146679 |
| | percOf_CD8_CD8EM1 | 0.4 | 0.011225 | 40 | 0.087303203 |
| | percOf_CD8_CD8EMRA | -0.48 | 0.001653 | 40 | 0.384334434 |
| | percOf_CD8_TBETpos | -0.48 | 0.001862 | 40 | 0.621801498 |
| | percOf_CD8nonNaive_TBETpos | -0.47 | 0.002472 | 40 | 0.535187498 |
| | percOf_Live_CD4EMRA | -0.41 | 0.008632 | 40 | 0.532803297 |
| | percOf_Live_CD8EMRA | -0.43 | 0.005478 | 40 | 0.906272113 |
| | umap_component2 | -0.4 | 0.01127 | 39 | 0.513113445 |
| Proteins | CAM_P01033 Metalloproteinase inhibitor 1 (TIMP1) | -0.4 | 0.000697 | 67 | 0.000118445 |
| | CVD2_P05231 Interleukin-6 (IL6) | -0.44 | 0.000172 | 67 | 0.00810288 |
| | CVD2_P07711 Cathepsin L1 (CTSL1) | -0.45 | 0.000145 | 67 | 8.22E-07 |
| | CVD2_P09237 Matrix metalloproteinase-7 | -0.44 | 0.00 | 67 | 0.032766248 |

| | | | | | |
|---|---|---|---|---|---|
| | (MMP-7) | | 023<br>4 | | |
| | CVD3_O00300 Osteoprotegerin (OPG) | -0.41 | 0.00<br>049<br>9 | 67 | 0.000215128 |
| | CVD3_P20160 Azurocidin (AZU1 | -0.42 | 0.00<br>047<br>6 | 67 | 7.12E-06 |
| | DEV_Q96QR1 Secretoglobin family 3A member 1 (SCGB3A1) | 0.4 | 0.00<br>067<br>9 | 67 | 0.000262582 |
| | INF_O00300 Osteoprotegerin (OPG) | -0.43 | 0.00<br>028<br>4 | 67 | 0.000558337 |
| | INF_P05231 Interleukin-6 (IL6) | -0.44 | 0.00<br>020<br>4 | 67 | 0.007025703 |
| | INF_P10145 Interleukin-8 (IL-8) | -0.42 | 0.00<br>037<br>2 | 67 | 0.00036114 |
| | INF_P14210 Hepatocyte growth factor (HGF) | -0.42 | 0.00<br>042<br>1 | 67 | 7.29E-07 |
| | INF_P15018 Leukemia inhibitory factor (LIF) | -0.57 | 3.74<br>E-<br>07 | 67 | 0.000706887 |
| | INF_P55773 C-C motif chemokine 23 (CCL23) | -0.49 | 2.58<br>E-<br>05 | 67 | 7.12E-06 |
| | INF_P78556 C-C motif chemokine 20 (CCL20) | -0.51 | 9.28<br>E-<br>06 | 67 | 0.000218196 |
| | INF_Q13007 Interleukin-24 (IL-24) | -0.42 | 0.00<br>034<br>5 | 67 | 7.42E-05 |
| | IRE_P05231 Interleukin-6 (IL6) | -0.43 | 0.00<br>030<br>1 | 67 | 0.00956383 |
| | ODA_P01258 Calcitonin (CALCA) | -0.41 | 0.00<br>049<br>1 | 67 | 7.10E-06 |
| sPLA2 | P14555 sPLA2 | -0.43 | 0.00<br>822<br>5 | 36 | 0.133848614 |
| PC O-30:0 | | | | | |

| | | | | | |
|---|---|---|---|---|---|
| Immunotype | percOf_Bcell_EOMESpos | -0.4 | 0.01035 | 40 | 0.050295441 |
| | percOf_CD4cTfh_CD4acTfh | -0.48 | 0.001567 | 40 | 0.025195859 |
| | percOf_CD4EM1_HLADRposCD38pos | -0.45 | 0.003399 | 40 | 0.000783423 |
| | percOf_CD4EM1_KI67pos | -0.42 | 0.00758 | 40 | 0.026755803 |
| | percOf_CD8EM1_HLADRposCD38pos | -0.43 | 0.005366 | 40 | 0.012293841 |
| | percOf_CD8EM1_KI67pos | -0.43 | 0.005084 | 40 | 0.114591817 |
| | percOf_CD8RAposCD27posR7neg_KI67pos | -0.42 | 0.006808 | 40 | 0.025674422 |
| | percOf_CD8RAposCD27posR7posCD95pos_HLADRposCD38pos | -0.46 | 0.002833 | 40 | 0.037329622 |
| | percOf_CD8RAposCD27posR7posCD95pos_KI67pos | -0.56 | 0.000166 | 40 | 0.031992847 |
| | percOf_Live_CD4EM2 | 0.41 | 0.008612 | 40 | 0.041352462 |
| | percOf_Live_CD4nonNaive | 0.4 | 0.010389 | 40 | 1.48E-07 |
| | umap_component1 | -0.41 | 0.009828 | 39 | 0.000317432 |
| Protein | CAM_Q9BXR6 Complement factor H-related protein 5 (CFHR5) | -0.45 | 0.000119 | 67 | 0.549657525 |
| | CVD3_P01130 Low-density lipoprotein receptor (LDL receptor) | -0.42 | 0.000461 | 67 | 0.000462016 |
| | CVD3_P20160 Azurocidin (AZU1 | -0.4 | 0.000744 | 67 | 7.12E-06 |
| | INF_P13232 Interleukin-7 (IL-7) | -0.48 | 4.24E-05 | 67 | 7.46E-05 |

| | | | | | |
|---|---|---|---|---|---|
| | INF_P14210 Hepatocyte growth factor (HGF) | -0.42 | 0.000404 | 67 | 7.29E-07 |
| | INF_P80511 Protein S100-A12 (EN-RAGE ) | -0.48 | 3.55E-05 | 67 | 1.21E-08 |
| | IRE_P78362 SRSF protein kinase 2 (SRPK2) | -0.4 | 0.000832 | 67 | 0.000194136 |
| | IRE_Q05516 Zinc finger and BTB domain-containing protein 16 (ZBTB16) | -0.4 | 0.000734 | 67 | 2.36E-05 |
| | IRE_Q9GZT9 Egl nine homolog 1 (EGLN1) | -0.41 | 0.00059 | 67 | 0.031265046 |
| | MET_P27695 DNA-(apurinic or apyrimidinic site) lyase (APEX1) | -0.46 | 8.20E-05 | 67 | 0.000162327 |
| | MET_Q02790 Peptidyl-prolyl cis-trans isomerase FKBP4 (FKBP4) | -0.41 | 0.000511 | 67 | 0.001680452 |
| | ODA_O60934 Nibrin (NBN) | -0.42 | 0.00041 | 67 | 0.001934367 |
| | ODA_P09769 Tyrosine-protein kinase Fgr (FGR) | -0.47 | 5.65E-05 | 67 | 2.10E-06 |
| | ODA_Q02880 DNA topoisomerase 2-beta (TOP2B) | -0.41 | 0.000603 | 67 | 0.000215128 |
| | ODA_Q9HAW4 Claspin (CLSPN) | -0.45 | 0.000155 | 67 | 0.001265811 |
| ChoE-18:3 | | | | | |
| Immunotype | percOf_Bcell_CD138pos | -0.45 | 0.003429 | 40 | 0.062190734 |
| | percOf_Bcell_CD39pos | -0.46 | 0.002833 | 40 | 0.114591817 |
| | percOf_Bcell_HLADRpos | 0.45 | 0.003908 | 40 | 0.009618759 |
| | percOf_Bcell_KI67pos | -0.43 | 0.005074 | 40 | 0.105547477 |
| | percOf_BcellnotPB_CD138pos | -0.41 | 0.00 | 40 | 0.982727613 |

| | | | | | |
|---|---|---|---|---|---|
| | | | 8247 | | |
| Proteins | CVD2_O00182 Galectin-9 (Gal-9) | -0.4 | 0.000925 | 67 | 0.000460426 |
| | CVD2_Q9NQ25 SLAM family member 7 (SLAMF7) | -0.44 | 0.00017 | 67 | 0.945889856 |
| | CVD3_P35247 Pulmonary surfactant-associated protein D (PSP-D) | -0.41 | 0.000488 | 67 | 7.60E-05 |
| | DEV_Q96RD9 Fc receptor-like protein 5 (FCRL5) | -0.44 | 0.000227 | 67 | 0.211300783 |
| | DEV_Q9NZV1 Cysteine-rich motor neuron 1 protein (CRIM1) | -0.41 | 0.000647 | 67 | 0.040625923 |
| | DEV_Q9Y279 V-set and immunoglobulin domain-containing protein 4 (VSIG4) | -0.4 | 0.000686 | 67 | 2.47E-06 |
| | INF_Q5T4W7 Artemin (ARTN) | -0.42 | 0.000412 | 67 | 0.042728343 |
| | INF_Q9NZQ7 Programmed cell death 1 ligand 1 (PD-L1) | -0.43 | 0.000331 | 67 | 0.001304588 |
| | IRE_P27540 Aryl hydrocarbon receptor nuclear translocator (ARNT) | -0.43 | 0.000287 | 67 | 0.598459965 |
| | IRE_Q8IU57 Interferon lambda receptor 1 (IFNLR1) | -0.44 | 0.000222 | 67 | 0.050771359 |
| | N/A_Q9UNK4 PLA2G2D | -0.56 | 0.048729 | 13 | 0.917641149 |

**Supplementary Table 8:** Functional enrichment analysis of biological processes from proteins significantly associated (Spearman's rho < -0.4 or > 0.4, p < 0.05) with LPC-O-16:0, PC-O-30:0 and ChoE-18:3

| Goterm | term description | observed gene count | background gene count | strength | false discovery rate | Observed proteins |
|---|---|---|---|---|---|---|
| LPC-O-16:0 | | | | | | |
| GO:0060326 | Cell chemotaxis | 7 | 204 | 1.68 | 6.32 E-07 | HGF,AZU1,CXCL8,CALCA,CCL20,IL6,CCL23 |
| GO:0019221 | Cytokine-mediated signaling pathway | 9 | 678 | 1.27 | 7.71 E-07 | TIMP1,HGF,LIF,TNFRSF11B,CXCL8,CCL20,IL24,IL6,CCL23 |
| GO:0071345 | Cellular response to cytokine stimulus | 10 | 1013 | 1.14 | 7.71 E-07 | TIMP1,HGF,LIF,TNFRSF11B,CXCL8,CALCA,CCL20,IL24,IL6,CCL23 |
| GO:0097529 | Myeloid leukocyte migration | 6 | 123 | 1.83 | 7.71 E-07 | AZU1,CXCL8,CALCA,CCL20,IL6,CCL23 |
| GO:0030595 | Leukocyte chemotaxis | 6 | 142 | 1.77 | 1.03 E-06 | AZU1,CXCL8,CALCA,CCL20,IL6,CCL23 |
| GO:0070887 | Cellular response to chemical stimulus | 12 | 2919 | 0.76 | 1.54 E-05 | TIMP1,HGF,AZU1,LIF,TNFRSF11B,CXCL8,CALCA,CTSL,CCL20,IL24,IL6,CCL23 |
| GO:0071310 | Cellular response to organic substance | 11 | 2369 | 0.81 | 3.45 E-05 | TIMP1,HGF,LIF,TNFRSF11B,CXCL8,CALCA,CTSL,CCL20,IL24,IL6,CCL23 |
| GO:0002548 | Monocyte chemotaxis | 4 | 43 | 2.11 | 3.83 E-05 | CALCA,CCL20,IL6,CCL23 |
| GO:0006954 | Inflammatory response | 7 | 515 | 1.28 | 3.83 E-05 | TIMP1,AZU1,CXCL8,CALCA,CCL20,IL6,CCL23 |

| | | | | | | |
|---|---|---|---|---|---|---|
| GO:0001934 | Positive regulation of protein phosphorylation | 8 | 1019 | 1.04 | 0.00011 | HGF,AZU1,LIF,CALCA,CCL20,IL24,IL6,CCL23 |
| GO:0007165 | Signal transduction | 13 | 4876 | 0.57 | 0.00013 | TIMP1,HGF,AZU1,LIF,SCGB3A1,TNFRSF11B,CXCL8,CALCA,CTSL,CCL20,IL24,IL6,CCL23 |
| GO:0045779 | Negative regulation of bone resorption | 3 | 14 | 2.48 | 0.00015 | TNFRSF11B,CALCA,IL6 |
| GO:0051716 | Cellular response to stimulus | 14 | 6489 | 0.48 | 0.00015 | TIMP1,HGF,AZU1,LIF,MMP7,SCGB3A1,TNFRSF11B,CXCL8,CALCA,CTSL,CCL20,IL24,IL6,CCL23 |
| GO:0009605 | Response to external stimulus | 10 | 2310 | 0.78 | 0.0002 | HGF,AZU1,MMP7,TNFRSF11B,CXCL8,CALCA,CCL20,IL24,IL6,CCL23 |
| GO:0071356 | Cellular response to tumor necrosis factor | 5 | 245 | 1.46 | 0.00028 | TNFRSF11B,CXCL8,CALCA,CCL20,CCL23 |
| GO:0006959 | Humoral immune response | 5 | 275 | 1.41 | 0.00042 | AZU1,CXCL8,CALCA,CCL20,IL6 |
| GO:0040012 | Regulation of locomotion | 7 | 969 | 1 | 0.00071 | TIMP1,HGF,AZU1,CXCL8,CCL20,IL24,IL6 |
| GO:0009617 | Response to bacterium | 6 | 634 | 1.12 | 0.00097 | AZU1,CXCL8,CALCA,CCL20,IL24,IL6 |
| GO:0065009 | Regulation of molecular function | 12 | 4913 | 0.53 | 0.0011 | TIMP1,HGF,AZU1,LIF,SCGB3A1,TNFRSF11B,CXCL8,CALCA,CCL20,IL24,IL6,CCL23 |
| GO:0006955 | Immune response | 8 | 1588 | 0.85 | 0.0012 | AZU1,LIF,CXCL8,CALCA,CTSL,CCL20,IL6,CCL23 |
| GO:0019730 | Antimicrobial humoral response | 4 | 160 | 1.54 | 0.0013 | AZU1,CXCL8,CALCA,CCL20 |
| GO:0 | Myeloid | 2 | 5 | 2.75 | 0.00 | CTSL,IL6 |

| | | | | | | |
|---|---|---|---|---|---|---|
| 033028 | cell apoptotic process | | | | 26 | |
| GO:0050731 | Positive regulation of peptidyl-tyrosine phosphorylation | 4 | 196 | 1.46 | 0.0026 | HGF,LIF,IL24,IL6 |
| GO:0051707 | Response to other organism | 7 | 1256 | 0.89 | 0.0026 | AZU1,CXCL8,CALCA,CCL20,IL24,IL6,CCL23 |
| GO:0006925 | Inflammatory cell apoptotic process | 2 | 6 | 2.67 | 0.0029 | CTSL,IL6 |
| GO:0022617 | Extracellular matrix disassembly | 3 | 66 | 1.8 | 0.0032 | TIMP1,MMP7,CTSL |
| GO:0030334 | Regulation of cell migration | 6 | 865 | 0.99 | 0.0034 | TIMP1,HGF,CXCL8,CCL20,IL24,IL6 |
| GO:0042531 | Positive regulation of tyrosine phosphorylation of stat protein | 3 | 68 | 1.79 | 0.0034 | LIF,IL24,IL6 |
| GO:0050793 | Regulation of developmental process | 9 | 2648 | 0.68 | 0.0034 | TIMP1,HGF,LIF,SCGB3A1,TNFRSF11B,CXCL8,CALCA,CTSL,IL6 |
| GO:0032268 | Regulation of cellular protein metabolic process | 9 | 2693 | 0.67 | 0.0037 | TIMP1,HGF,AZU1,LIF,CALCA,CCL20,IL24,IL6,CCL23 |
| GO:0030593 | Neutrophil chemotaxis | 3 | 74 | 1.75 | 0.0041 | CXCL8,CCL20,CCL23 |
| GO:0 | Regulation | 9 | 2740 | 0.66 | 0.00 | TIMP1,HGF,AZU1,LIF,CXCL8,CALCA,CCL20,IL2 |

| | | | | | | |
|---|---|---|---|---|---|---|
| 032879 | of localization | | | | 42 | 4,IL6 |
| GO:0070098 | Chemokine-mediated signaling pathway | 3 | 80 | 1.72 | 0.0047 | CXCL8,CCL20,CCL23 |
| GO:0043410 | Positive regulation of mapk cascade | 5 | 543 | 1.11 | 0.0049 | HGF,LIF,CCL20,IL6,CCL23 |
| GO:2000026 | Regulation of multicellular organismal development | 8 | 2096 | 0.73 | 0.0049 | TIMP1,HGF,LIF,TNFRSF11B,CXCL8,CALCA,CTSL,IL6 |
| GO:0040017 | Positive regulation of locomotion | 5 | 562 | 1.09 | 0.0053 | HGF,AZU1,CXCL8,CCL20,IL6 |
| GO:0031640 | Killing of cells of other organism | 3 | 91 | 1.66 | 0.0058 | AZU1,CXCL8,CCL20 |
| GO:0042742 | Defense response to bacterium | 4 | 277 | 1.3 | 0.0058 | AZU1,CALCA,CCL20,IL6 |
| GO:0050829 | Defense response to gram-negative bacterium | 3 | 98 | 1.63 | 0.0069 | AZU1,CALCA,IL6 |
| GO:0048584 | Positive regulation of response to stimulus | 8 | 2257 | 0.69 | 0.0073 | HGF,AZU1,LIF,CXCL8,CCL20,IL24,IL6,CCL23 |
| GO:0009966 | Regulation of signal transduction | 9 | 3107 | 0.61 | 0.0086 | TIMP1,HGF,LIF,CXCL8,CALCA,CCL20,IL24,IL6,CCL23 |
| GO:0042127 | Regulation of cell population | 7 | 1642 | 0.78 | 0.0086 | TIMP1,LIF,SCGB3A1,CXCL8,IL24,IL6,CCL23 |

| | | | | | | |
|---|---|---|---|---|---|---|
| | proliferation | | | | | |
| GO:0022603 | Regulation of anatomical structure morphogenesis | 6 | 1095 | 0.88 | 0.0089 | HGF,LIF,SCGB3A1,TNFRSF11B,CXCL8,IL6 |
| GO:0045651 | Positive regulation of macrophage differentiation | 2 | 15 | 2.27 | 0.009 | LIF,CALCA |
| GO:0050930 | Induction of positive chemotaxis | 2 | 15 | 2.27 | 0.009 | AZU1,CXCL8 |
| GO:0061844 | Antimicrobial humoral immune response mediated by antimicrobial peptide | 3 | 113 | 1.57 | 0.0096 | CXCL8,CALCA,CCL20 |
| GO:0048583 | Regulation of response to stimulus | 10 | 4114 | 0.53 | 0.0099 | TIMP1,HGF,AZU1,LIF,CXCL8,CALCA,CCL20,IL24,IL6,CCL23 |
| GO:0030198 | Extracellular matrix organization | 4 | 338 | 1.22 | 0.0107 | TIMP1,MMP7,TNFRSF11B,CTSL |
| GO:0051239 | Regulation of multicellular organismal process | 9 | 3227 | 0.59 | 0.0107 | TIMP1,HGF,AZU1,LIF,TNFRSF11B,CXCL8,CALCA,CTSL,IL6 |
| GO:0008285 | Negative regulation of cell population proliferatio | 5 | 696 | 1 | 0.0115 | LIF,CXCL8,IL24,IL6,CCL23 |

| | | | | | | |
|---|---|---|---|---|---|---|
| | n | | | | | |
| GO:0019932 | Second-messenger-mediated signaling | 4 | 354 | 1.2 | 0.0122 | AZU1,CXCL8,CALCA,CCL20 |
| GO:0048519 | Negative regulation of biological process | 11 | 5389 | 0.46 | 0.0133 | TIMP1,HGF,AZU1,LIF,SCGB3A1,TNFRSF11B,CXCL8,CALCA,IL24,IL6,CCL23 |
| GO:0002687 | Positive regulation of leukocyte migration | 3 | 144 | 1.46 | 0.0169 | CXCL8,CCL20,IL6 |
| GO:0033135 | Regulation of peptidyl-serine phosphorylation | 3 | 146 | 1.46 | 0.0174 | HGF,LIF,IL6 |
| GO:0035584 | Calcium-mediated signaling using intracellular calcium source | 2 | 24 | 2.07 | 0.0175 | AZU1,CCL20 |
| GO:0048522 | Positive regulation of cellular process | 11 | 5579 | 0.44 | 0.0175 | TIMP1,HGF,AZU1,LIF,SCGB3A1,CXCL8,CALCA,CCL20,IL24,IL6,CCL23 |
| GO:0050921 | Positive regulation of chemotaxis | 3 | 147 | 1.46 | 0.0175 | AZU1,CXCL8,IL6 |
| GO:0071887 | Leukocyte apoptotic process | 2 | 26 | 2.03 | 0.0198 | CTSL,IL6 |
| GO:0048710 | Regulation of astrocyte differentiation | 2 | 27 | 2.02 | 0.0208 | LIF,IL6 |
| GO:0 | Calcium- | 3 | 165 | 1.41 | 0.02 | AZU1,CXCL8,CCL20 |

| | | | | | | |
|---|---|---|---|---|---|---|
| 019722 | mediated signaling | | | | 26 | |
| GO:0051094 | Positive regulation of developmental process | 6 | 1389 | 0.78 | 0.0251 | HGF,LIF,SCGB3A1,CXCL8,CALCA,IL6 |
| GO:0071347 | Cellular response to interleukin-1 | 3 | 174 | 1.38 | 0.0259 | CXCL8,CCL20,CCL23 |
| GO:0001819 | Positive regulation of cytokine production | 4 | 461 | 1.08 | 0.0272 | HGF,AZU1,CALCA,IL6 |
| GO:0007565 | Female pregnancy | 3 | 183 | 1.36 | 0.0295 | LIF,MMP7,CALCA |
| GO:0009719 | Response to endogenous stimulus | 6 | 1447 | 0.76 | 0.0302 | TIMP1,TNFRSF11B,CXCL8,CALCA,CTSL,IL6 |
| GO:0071222 | Cellular response to lipopolysaccharide | 3 | 185 | 1.36 | 0.0302 | CXCL8,IL24,IL6 |
| GO:0098542 | Defense response to other organism | 5 | 900 | 0.89 | 0.0302 | AZU1,CALCA,CCL20,IL6,CCL23 |
| GO:0048523 | Negative regulation of cellular process | 10 | 4874 | 0.46 | 0.0335 | TIMP1,HGF,AZU1,LIF,SCGB3A1,CXCL8,CALCA,IL24,IL6,CCL23 |
| GO:0030278 | Regulation of ossification | 3 | 197 | 1.33 | 0.0348 | HGF,CALCA,IL6 |
| GO:0002684 | Positive regulation of immune system process | 5 | 949 | 0.87 | 0.0363 | LIF,CXCL8,CALCA,CCL20,IL6 |

| | | | | | | |
|---|---|---|---|---|---|---|
| GO:0007566 | Embryo implantation | 2 | 41 | 1.83 | 0.0399 | LIF,CALCA |
| GO:0030335 | Positive regulation of cell migration | 4 | 522 | 1.03 | 0.0399 | HGF,CXCL8,CCL20,IL6 |
| GO:0030574 | Collagen catabolic process | 2 | 43 | 1.81 | 0.043 | MMP7,CTSL |
| GO:0045597 | Positive regulation of cell differentiation | 5 | 993 | 0.85 | 0.043 | HGF,LIF,SCGB3A1,CALCA,IL6 |
| GO:0045744 | Negative regulation of g protein-coupled receptor signaling pathway | 2 | 45 | 1.79 | 0.0455 | CXCL8,CALCA |
| PC-O-30:0 | | | | | | |
| GO:0010604 | Positive regulation of macromolecule metabolic process | 12 | 3600 | 0.64 | 0.0051 | APEX1,HGF,AZU1,IL7,NBN,CLSPN,ZBTB16,EGLN1,S100A12,FGR,SRPK2,LDLR |
| GO:0048518 | Positive regulation of biological process | 14 | 6112 | 0.48 | 0.0058 | APEX1,HGF,AZU1,CFHR5,IL7,NBN,CLSPN,ZBTB16,EGLN1,S100A12,FGR,SRPK2,TOP2B,LDLR |
| GO:0048519 | Negative regulation of biological | 13 | 5389 | 0.5 | 0.0099 | FKBP4,APEX1,HGF,AZU1,IL7,NBN,CLSPN,ZBTB16,EGLN1,FGR,SRPK2,TOP2B,LDLR |

| | | | | | | |
|---|---|---|---|---|---|---|
| | process | | | | | |
| GO:0033674 | Positive regulation of kinase activity | 6 | 624 | 1.1 | 0.0109 | HGF,AZU1,NBN,CLSPN,S100A12,FGR |
| GO:0043170 | Macromolecule metabolic process | 13 | 6137 | 0.44 | 0.0278 | FKBP4,APEX1,HGF,AZU1,NBN,CLSPN,ZBTB16,EGLN1,S100A12,FGR,SRPK2,TOP2B,LDLR |
| GO:0019220 | Regulation of phosphate metabolic process | 8 | 1816 | 0.76 | 0.0317 | HGF,AZU1,IL7,NBN,CLSPN,S100A12,FGR,LDLR |
| GO:0031325 | Positive regulation of cellular metabolic process | 10 | 3413 | 0.58 | 0.043 | APEX1,HGF,AZU1,NBN,CLSPN,ZBTB16,EGLN1,S100A12,FGR,LDLR |
| GO:0001932 | Regulation of protein phosphorylation | 7 | 1459 | 0.8 | 0.044 | HGF,AZU1,IL7,NBN,CLSPN,S100A12,FGR |
| GO:0006950 | Response to stress | 10 | 3485 | 0.57 | 0.044 | APEX1,AZU1,CFHR5,NBN,CLSPN,EGLN1,S100A12,FGR,SRPK2,LDLR |
| GO:0006959 | Humoral immune response | 4 | 275 | 1.28 | 0.044 | AZU1,CFHR5,IL7,S100A12 |
| GO:0031640 | Killing of cells of other organism | 3 | 91 | 1.63 | 0.044 | AZU1,CFHR5,S100A12 |
| GO:0048522 | Positive regulation of cellular process | 12 | 5579 | 0.45 | 0.044 | APEX1,HGF,AZU1,IL7,NBN,CLSPN,ZBTB16,EGLN1,S100A12,FGR,SRPK2,LDLR |
| GO:0048731 | System development | 11 | 4426 | 0.51 | 0.044 | FKBP4,HGF,AZU1,IL7,NBN,ZBTB16,EGLN1,FGR,SRPK2,TOP2B,LDLR |
| GO:0051246 | Regulation of protein metabolic | 9 | 2828 | 0.62 | 0.044 | HGF,AZU1,IL7,NBN,CLSPN,EGLN1,S100A12,FGR,LDLR |

| | | | | | | |
|---|---|---|---|---|---|---|
| | process | | | | | |
| GO:0009653 | Anatomical structure morphogenesis | 8 | 2165 | 0.68 | 0.0481 | HGF,IL7,ZBTB16,EGLN1,FGR,SRPK2,TOP2B,LDLR |
| GO:0044774 | Mitotic DNA integrity checkpoint | 3 | 109 | 1.56 | 0.0482 | NBN,CLSPN,TOP2B |
| GO:0006955 | Immune response | 7 | 1588 | 0.76 | 0.0491 | AZU1,CFHR5,IL7,NBN,S100A12,FGR,SRPK2 |
| GO:0032502 | Developmental process | 12 | 5841 | 0.43 | 0.0491 | FKBP4,APEX1,HGF,AZU1,IL7,NBN,ZBTB16,EGLN1,FGR,SRPK2,TOP2B,LDLR |
| GO:0051171 | Regulation of nitrogen compound metabolic process | 12 | 5836 | 0.43 | 0.0491 | APEX1,HGF,AZU1,IL7,NBN,CLSPN,ZBTB16,EGLN1,S100A12,FGR,SRPK2,LDLR |
| ChoE-18:3 | | | | | | |
| GO:0042130 | Negative regulation of T cell proliferation | 4 | 66 | 2.07 | 0.0004 | SFTPD,VSIG4,CD274,LGALS9 |
| GO:0002682 | Regulation of immune system process | 7 | 1514 | 0.96 | 0.0022 | IFNLR1,ARNT,SLAMF7,SFTPD,VSIG4,CD274,LGALS9 |
| GO:0001818 | Negative regulation of cytokine production | 4 | 280 | 1.45 | 0.0068 | SFTPD,VSIG4,CD274,LGALS9 |
| GO:2000562 | Negative regulation of CD4-positive, alpha-beta T cell proliferatio | 2 | 8 | 2.69 | 0.0075 | CD274,LGALS9 |

| | | | | | | |
|---|---|---|---|---|---|---|
| | n | | | | | |
| GO:0008285 | Negative regulation of cell population proliferation | 5 | 696 | 1.15 | 0.008 | IFNLR1,SFTPD,VSIG4,CD274,LGALS9 |
| GO:0045087 | Innate immune response | 5 | 703 | 1.14 | 0.008 | IFNLR1,SLAMF7,SFTPD,VSIG4,LGALS9 |
| GO:0051707 | Response to other organism | 6 | 1256 | 0.97 | 0.008 | IFNLR1,SLAMF7,SFTPD,VSIG4,CD274,LGALS9 |
| GO:0001817 | Regulation of cytokine production | 5 | 742 | 1.12 | 0.0082 | ARNT,SFTPD,VSIG4,CD274,LGALS9 |
| GO:0046007 | Negative regulation of activated T cell proliferation | 2 | 13 | 2.48 | 0.0109 | CD274,LGALS9 |
| GO:0009605 | Response to external stimulus | 7 | 2310 | 0.77 | 0.012 | IFNLR1,SLAMF7,SFTPD,VSIG4,CD274,LGALS9,ARTN |
| GO:0070234 | Positive regulation of T cell apoptotic process | 2 | 15 | 2.42 | 0.0132 | CD274,LGALS9 |
| GO:0002831 | Regulation of response to biotic stimulus | 4 | 406 | 1.29 | 0.0145 | IFNLR1,VSIG4,CD274,LGALS9 |
| GO:0002376 | Immune system process | 7 | 2481 | 0.74 | 0.0165 | IFNLR1,SLAMF7,SFTPD,VSIG4,CD274,LGALS9,ARTN |
| GO:0006955 | Immune response | 6 | 1588 | 0.87 | 0.0165 | IFNLR1,SLAMF7,SFTPD,VSIG4,CD274,LGALS9 |
| GO:0050776 | Regulation of immune response | 5 | 896 | 1.04 | 0.0165 | SLAMF7,SFTPD,VSIG4,CD274,LGALS9 |

| | | | | | | |
|---|---|---|---|---|---|---|
| GO:0042127 | Regulation of cell population proliferation | 6 | 1642 | 0.85 | 0.0172 | IFNLR1,ARNT,SFTPD,VSIG4,CD274,LGALS9 |
| GO:0032703 | Negative regulation of interleukin-2 production | 2 | 23 | 2.23 | 0.0214 | SFTPD,VSIG4 |
| GO:0048583 | Regulation of response to stimulus | 8 | 4114 | 0.58 | 0.0334 | IFNLR1,ARNT,SLAMF7,SFTPD,VSIG4,CD274,LGALS9,ARTN |
| GO:0050896 | Response to stimulus | 10 | 8046 | 0.39 | 0.0357 | CRIM1,IFNLR1,ARNT,FCRL5,SLAMF7,SFTPD,VSIG4,CD274,LGALS9,ARTN |
| GO:0032689 | Negative regulation of interferon-gamma production | 2 | 35 | 2.05 | 0.039 | CD274,LGALS9 |
| GO:0032733 | Positive regulation of interleukin-10 production | 2 | 39 | 2 | 0.0462 | CD274,LGALS9 |

**Supplementary table 9**: Tissue/cell type enrichment analysis of proteins associated to LPC-O-16:0 (refer to supplementary table S7

| #term ID | term description | observed gene count | background gene count | strength | false discovery rate | matching proteins in your network (IDs | |
|---|---|---|---|---|---|---|---|
| BTO:0000519 | Gingiva | 2 | 5 | 2.75 | 0.0135 | 9606.ENSP00000306512,9606.ENSP00000385675 | CXCL8,IL6 |
| BTO:0001370 | THP-1 cell | 2 | 4 | 2.84 | 0.0135 | 9606.ENSP00000306512,9606.ENSP00000385675 | CXCL8,IL6 |
| BTO:0003861 | Inflammatory cell | 2 | 7 | 2.6 | 0.0135 | 9606.ENSP00000306512,9606.ENSP00000385675 | CXCL8,IL6 |
| BTO:0005265 | Nonparenchymal liver cell | 2 | 9 | 2.49 | 0.0135 | 9606.ENSP00000218388,9606.ENSP00000385675 | TIMP1,IL6 |
| BTO:0000473 | Fetal membrane | 2 | 14 | 2.3 | 0.0157 | 9606.ENSP00000306512,9606.ENSP00000385675 | CXCL8,IL6 |
| BTO:0001539 | Parenchyma | 2 | 14 | 2.3 | 0.0157 | 9606.ENSP00000218388,9606.ENSP00000385675 | TIMP1,IL6 |
| BTO:0000878 | Mononuclear cell | 3 | 117 | 1.55 | 0.0165 | 9606.ENSP00000306512,9606.ENSP00000385675,9606.ENSP00000481357 | CXCL8,IL6,CCL23 |
| BTO:0001044 | Phagocyte | 3 | 117 | 1.55 | 0.0165 | 9606.ENSP00000233997,9606.ENSP00000306512,9606.ENSP00000481357 | AZU1,CXCL8,CCL23 |
| BTO:0000130 | Neutrophil | 2 | 34 | 1.91 | 0.0468 | 9606.ENSP00000233997,9606.ENSP00000306512 | AZU1,CXCL8 |

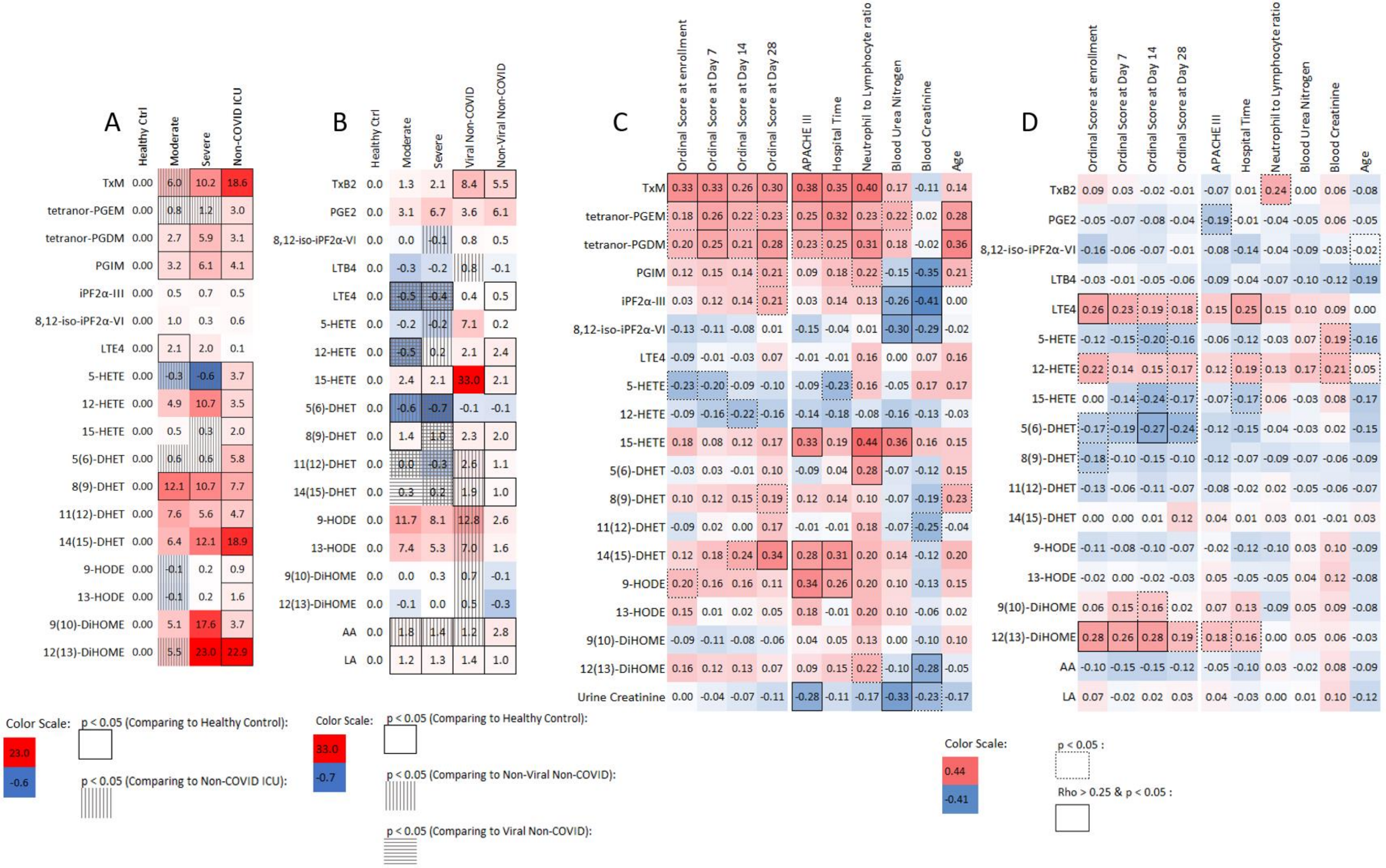
A
Healthy Ctrl | Moderate | Severe | Non-COVID ICU
TxM 0.00 6.0 10.2 18.6
tetranor-PGEM 0.00 0.8 1.2 3.0
tetranor-PGDM 0.00 2.7 5.9 3.1
PGIM 0.00 3.2 6.1 4.1
iPF2α-III 0.00 0.5 0.7 0.5
8,12-iso-iPF2α-VI 0.00 1.0 0.3 0.6
LTE4 0.00 2.1 2.0 0.1
5-HETE 0.00 -0.3 -0.6 3.7
12-HETE 0.00 4.9 10.7 3.5
15-HETE 0.00 0.5 0.3 2.0
5(6)-DHET 0.00 0.6 0.6 5.8
8(9)-DHET 0.00 12.1 10.7 7.7
11(12)-DHET 0.00 7.6 5.6 4.7
14(15)-DHET 0.00 6.4 12.1 18.9
9-HODE 0.00 -0.1 0.2 0.9
13-HODE 0.00 -0.1 0.2 1.6
9(10)-DiHOME 0.00 5.1 17.6 3.7
12(13)-DiHOME 0.00 5.5 23.0 22.9
Color Scale: 23.0 -0.6
p < 0.05 (Comparing to Healthy Control):
p < 0.05 (Comparing to Non-COVID ICU):
B
Healthy Ctrl | Moderate | Severe | Viral Non-COVID | Non-Viral Non-COVID
TxB2 0.0 1.3 2.1 8.4 5.5
PGE2 0.0 3.1 6.7 3.6 6.1
8,12-iso-iPF2α-VI 0.0 0.0 -0.1 0.8 0.5
LTB4 0.0 -0.3 -0.2 0.8 -0.1
LTE4 0.0 -0.5 -0.4 0.4 0.5
5-HETE 0.0 -0.2 -0.2 7.1 0.2
12-HETE 0.0 -0.5 0.2 2.1 2.4
15-HETE 0.0 2.4 2.1 33.0 2.1
5(6)-DHET 0.0 -0.6 -0.7 -0.1 -0.1
8(9)-DHET 0.0 1.4 1.0 2.3 2.0
11(12)-DHET 0.0 0.0 -0.3 2.6 1.1
14(15)-DHET 0.0 0.3 0.2 1.9 1.0
9-HODE 0.0 11.7 8.1 12.8 2.6
13-HODE 0.0 7.4 5.3 7.0 1.6
9(10)-DiHOME 0.0 0.0 0.3 0.7 -0.1
12(13)-DiHOME 0.0 -0.1 0.0 0.5 -0.3
AA 0.0 1.8 1.4 1.2 2.8
LA 0.0 1.2 1.3 1.4 1.0
Color Scale: 33.0 -0.7
p < 0.05 (Comparing to Healthy Control):
p < 0.05 (Comparing to Non-Viral Non-COVID):
p < 0.05 (Comparing to Viral Non-COVID):
C
Ordinal Score at enrollment | Ordinal Score at Day 7 | Ordinal Score at Day 14 | Ordinal Score at Day 28 | APACHE III | Hospital Time | Neutrophil to Lymphocyte ratio | Blood Urea Nitrogen | Blood Creatinine | Age
TxM 0.33 0.33 0.26 0.30 0.38 0.35 0.40 0.17 -0.11 0.14
tetranor-PGEM 0.18 0.26 0.22 0.23 0.25 0.32 0.23 0.22 0.02 0.28
tetranor-PGDM 0.20 0.25 0.21 0.28 0.23 0.25 0.31 0.18 -0.02 0.36
PGIM 0.12 0.15 0.14 0.21 0.09 0.18 0.22 -0.15 -0.35 0.21
iPF2α-III 0.03 0.12 0.14 0.21 0.03 0.14 0.13 -0.26 -0.41 0.00
8,12-iso-iPF2α-VI -0.13 -0.11 -0.08 0.01 -0.15 -0.04 0.01 -0.30 -0.29 -0.02
LTE4 -0.09 -0.01 -0.03 0.07 -0.01 -0.01 0.16 0.00 0.07 0.16
5-HETE -0.23 -0.20 -0.09 -0.10 -0.09 -0.23 0.16 -0.05 0.17 0.17
12-HETE -0.09 -0.16 -0.22 -0.16 -0.14 -0.18 -0.08 -0.16 -0.13 -0.03
15-HETE 0.18 0.08 0.12 0.17 0.33 0.19 0.44 0.36 0.16 0.15
5(6)-DHET -0.03 0.03 -0.01 0.10 -0.09 0.04 0.28 -0.07 -0.12 0.15
8(9)-DHET 0.10 0.12 0.15 0.19 0.12 0.14 0.10 -0.07 -0.19 0.23
11(12)-DHET -0.09 0.02 0.00 0.17 -0.01 -0.01 0.18 -0.07 -0.25 -0.04
14(15)-DHET 0.12 0.18 0.24 0.34 0.28 0.31 0.20 0.14 -0.12 0.20
9-HODE 0.20 0.16 0.16 0.11 0.34 0.26 0.20 0.10 -0.13 0.15
13-HODE 0.15 0.01 0.02 0.05 0.18 -0.01 0.20 0.10 -0.06 0.02
9(10)-DiHOME -0.09 -0.11 -0.08 -0.06 0.04 0.05 0.13 0.00 -0.10 0.10
12(13)-DiHOME 0.16 0.12 0.13 0.07 0.09 0.15 0.22 -0.10 -0.28 -0.05
Urine Creatinine 0.00 -0.04 -0.07 -0.11 -0.28 -0.11 -0.17 -0.33 -0.23 -0.17
D
Ordinal Score at enrollment | Ordinal Score at Day 7 | Ordinal Score at Day 14 | Ordinal Score at Day 28 | APACHE III | Hospital Time | Neutrophil to Lymphocyte ratio | Blood Urea Nitrogen | Blood Creatinine | Age
TxB2 0.09 0.03 -0.02 -0.01 -0.07 0.01 0.24 0.00 0.06 -0.08
PGE2 -0.05 -0.07 -0.08 -0.04 -0.19 -0.01 -0.04 -0.05 0.06 -0.05
8,12-iso-iPF2α-VI -0.16 -0.06 -0.07 -0.01 -0.08 -0.14 -0.04 -0.09 -0.03 -0.02
LTB4 -0.03 -0.01 -0.05 -0.06 -0.09 -0.04 -0.07 -0.10 -0.12 -0.19
LTE4 0.26 0.23 0.19 0.18 0.15 0.25 0.15 0.10 0.09 0.00
5-HETE -0.12 -0.15 -0.20 -0.16 -0.06 -0.12 -0.03 0.07 0.19 -0.16
12-HETE 0.22 0.14 0.15 0.17 0.12 0.19 0.13 0.17 0.21 0.05
15-HETE 0.00 -0.14 -0.24 -0.17 -0.07 -0.17 0.06 -0.03 0.08 -0.17
5(6)-DHET -0.17 -0.19 -0.27 -0.24 -0.12 -0.15 -0.04 -0.03 0.02 -0.15
8(9)-DHET -0.18 -0.10 -0.15 -0.10 -0.12 -0.07 -0.09 -0.07 -0.06 -0.09
11(12)-DHET -0.13 -0.06 -0.11 -0.07 -0.08 -0.02 0.02 -0.05 -0.06 -0.07
14(15)-DHET 0.00 0.00 0.01 0.12 0.04 0.01 0.03 0.01 -0.01 0.03
9-HODE -0.11 -0.08 -0.10 -0.07 -0.02 -0.12 -0.10 0.03 0.10 -0.09
13-HODE -0.02 0.00 -0.02 -0.03 0.05 -0.05 -0.05 0.04 0.12 -0.08
9(10)-DiHOME 0.06 0.15 0.16 0.02 0.07 0.13 -0.09 0.05 0.09 -0.08
12(13)-DiHOME 0.28 0.26 0.28 0.19 0.18 0.16 0.00 0.05 0.06 -0.03
AA -0.10 -0.15 -0.15 -0.12 -0.05 -0.10 0.03 -0.02 0.08 -0.09
LA 0.07 -0.02 0.02 0.03 0.04 -0.03 0.00 0.01 0.10 -0.12
Color Scale: 0.44 -0.41
p < 0.05 :
Rho > 0.25 & p < 0.05 :

Figure 1

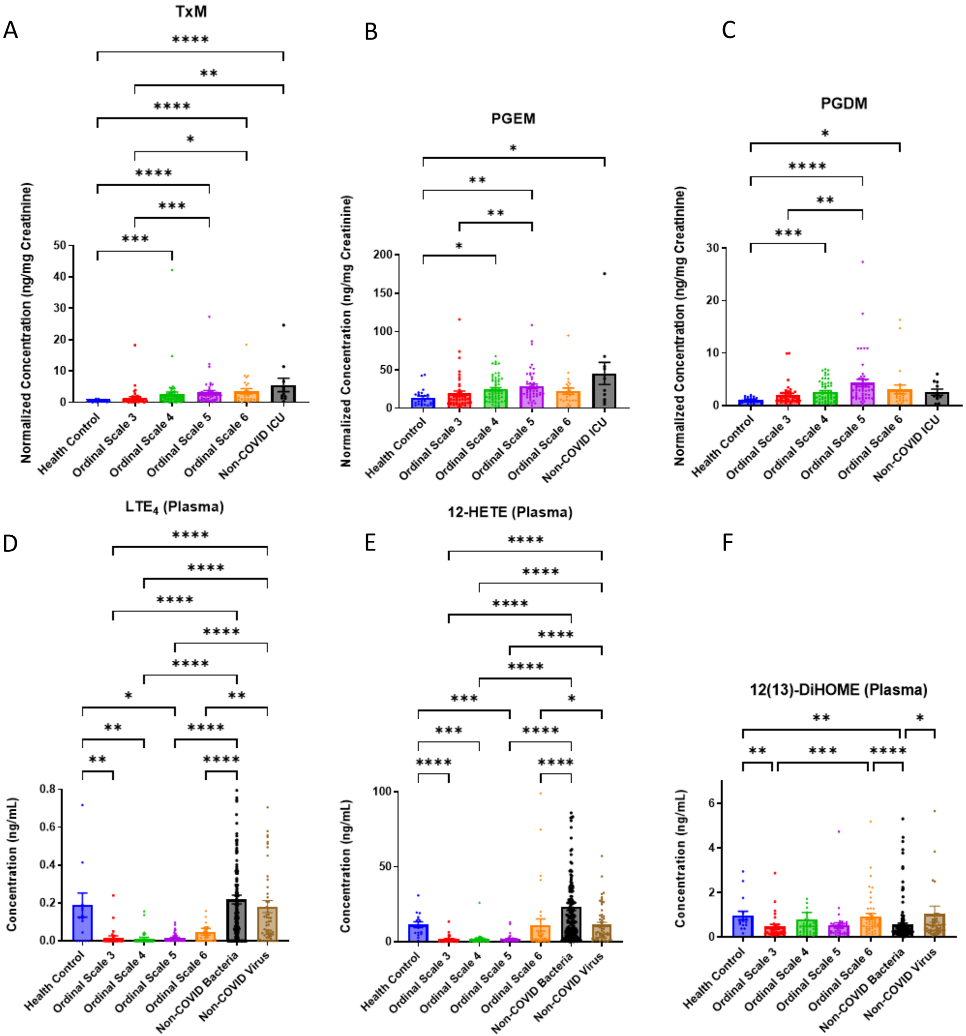

A
TxM
B
PGEM
C
PGDM
D
LTE4 (Plasma)
E
12-HETE (Plasma)
F
12(13)-DiHOME (Plasma)
Normalized Concentration (ng/mg Creatinine)
Concentration (ng/mL)
Health Control
Ordinal Scale 3
Ordinal Scale 4
Ordinal Scale 5
Ordinal Scale 6
Non-COVID ICU
Non-COVID Bacteria
Non-COVID Virus

Figure 2

A

| ρ / Urine | Percentage of Live All non-T non-B CD16+ | Percentage of Live All non-T non-B | Percentage of CD4+ HLA DR+ | Percentage of CD4+ non-Naive HLA DR+ | Percentage of CD4+ HLA DR+ CD38+ | Percentage of CD4+ non-Naive HLA DR+ CD38+ | Percentage of CD4+EM1 HLA DR+ CD38+ | Percentage of CD4+CM HLA DR+ CD38+ | Percentage of CD8+EM1 HLA DR+ CD38+ | Percentage of CD8+RA+ CD27+ R7- HLA DR+ CD38+ | Percentage of CD8+CM HLA DR+ CD38+ | Percentage of CD8+ CM KI67+ | Percentage of CD8+RA+ CD27+ R7+ CD95+ KI67+ | Percentage of B cell not PB | Percentage of B cell PB | Percentage of Live CD8+ RA+ CD27+ R7- | Percentage of B cell CXCR5+ | Percentage of B cell HLA DR+ | Percentage of Live CD4+CM | Percentage of B cell not PB HLA DR+ | Percentage of Live T cell | Percentage of Live CD4+ non-Naive | Percentage of Live CD4+ |
|---|---|---|---|---|---|---|---|---|---|---|---|---|---|---|---|---|---|---|---|---|---|---|---|
| TxM | 0.19 | 0.22 | 0.12 | 0.15 | 0.32 | 0.31 | 0.25 | 0.24 | 0.22 | 0.13 | 0.13 | 0.03 | -0.13 | -0.07 | 0.07 | -0.15 | -0.19 | -0.04 | -0.16 | -0.09 | -0.26 | -0.29 | -0.30 |
| tetranor-PGEM | 0.20 | 0.11 | 0.08 | 0.10 | 0.30 | 0.32 | 0.30 | 0.33 | 0.06 | 0.06 | 0.14 | 0.11 | -0.01 | -0.14 | 0.14 | 0.17 | -0.09 | -0.01 | 0.00 | -0.03 | -0.01 | -0.04 | -0.06 |
| tetranor-PGDM | 0.43 | 0.38 | 0.29 | 0.28 | 0.28 | 0.27 | 0.27 | 0.26 | 0.09 | 0.08 | 0.07 | 0.04 | 0.02 | 0.01 | -0.01 | -0.02 | -0.07 | -0.19 | -0.27 | -0.29 | -0.32 | -0.34 | -0.42 |
| PGIM | 0.12 | 0.17 | -0.13 | -0.13 | -0.06 | 0.02 | 0.09 | -0.04 | -0.05 | -0.08 | -0.07 | -0.02 | -0.04 | 0.01 | -0.01 | -0.04 | 0.06 | -0.14 | -0.07 | -0.23 | -0.22 | -0.20 | -0.15 |
| iPF2a-III | 0.00 | -0.02 | -0.18 | -0.15 | -0.12 | -0.03 | 0.06 | -0.08 | 0.09 | 0.05 | -0.03 | -0.02 | -0.02 | -0.11 | 0.10 | 0.17 | -0.07 | 0.01 | 0.14 | 0.05 | 0.03 | -0.04 | 0.03 |
| 8,12-iso-iPF2a-VI | -0.26 | -0.22 | -0.29 | -0.22 | -0.19 | -0.09 | 0.00 | -0.13 | 0.01 | -0.08 | -0.12 | -0.09 | -0.05 | 0.01 | -0.01 | 0.04 | 0.10 | 0.10 | 0.20 | 0.19 | 0.16 | 0.03 | 0.19 |
| LTE4 | -0.24 | -0.16 | 0.05 | 0.06 | 0.15 | 0.14 | -0.03 | -0.04 | -0.07 | -0.10 | -0.10 | -0.02 | 0.03 | 0.06 | -0.06 | -0.02 | 0.08 | 0.01 | 0.15 | -0.03 | 0.06 | 0.08 | 0.03 |
| 5-HETE | -0.03 | -0.04 | 0.03 | -0.02 | -0.18 | -0.21 | -0.10 | -0.09 | -0.03 | -0.03 | -0.15 | -0.17 | -0.05 | -0.10 | 0.10 | 0.16 | -0.12 | -0.23 | 0.14 | -0.14 | 0.19 | 0.18 | 0.15 |
| 12-HETE | 0.31 | 0.30 | 0.10 | 0.15 | 0.07 | 0.09 | 0.33 | 0.17 | 0.42 | 0.20 | 0.09 | 0.00 | 0.08 | 0.06 | -0.06 | -0.35 | -0.06 | -0.08 | 0.00 | -0.17 | -0.27 | -0.07 | -0.13 |
| 15-HETE | 0.23 | 0.28 | 0.14 | 0.32 | 0.57 | 0.57 | 0.32 | 0.72 | 0.46 | 0.46 | 0.62 | 0.69 | 0.76 | -0.56 | 0.56 | -0.20 | -0.51 | -0.63 | -0.31 | -0.29 | -0.20 | -0.36 | -0.21 |
| 5(6)-DHET | 0.08 | 0.06 | 0.11 | 0.09 | 0.20 | 0.23 | 0.26 | 0.06 | -0.03 | 0.03 | -0.02 | 0.13 | 0.14 | -0.02 | 0.02 | 0.09 | -0.19 | -0.16 | 0.07 | -0.05 | 0.04 | -0.04 | 0.02 |
| 8(9)-DHET | 0.10 | 0.05 | 0.12 | 0.12 | 0.04 | 0.05 | 0.24 | 0.22 | 0.11 | 0.07 | -0.11 | -0.11 | -0.21 | 0.02 | -0.02 | 0.01 | 0.14 | 0.11 | 0.01 | 0.14 | 0.05 | 0.00 | -0.02 |
| 11(12)-DHET | 0.15 | 0.16 | 0.21 | 0.23 | 0.19 | 0.25 | 0.34 | 0.25 | 0.10 | 0.07 | -0.01 | -0.05 | 0.02 | 0.13 | -0.13 | 0.08 | -0.02 | 0.01 | -0.06 | -0.07 | -0.13 | -0.18 | -0.16 |
| 14(15)-DHET | 0.20 | 0.16 | 0.12 | 0.27 | 0.18 | 0.28 | 0.31 | 0.31 | 0.25 | 0.16 | 0.07 | 0.05 | -0.01 | -0.02 | 0.02 | 0.03 | -0.11 | -0.03 | -0.11 | 0.01 | -0.01 | -0.18 | -0.02 |
| 9-HODE | 0.17 | 0.20 | 0.15 | 0.17 | 0.13 | 0.11 | 0.10 | 0.04 | 0.03 | -0.13 | 0.08 | -0.03 | 0.06 | -0.02 | 0.02 | -0.08 | -0.11 | -0.35 | -0.02 | -0.30 | -0.17 | -0.10 | -0.15 |
| 13-HODE | 0.18 | 0.23 | 0.12 | 0.13 | 0.20 | 0.16 | 0.22 | 0.20 | 0.18 | -0.08 | 0.16 | 0.01 | 0.09 | -0.12 | 0.12 | -0.17 | -0.22 | -0.40 | -0.07 | -0.40 | -0.24 | -0.11 | -0.12 |
| 9(10)-DiHOME | 0.38 | 0.24 | 0.27 | 0.24 | 0.16 | 0.15 | 0.29 | 0.21 | 0.02 | 0.06 | -0.11 | -0.12 | -0.17 | 0.15 | -0.15 | -0.03 | -0.01 | -0.05 | -0.06 | -0.08 | 0.00 | 0.01 | -0.09 |
| 12(13)-DiHOME | 0.18 | 0.07 | 0.19 | 0.12 | 0.25 | 0.26 | 0.44 | 0.21 | 0.10 | 0.11 | 0.06 | 0.08 | 0.05 | 0.10 | -0.10 | -0.14 | 0.07 | 0.01 | -0.01 | -0.02 | -0.02 | 0.03 | 0.01 |

Color Scale: 0.76 / -0.56

p < 0.05 :

Rho > 0.25 & p < 0.05 :

B

| ρ / Plasma | Percentage of CD4+ non-Naive PD1+ | Percentage of CD8+EM1 HLA DR+CD38+ | Percentage of CD8+CM HLA DR+ CD38+ | Percentage of CD8+CM KI67+ | Percentage of CD8+ RA+ CD27+ R7+ CD95+ KI67+ | Percentage of B cell not PB | Percentage of B cell PB | Percentage of Live CD8+ RA+ CD27+ R7- | Percentage of Live CD8+CM | Percentage of Live CD4+EM1 | Percentage of Live T cell | Percentage of Live CD4+ non-Naive | Percentage of Live CD4+ KI67+ | Percentage of CD4+EM3 KI67+ | Percentage of CD8+ CD38+ | Percentage of CD8+EMRA HLADR+CD38+ | Percentage of CD8+nonNaive CD38+ | Percentage of CD8+nonNaive HLADR+CD38+ | Percentage of CD8+nonNaive CD95+ |
|---|---|---|---|---|---|---|---|---|---|---|---|---|---|---|---|---|---|---|---|
| TxB2 | 0.05 | 0.26 | 0.21 | 0.21 | 0.23 | -0.07 | 0.06 | 0.09 | -0.03 | 0.03 | -0.02 | 0.05 | 0.25 | 0.07 | 0.23 | 0.18 | 0.26 | 0.23 | 0.30 |
| PGE2 | -0.10 | 0.12 | 0.06 | 0.08 | 0.13 | -0.07 | 0.07 | -0.06 | -0.02 | 0.06 | 0.03 | 0.07 | 0.10 | -0.01 | 0.13 | 0.07 | 0.11 | 0.14 | 0.29 |
| iso-iPF2a-VI | -0.03 | 0.16 | 0.10 | 0.08 | 0.00 | -0.10 | 0.10 | -0.06 | -0.05 | -0.02 | -0.04 | 0.07 | 0.07 | 0.04 | 0.16 | 0.11 | 0.14 | 0.13 | 0.08 |
| LTB4 | -0.08 | -0.03 | 0.08 | 0.09 | -0.02 | 0.06 | -0.06 | 0.03 | 0.14 | 0.14 | 0.09 | 0.11 | 0.14 | 0.01 | 0.08 | 0.05 | 0.04 | 0.03 | 0.04 |
| LTE4 | -0.05 | 0.10 | 0.13 | 0.11 | 0.11 | -0.07 | 0.07 | -0.11 | -0.22 | -0.23 | -0.22 | -0.21 | -0.02 | 0.19 | 0.01 | -0.09 | 0.01 | 0.05 | -0.20 |
| 5-HETE | -0.19 | -0.04 | -0.18 | -0.17 | -0.10 | 0.21 | -0.22 | 0.14 | 0.07 | 0.24 | 0.14 | 0.14 | 0.08 | 0.09 | 0.00 | 0.03 | 0.03 | 0.00 | -0.05 |
| 12-HETE | -0.07 | 0.06 | -0.03 | -0.12 | -0.03 | 0.08 | -0.08 | -0.01 | -0.01 | 0.14 | 0.01 | 0.06 | 0.00 | 0.02 | -0.03 | 0.00 | -0.01 | 0.07 | 0.19 |
| 15-HETE | -0.13 | 0.20 | 0.08 | 0.08 | 0.04 | 0.13 | -0.14 | -0.01 | 0.06 | 0.12 | 0.04 | 0.06 | 0.18 | 0.09 | 0.16 | 0.06 | 0.16 | 0.18 | 0.14 |
| 5(6)-DHET | -0.24 | 0.06 | -0.04 | 0.01 | -0.02 | 0.01 | -0.01 | -0.05 | 0.15 | -0.01 | -0.01 | -0.03 | 0.05 | -0.01 | 0.10 | 0.04 | 0.13 | 0.09 | -0.04 |
| 8(9)-DHET | -0.06 | 0.01 | 0.00 | 0.04 | 0.05 | 0.06 | -0.06 | -0.07 | 0.22 | 0.02 | 0.08 | 0.04 | 0.23 | 0.05 | 0.08 | 0.07 | 0.10 | 0.07 | 0.09 |
| 11(12)-DHET | -0.08 | -0.03 | -0.06 | -0.08 | -0.17 | 0.11 | -0.11 | 0.02 | 0.21 | -0.01 | 0.06 | 0.04 | 0.04 | -0.05 | 0.10 | 0.15 | 0.04 | 0.05 | -0.03 |
| 14(15)-DHET | -0.27 | 0.01 | 0.03 | 0.01 | -0.12 | 0.10 | -0.10 | -0.12 | 0.13 | -0.03 | -0.05 | 0.07 | -0.11 | -0.08 | 0.04 | 0.07 | 0.05 | 0.11 | 0.04 |
| 9-HODE | -0.15 | -0.05 | -0.04 | -0.03 | -0.12 | 0.08 | -0.08 | 0.02 | -0.02 | 0.23 | 0.17 | 0.22 | 0.05 | -0.02 | 0.03 | 0.03 | 0.01 | 0.00 | -0.07 |
| 13-HODE | -0.22 | -0.06 | -0.05 | -0.05 | -0.09 | 0.06 | -0.06 | -0.06 | -0.06 | 0.17 | 0.08 | 0.13 | -0.01 | -0.03 | -0.02 | -0.03 | -0.04 | -0.02 | -0.07 |
| 10)-DiHOME | -0.06 | -0.13 | -0.11 | -0.09 | -0.07 | -0.05 | 0.05 | -0.01 | -0.02 | 0.06 | 0.20 | 0.07 | 0.03 | -0.04 | -0.09 | -0.13 | -0.10 | -0.15 | -0.18 |
| 13)-DiHOME | -0.09 | 0.02 | 0.20 | 0.13 | -0.07 | -0.06 | 0.06 | -0.22 | -0.13 | -0.18 | -0.05 | -0.09 | -0.15 | -0.04 | 0.05 | 0.07 | 0.02 | 0.01 | -0.10 |
| AA | -0.19 | -0.08 | -0.12 | -0.07 | -0.10 | 0.05 | -0.05 | 0.01 | 0.01 | 0.18 | 0.14 | 0.14 | 0.09 | 0.06 | -0.02 | -0.06 | -0.02 | -0.05 | -0.04 |
| LA | -0.25 | -0.06 | -0.02 | 0.01 | -0.08 | 0.02 | -0.02 | -0.08 | 0.00 | 0.11 | 0.05 | 0.08 | 0.00 | 0.00 | -0.03 | -0.14 | -0.01 | -0.03 | -0.16 |
| 2-AG | 0.24 | 0.17 | -0.05 | -0.24 | -0.67 | 0.12 | -0.12 | 0.07 | -0.19 | -0.31 | -0.36 | -0.10 | 0.00 | 0.36 | 0.10 | 0.48 | 0.26 | 0.29 | 0.10 |
| AEA | 0.24 | 0.52 | 0.02 | 0.19 | -0.19 | 0.00 | 0.00 | 0.38 | 0.43 | 0.55 | 0.41 | 0.52 | 0.86 | 0.74 | 0.81 | 0.86 | 0.88 | 0.79 | 0.76 |

Color Scale: 0.88 / -0.67

p < 0.05 :

Rho > 0.25 & p < 0.05 :

Figure 3

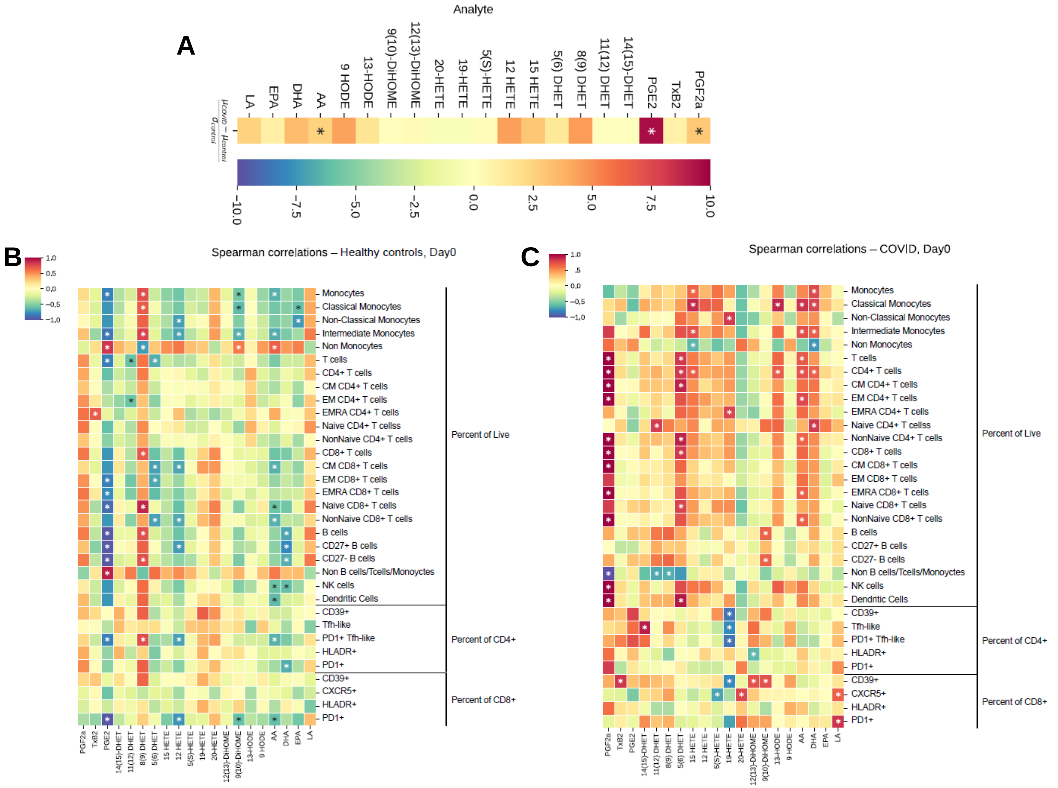
A
Analyte
LA
EPA
DHA
AA
9 HODE
13-HODE
9(10)-DiHOME
12(13)-DiHOME
20-HETE
19-HETE
5(S)-HETE
12 HETE
15 HETE
5(6) DHET
8(9) DHET
11(12) DHET
14(15)-DHET
PGE2
TxB2
PGF2a
B
Spearman correlations – Healthy controls, Day0
C
Spearman correlations – COVID, Day0
Monocytes
Classical Monocytes
Non-Classical Monocytes
Intermediate Monocytes
Non Monocytes
T cells
CD4+ T cells
CM CD4+ T cells
EM CD4+ T cells
EMRA CD4+ T cells
Naive CD4+ T cellss
NonNaive CD4+ T cells
CD8+ T cells
CM CD8+ T cells
EM CD8+ T cells
EMRA CD8+ T cells
Naive CD8+ T cells
NonNaive CD8+ T cells
B cells
CD27+ B cells
CD27- B cells
Non B cells/Tcells/Monoyctes
NK cells
Dendritic Cells
CD39+
Tfh-like
PD1+ Tfh-like
HLADR+
PD1+
CD39+
CXCR5+
HLADR+
PD1+
Percent of Live
Percent of CD4+
Percent of CD8+

Figure 4

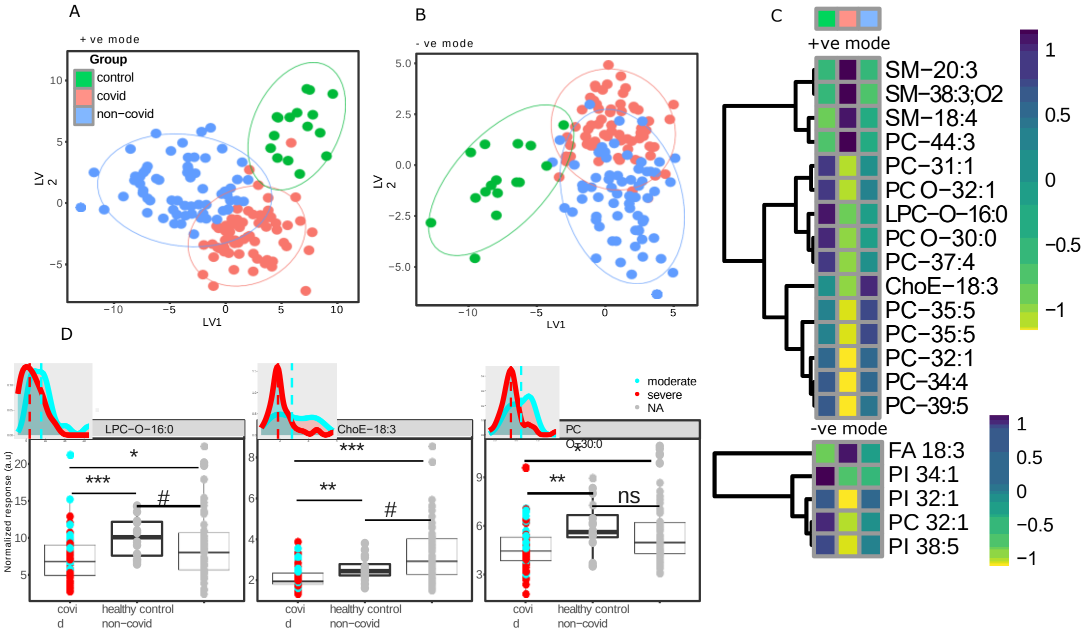
A
+ ve mode
Group
control
covid
non−covid
LV2
LV1
B
- ve mode
LV2
LV1
C
+ve mode
SM−20:3
SM−38:3;O2
SM−18:4
PC−44:3
PC−31:1
PC O−32:1
LPC−O−16:0
PC O−30:0
PC−37:4
ChoE−18:3
PC−35:5
PC−35:5
PC−32:1
PC−34:4
PC−39:5
-ve mode
FA 18:3
PI 34:1
PI 32:1
PC 32:1
PI 38:5
D
moderate
severe
NA
LPC−O−16:0
ChoE−18:3
PC O-30:0
Normalized response (a.u)
covid
healthy control
non−covid
ns

Figure 5

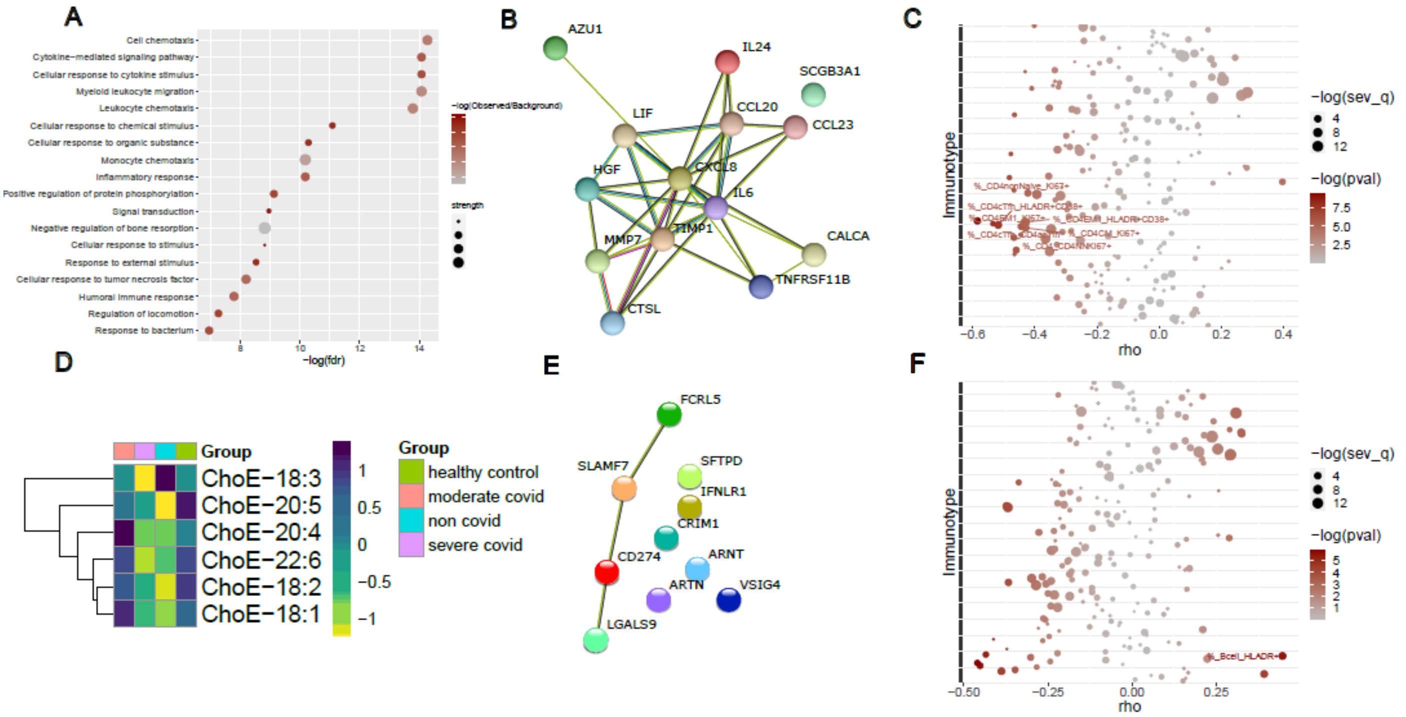
A
Cell chemotaxis
Cytokine−mediated signaling pathway
Cellular response to cytokine stimulus
Myeloid leukocyte migration
Leukocyte chemotaxis
Cellular response to chemical stimulus
Cellular response to organic substance
Monocyte chemotaxis
Inflammatory response
Positive regulation of protein phosphorylation
Signal transduction
Negative regulation of bone resorption
Cellular response to stimulus
Response to external stimulus
Cellular response to tumor necrosis factor
Humoral immune response
Regulation of locomotion
Response to bacterium
8
10
12
14
−log(fdr)
−log(Observed/Background)
strength
B
AZU1
IL24
SCGB3A1
LIF
CCL20
CCL23
HGF
CXCL8
IL6
MMP7
TIMP1
CALCA
TNFRSF11B
CTSL
C
Immunotype
%_CD4nonNaive_KI67+
%_CD4cTfh_HLADR+CD38+
%_CD4CM_KI67+
%_CD4CM_KI67+
−0.6
−0.4
−0.2
0.0
0.2
0.4
rho
−log(sev_q)
4
8
12
−log(pval)
7.5
5.0
2.5
D
Group
ChoE−18:3
ChoE−20:5
ChoE−20:4
ChoE−22:6
ChoE−18:2
ChoE−18:1
1
0.5
0
−0.5
−1
Group
healthy control
moderate covid
non covid
severe covid
E
FCRL5
SLAMF7
SFTPD
IFNLR1
CRIM1
CD274
ARNT
ARTN
VSIG4
LGALS9
F
Immunotype
%_Bcell_HLADR+
−0.50
−0.25
0.00
0.25
rho
−log(sev_q)
4
8
12
−log(pval)
5
4
3
2
1

Figure 6

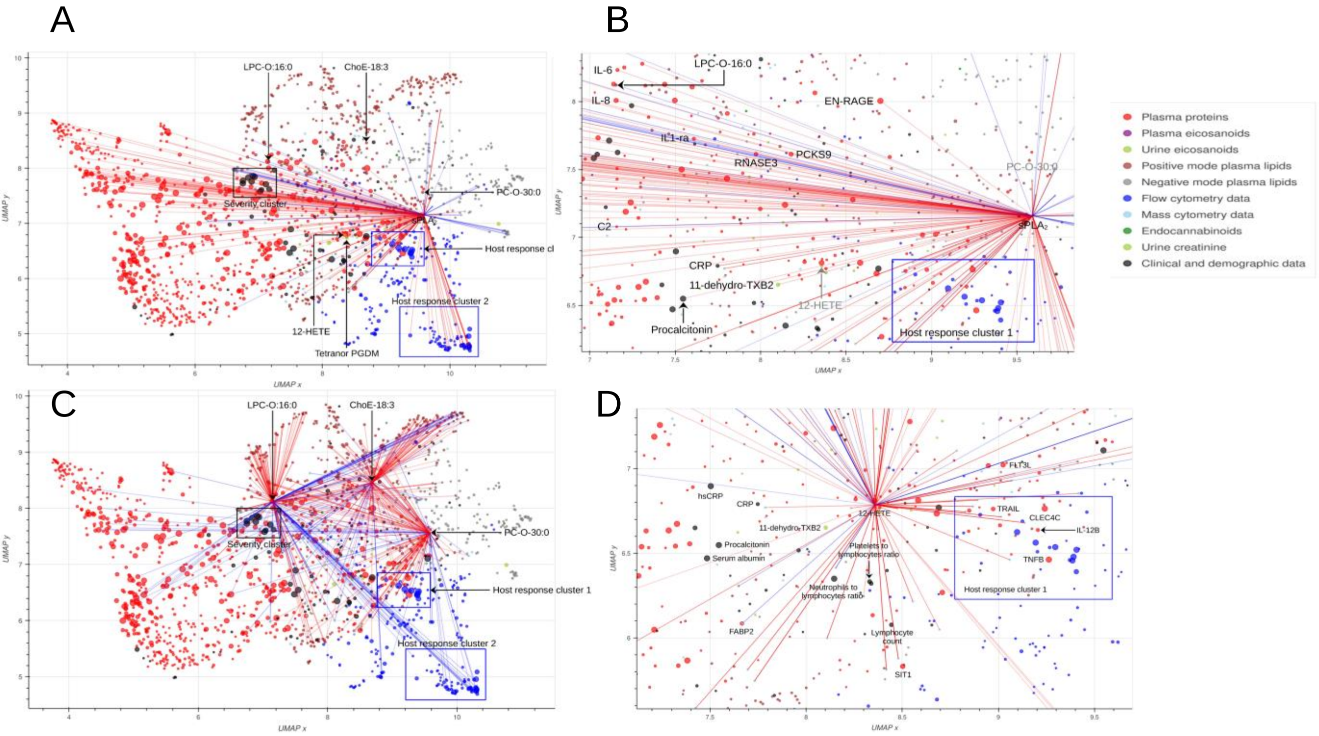
A
B
C
D
LPC-O:16:0
ChoE-18:3
PC-O-30:0
Severity cluster
Host response cluster 1
Host response cluster 2
12-HETE
Tetranor PGDM
IL-6
IL-8
LPC-O-16:0
EN-RAGE
IL1-ra
PCKS9
RNASE3
C2
CRP
11-dehydro-TXB2
Procalcitonin
hsCRP
Serum albumin
FLT3L
TRAIL
CLEC4C
IL-12B
TNFB
FABP2
SIT1
Lymphocyte count
Platelets to lymphocytes ratio
Neutrophils to lymphocytes ratio
UMAP x
UMAP y
Plasma proteins
Plasma eicosanoids
Urine eicosanoids
Positive mode plasma lipids
Negative mode plasma lipids
Flow cytometry data
Mass cytometry data
Endocannabinoids
Urine creatinine
Clinical and demographic data

Figure 7
