## Supplementary material for "Deep Phenotyping of the Lipidomic Response in COVID and non-COVID Sepsis": All Supplementary Tables

| | | | | |
|---|---|---|---|---|
| | mitochondrial (DECR1) | 03 | 6 | |
| | CVD2_Q99523 Sortilin (SORT1) | 0.289886233 | 0.018226222 | 66 |
| | CVD2_Q9BQR3 Serine protease 27 (PRSS27) | -0.374094562 | 0.001972916 | 66 |
| | CVD2_Q9BUD6 Spondin-2 (SPON2) | 0.275065233 | 0.025400751 | 66 |
| | CVD2_Q9BWV1 Brother of CDO (Protein BOC) | -0.28713078 | 0.019410723 | 66 |
| | CVD2_Q9UJM8 Hydroxyacid oxidase 1 (HAOX1) | 0.349754723 | 0.003993825 | 66 |
| | CVD2_Q9UKP3 Melusin (ITGB1BP2) | 0.42578019 | 0.000364957 | 66 |
| | CVD2_Q9Y6K9 NF-kappa-B essential modulator (NEMO) | 0.370504123 | 0.002196722 | 66 |
| | CVD3_O00300 Osteoprotegerin (OPG) | 0.283435315 | 0.02214175 | 65 |
| | CVD3_P00749 Urokinase-type plasminogen activator (uPA) | 0.403715035 | 0.000853412 | 65 |
| | CVD3_P01130 Low-density lipoprotein receptor (LDL receptor) | 0.289554196 | 0.019308338 | 65 |
| | CVD3_P04275 von Willebrand factor (vWF) | 0.257124126 | 0.038669934 | 65 |
| | CVD3_P05121 Plasminogen activator inhibitor 1 (PAI) | 0.382036713 | 0.001687415 | 65 |
| | CVD3_P05164 Myeloperoxidase (MPO) | 0.377272727 | 0.001948293 | 65 |
| | CVD3_P10451 Osteopontin (OPN) | 0.36486014 | 0.002805602 | 65 |
| | CVD3_P10646 Tissue factor pathway inhibitor (TFPI) | 0.261363636 | 0.035465751 | 65 |
| | CVD3_P13500 Monocyte chemotactic protein 1 | 0.4062062 | 0.000 | 65 |

| | | | | |
|---|---|---|---|---|
| | | | 5 | |
| | INF_P09341 C-X-C motif chemokine 1 (CXCL1) | 0.447740319 | 0.000163537 | 66 |
| | INF_P09603 Macrophage colony-stimulating factor 1 (CSF-1) | 0.429078384 | 0.000324622 | 66 |
| | INF_P10145 Interleukin-8 (IL-8) | 0.465275023 | 8.27499E-05 | 66 |
| | INF_P13232 Interleukin-7 (IL-7) | 0.402567582 | 0.000805265 | 66 |
| | INF_P13500 Monocyte chemotactic protein 1 (MCP-1) | 0.442271162 | 0.000200753 | 66 |
| | INF_P13725 Oncostatin-M (OSM) | 0.291597954 | 0.017521957 | 66 |
| | INF_P14210 Hepatocyte growth factor (HGF) | 0.323160422 | 0.008129881 | 66 |
| | INF_P15018 Leukemia inhibitory factor (LIF) | 0.276985701 | 0.024354056 | 66 |
| | INF_P21583 Stem cell factor (SCF) | -0.272393278 | 0.026920046 | 66 |
| | INF_P22301 Interleukin-10 (IL10) | 0.474585116 | 5.67598E-05 | 66 |
| | INF_P42830 C-X-C motif chemokine 5 (CXCL5 ) | 0.247761194 | 0.04488244 | 66 |
| | INF_P50225 Sulfotransferase 1A1 (ST1A1) | 0.284876318 | 0.020428115 | 66 |
| | INF_P55773 C-C motif chemokine 23 (CCL23) | 0.352468427 | 0.003701635 | 66 |
| | INF_P78556 C-C motif chemokine 20 (CCL20) | 0.345872038 | 0.004447407 | 66 |
| | INF_P80075 Monocyte chemotactic protein 2 (MCP-2) | 0.441853669 | 0.000203891 | 66 |
| | INF_P80098 Monocyte chemotactic protein 3 | 0.5497338 | 1.746 | 66 |

| | | | | |
|---|---|---|---|---|
| | | | 8 | |
| | IRE_Q04637 Eukaryotic translation initiation factor 4 gamma 1 (EIF4G1) | 0.413088404 | 0.000566519 | 66 |
| | IRE_Q05084 Islet cell autoantigen 1 (ICA1) | 0.423024736 | 0.000402107 | 66 |
| | IRE_Q05516 Zinc finger and BTB domain-containing protein 16 (ZBTB16) | 0.398726646 | 0.000912993 | 66 |
| | IRE_Q06830 Peroxiredoxin-1 (PRDX1) | 0.339776641 | 0.005251649 | 66 |
| | IRE_Q07065 Cytoskeleton-associated protein 4 (CKAP4) | 0.382068678 | 0.001547295 | 66 |
| | IRE_Q12968 Nuclear factor of activated T-cells, cytoplasmic 3 (NFATC3) | 0.301868281 | 0.013766755 | 66 |
| | IRE_Q13490 Baculoviral IAP repeat-containing protein 2 (BIRC2) | 0.421104269 | 0.000430004 | 66 |
| | IRE_Q14203 Dynactin subunit 1 (DCTN1) | 0.342573844 | 0.004867902 | 66 |
| | IRE_Q8IU57 Interferon lambda receptor 1 (IFNLR1) | 0.395261455 | 0.001021193 | 66 |
| | IRE_Q8NHJ6 Leukocyte immunoglobulin-like receptor subfamily B member 4 (LILRB4) | 0.418307066 | 0.0004738 | 66 |
| | IRE_Q96DB9 FXYD domain-containing ion transport regulator 5 (FXYD5) | 0.309717149 | 0.011385523 | 66 |
| | IRE_Q96SB3 Neurabin-2 (PPP1R9B) | 0.33927565 | 0.005323128 | 66 |
| | IRE_Q9C035 Tripartite motif-containing protein 5 (TRIM5) | 0.510406012 | 1.19659E-05 | 66 |
| | IRE_Q9GZT9 Egl nine homolog 1 (EGLN1) | 0.252520614 | 0.04079862 | 66 |
| | IRE_Q9HCM2 Plexin-A4 (PLXNA4) | 0.274146749 | 0.025914625 | 66 |
| | IRE_Q9NWZ3 Interleukin-1 receptor-associated kinase 4 (IRAK4) | 0.26813485 | 0.02949943 | 66 |

| | | | | |
|---|---|---|---|---|
| | | | 5 | |
| | IRE_Q9UKX5 Integrin alpha-11 (ITGA11) | -0.251059388 | 0.042018348 | 66 |
| | IRE_Q9UQQ2 SH2B adapter protein 3 (SH2B3) | 0.288174512 | 0.018954543 | 66 |
| | IRE_Q9UQV4 Lysosome-associated membrane glycoprotein 3 (LAMP3) | 0.333889991 | 0.006147413 | 66 |
| | IRE_Q9Y2J8 Protein-arginine deiminase type-2 (PADI2) | 0.275440977 | 0.025193024 | 66 |
| | IRE_Q9Y3P8 Signaling threshold-regulating transmembrane adapter 1 (SIT1) | -0.364032982 | 0.002658081 | 66 |
| | MET_O00161 Synaptosomal-associated protein 23 (SNAP23) | 0.269512577 | 0.028643127 | 66 |
| | MET_O15123 Angiopoietin-2 (ANGPT2) | 0.399185889 | 0.00089946 | 66 |
| | MET_O75356 Ectonucleoside triphosphate diphosphohydrolase 5 (ENTPD5) | 0.383362906 | 0.001486604 | 66 |
| | MET_O75791 GRB2-related adapter protein 2 (GRAP2) | 0.36950214 | 0.002263106 | 66 |
| | MET_O95544 NAD kinase (NADK) | 0.490700344 | 2.8788E-05 | 66 |
| | MET_O95841 Angiopoietin-related protein 1 (ANGPTL1) | 0.344744807 | 0.004587358 | 66 |
| | MET_P09104 Gamma-enolase (ENO2) | 0.294102912 | 0.016533367 | 66 |
| | MET_P09417 Dihydropteridine reductase (QDPR) | 0.45717566 | 0.000113871 | 66 |
| | MET_P09668 Pro-cathepsin H (CTSH) | 0.274522492 | 0.025703353 | 66 |
| | MET_P12724 Eosinophil cationic protein (RNASE3) | 0.416804091 | 0.000498974 | 66 |
| | MET_P16083 Ribosyldihydronicotinamide | 0.2589917 | 0.03574371 | 66 |

| | | | | |
|---|---|---|---|---|
| | dehydrogenase [quinone] (NQO2) | 55 | 9 | |
| | MET_P19022 Cadherin-2 (CDH2) | 0.358522075 | 0.003117168 | 66 |
| | MET_P19971 Thymidine phosphorylase (TYMP) | 0.46791958 | 8.48205E-05 | 65 |
| | MET_P21964 Catechol O-methyltransferase (COMT) | 0.445068364 | 0.000180845 | 66 |
| | MET_P23526 Adenosylhomocysteinase (AHCY) | 0.292182444 | 0.017286872 | 66 |
| | MET_P27695 DNA-(apurinic or apyrimidinic site) lyase (APEX1) | 0.437762238 | 0.000237112 | 66 |
| | MET_P31431 Syndecan-4 (SDC4) | 0.275399228 | 0.025216034 | 66 |
| | MET_P35754 Glutaredoxin-1 (GLRX) | 0.352677174 | 0.003679965 | 66 |
| | MET_P41236 Protein phosphatase inhibitor 2 (PPP1R2) | 0.455046446 | 0.000123677 | 66 |
| | MET_P43234 Cathepsin O (CTSO) | 0.427324914 | 0.000345528 | 66 |
| | MET_P46109 Crk-like protein (CRKL) | 0.42757541 | 0.000342468 | 66 |
| | MET_P50452 Serpin B8 (SERPINB8) | 0.408955224 | 0.000651339 | 66 |
| | MET_P51693 Amyloid-like protein 1 (APLP1) | -0.326124622 | 0.007532966 | 66 |
| | MET_P52888 Thimet oligopeptidase (THOP1) | 0.322575932 | 0.008252342 | 66 |
| | MET_P98082 Disabled homolog 2 (DAB2) | 0.428619142 | 0.000329982 | 66 |
| | MET_Q02790 Peptidyl-prolyl cis-trans isomerase FKBP4 (FKBP4) | 0.417806075 | 0.00048206 | 66 |
| | MET_Q13275 Semaphorin-3F (SEMA3F) | 0.3038304 | 0.013 | 66 |

| | | | | | |
|---|---|---|---|---|---|
| | | | 062 1 | | |
| | percOf_CD4cTfh_KI67pos | -0.53 | 0.00 037 9 | 40 | 0.301400751 |
| | percOf_CD4EM1_HLADRposCD38pos | -0.43 | 0.00 552 3 | 40 | 0.000783423 |
| | percOf_CD4EM1_KI67pos | -0.58 | 7.40 E- 05 | 40 | 0.026755803 |
| | percOf_CD4nonNaive_KI67pos | -0.42 | 0.00 667 2 | 40 | 0.031992847 |
| | percOf_CD4nonNaive_TBETpos | -0.47 | 0.00 238 8 | 40 | 0.544146679 |
| | percOf_CD8_CD8EM1 | 0.4 | 0.01 122 5 | 40 | 0.087303203 |
| | percOf_CD8_CD8EMRA | -0.48 | 0.00 165 3 | 40 | 0.384334434 |
| | percOf_CD8_TBETpos | -0.48 | 0.00 186 2 | 40 | 0.621801498 |
| | percOf_CD8nonNaive_TBETpos | -0.47 | 0.00 247 2 | 40 | 0.535187498 |
| | percOf_Live_CD4EMRA | -0.41 | 0.00 863 2 | 40 | 0.532803297 |
| | percOf_Live_CD8EMRA | -0.43 | 0.00 547 8 | 40 | 0.906272113 |
| | umap_component2 | -0.4 | 0.01 127 | 39 | 0.513113445 |
| Proteins | CAM_P01033 Metalloproteinase inhibitor 1 (TIMP1) | -0.4 | 0.00 069 7 | 67 | 0.000118445 |
| | CVD2_P05231 Interleukin-6 (IL6) | -0.44 | 0.00 017 2 | 67 | 0.00810288 |
| | CVD2_P07711 Cathepsin L1 (CTSL1) | -0.45 | 0.00 014 5 | 67 | 8.22E-07 |
| | CVD2_P09237 Matrix metalloproteinase-7 | -0.44 | 0.00 | 67 | 0.032766248 |

| | | | | | |
|---|---|---|---|---|---|
| | (MMP-7) | | 0234 | | |
| | CVD3_O00300 Osteoprotegerin (OPG) | -0.41 | 0.000499 | 67 | 0.000215128 |
| | CVD3_P20160 Azurocidin (AZU1 | -0.42 | 0.000476 | 67 | 7.12E-06 |
| | DEV_Q96QR1 Secretoglobin family 3A member 1 (SCGB3A1) | 0.4 | 0.000679 | 67 | 0.000262582 |
| | INF_O00300 Osteoprotegerin (OPG) | -0.43 | 0.000284 | 67 | 0.000558337 |
| | INF_P05231 Interleukin-6 (IL6) | -0.44 | 0.000204 | 67 | 0.007025703 |
| | INF_P10145 Interleukin-8 (IL-8) | -0.42 | 0.000372 | 67 | 0.00036114 |
| | INF_P14210 Hepatocyte growth factor (HGF) | -0.42 | 0.000421 | 67 | 7.29E-07 |
| | INF_P15018 Leukemia inhibitory factor (LIF) | -0.57 | 3.74E-07 | 67 | 0.000706887 |
| | INF_P55773 C-C motif chemokine 23 (CCL23) | -0.49 | 2.58E-05 | 67 | 7.12E-06 |
| | INF_P78556 C-C motif chemokine 20 (CCL20) | -0.51 | 9.28E-06 | 67 | 0.000218196 |
| | INF_Q13007 Interleukin-24 (IL-24) | -0.42 | 0.000345 | 67 | 7.42E-05 |
| | IRE_P05231 Interleukin-6 (IL6) | -0.43 | 0.000301 | 67 | 0.00956383 |
| | ODA_P01258 Calcitonin (CALCA) | -0.41 | 0.000491 | 67 | 7.10E-06 |
| sPLA2 | P14555 sPLA2 | -0.43 | 0.008225 | 36 | 0.133848614 |
| PC O-30:0 | | | | | |

| | | | | | |
|---|---|---|---|---|---|
| Immunotype | percOf_Bcell_EOMESpos | -0.4 | 0.01035 | 40 | 0.050295441 |
| | percOf_CD4cTfh_CD4acTfh | -0.48 | 0.001567 | 40 | 0.025195859 |
| | percOf_CD4EM1_HLADRposCD38pos | -0.45 | 0.003399 | 40 | 0.000783423 |
| | percOf_CD4EM1_KI67pos | -0.42 | 0.00758 | 40 | 0.026755803 |
| | percOf_CD8EM1_HLADRposCD38pos | -0.43 | 0.005366 | 40 | 0.012293841 |
| | percOf_CD8EM1_KI67pos | -0.43 | 0.005084 | 40 | 0.114591817 |
| | percOf_CD8RAposCD27posR7neg_KI67pos | -0.42 | 0.006808 | 40 | 0.025674422 |
| | percOf_CD8RAposCD27posR7posCD95pos_HLADRposCD38pos | -0.46 | 0.002833 | 40 | 0.037329622 |
| | percOf_CD8RAposCD27posR7posCD95pos_KI67pos | -0.56 | 0.000166 | 40 | 0.031992847 |
| | percOf_Live_CD4EM2 | 0.41 | 0.008612 | 40 | 0.041352462 |
| | percOf_Live_CD4nonNaive | 0.4 | 0.010389 | 40 | 1.48E-07 |
| | umap_component1 | -0.41 | 0.009828 | 39 | 0.000317432 |
| Protein | CAM_Q9BXR6 Complement factor H-related protein 5 (CFHR5) | -0.45 | 0.000119 | 67 | 0.549657525 |
| | CVD3_P01130 Low-density lipoprotein receptor (LDL receptor) | -0.42 | 0.000461 | 67 | 0.000462016 |
| | CVD3_P20160 Azurocidin (AZU1 | -0.4 | 0.000744 | 67 | 7.12E-06 |
| | INF_P13232 Interleukin-7 (IL-7) | -0.48 | 4.24E-05 | 67 | 7.46E-05 |

| | | | | | |
|---|---|---|---|---|---|
| | INF_P14210 Hepatocyte growth factor (HGF) | -0.42 | 0.00040 4 | 67 | 7.29E-07 |
| | INF_P80511 Protein S100-A12 (EN-RAGE ) | -0.48 | 3.55E-05 | 67 | 1.21E-08 |
| | IRE_P78362 SRSF protein kinase 2 (SRPK2) | -0.4 | 0.00083 2 | 67 | 0.000194136 |
| | IRE_Q05516 Zinc finger and BTB domain-containing protein 16 (ZBTB16) | -0.4 | 0.00073 4 | 67 | 2.36E-05 |
| | IRE_Q9GZT9 Egl nine homolog 1 (EGLN1) | -0.41 | 0.00059 | 67 | 0.031265046 |
| | MET_P27695 DNA-(apurinic or apyrimidinic site) lyase (APEX1) | -0.46 | 8.20E-05 | 67 | 0.000162327 |
| | MET_Q02790 Peptidyl-prolyl cis-trans isomerase FKBP4 (FKBP4) | -0.41 | 0.00051 1 | 67 | 0.001680452 |
| | ODA_O60934 Nibrin (NBN) | -0.42 | 0.00041 | 67 | 0.001934367 |
| | ODA_P09769 Tyrosine-protein kinase Fgr (FGR) | -0.47 | 5.65E-05 | 67 | 2.10E-06 |
| | ODA_Q02880 DNA topoisomerase 2-beta (TOP2B) | -0.41 | 0.00060 3 | 67 | 0.000215128 |
| | ODA_Q9HAW4 Claspin (CLSPN) | -0.45 | 0.00015 5 | 67 | 0.001265811 |
| ChoE-18:3 | | | | | |
| Immunotype | percOf_Bcell_CD138pos | -0.45 | 0.00342 9 | 40 | 0.062190734 |
| | percOf_Bcell_CD39pos | -0.46 | 0.00283 3 | 40 | 0.114591817 |
| | percOf_Bcell_HLADRpos | 0.45 | 0.00390 8 | 40 | 0.009618759 |
| | percOf_Bcell_KI67pos | -0.43 | 0.00507 4 | 40 | 0.105547477 |
| | percOf_BcellnotPB_CD138pos | -0.41 | 0.00 | 40 | 0.982727613 |
